# Recessive effects in cancer predisposition exposed by genome-wide and proteome-wide association studies

**DOI:** 10.1101/2020.09.16.299602

**Authors:** Nadav Brandes, Nathan Linial, Michal Linial

## Abstract

The characterization of germline genetic variation affecting cancer risk, known as cancer predisposition, is fundamental to preventive and personalized medicine. Current attempts to detect cancer predisposition genomic regions are typically based on small-scale familial studies or genome-wide association studies (GWAS) over dedicated case-control cohorts. In this study, we utilized the UK Biobank as a large-scale prospective cohort to conduct a comprehensive analysis of cancer predisposition using both GWAS and proteome-wide association study (PWAS), a method that highlights genetic associations mediated by functional alterations to protein-coding genes. We discovered 137 unique genomic loci implicated with cancer risk in the white British population across nine cancer types and pan-cancer. While most of these genomic regions are supported by external evidence, our results highlight novel loci as well. We performed a comparative analysis of cancer predisposition between cancer types, finding that most of the implicated regions are cancer-type specific. We further analyzed the role of recessive genetic effects in cancer predisposition. We found that 30 of the 137 cancer regions were recovered only by a recessive model, highlighting the importance of recessive inheritance outside of familial studies. Finally, we show that many of the cancer associations exert substantial cancer risk in the studied cohort, suggesting their clinical relevance.

## Introduction

Cancer is considered a genetic disease. Notably, that characterization typically refers to somatic mutations in tumors, not to inherited germline genetic variation. Nonetheless, like with any non-Mendelian disease, germline variants are known to contribute to cancer predisposition [1].

The best-known examples of cancer predisposition genes are the tumor suppressors BRCA1 and BRCA2 involved in breast and ovarian cancers [2]. By 2014, following thirty years of research, 114 cancer predisposition genes had been reported, mostly based on families with high prevalence of cancer [3]. As of 2016, the list was expanded to include genes from pediatric cancers [4]. Some of the genes most prevalent in children are also known cancer driver genes from the somatic context (e.g. TP53, APC, BRCA2, NF1, PMS2, RB1 and RUNX1). Recently, a unified list of 152 cancer predisposition genes was refined based on germline variants from The Cancer Genome Atlas (TCGA) across 33 cancer types [5] (see also the discussion in [6]).

Although the heritability of most cancers is considerable [7–9] and despite many years of research leading to the identification of numerous genes, the overall genetic risk explained by the discovered genes remains quite limited. It turned out that BRCA1 and BRCA2 are quite exceptional in their high penetrance, with most cancer predisposition genes showing only mild effects [10–13]. Notably, a gene’s effect size is highly dependent on the cancer-type and population context. For example, BRCA1 and BRCA2 are mostly specific to breast and ovarian cancers and are highly enriched among Ashkenazi Jews [2].

In recent years, the exponential growth in genotyped cohorts and the availability of genome-wide association studies (GWAS) have allowed the discovery of additional genetic loci associated with cancer. As opposed to family studies, GWAS can pick genetic factors with smaller effect sizes, provided that the variants are common enough. Several years of GWAS research into cancer predisposition have implicated numerous cancer predisposition loci (e.g. in breast [14, 15], lung [16], prostate [17], colorectal cancer [18] and melanoma [19]). It is anticipated that by increasing cohort sizes, more genetic associations will be found. Interestingly, the overlaps of significant loci in independent GWAS works studying the same cancer types tend to be surprisingly small, supporting the claim that GWAS efforts in this domain are far from saturated [20, 21]. Benefits from GWAS efforts are also constrained by the lack of interpretability of most implicated loci, which do not appear to affect any functional elements in the genome [22].

An important limitation of GWAS is its assumption of additive genetic effects, which leads to oversight of dominant and recessive effects. This limitation is particularly relevant to recessive inheritance. While some GWAS works have modeled recessive inheritance at the variant level, this solves only a small part of the problem, as recessive effects are likely to occur at the gene level. Specifically, it is anticipated that most recessive effects would manifest through compound heterozygosity, namely different variants affecting the two copies of the same gene [23, 24]. Unfortunately, variant-level approaches like GWAS cannot detect genetic patterns that transcend individual variants. As attested by family studies, recessive inheritance is likely to play a major role in cancer predisposition [25–27], like in other complex traits [28].

Recently, we developed Proteome-Wide Association Study (PWAS) [29], a new gene-based method that addresses many of the shortcomings of GWAS. PWAS detects gene-phenotype associations that are mediated by alterations to protein function. To do so, it considers the proteomic context of genetic variants and their functional effects. As a gene-based method, PWAS aggregates the signal from all variants affecting the same protein-coding gene, and can detect genes with dominant and recessive effects. Unlike GWAS, PWAS associations are supported by concrete functional effects in coding genes, making them more interpretable.

The vast majority of GWAS projects are restricted to specific cancer types, limiting our capacity to draw strong conclusions about the similarities and differences between the genetic signatures of cancer types. Moreover, the collective study of all cancer types as a single phenotype, known as pan-cancer, is prevalent in the somatic world [30], but almost totally absent from germline studies. To perform a comparative GWAS analysis of different cancer types and consider them uniformly, a homogenous cohort is required.

The UK Biobank (UKB) cohort [31, 32] is highly suitable for case-control studies, due to its uniform standards and protocols applied across all samples. The UKB covers ∼500,000 genotyped individuals with open medical records. As it recruited only individuals older than 40 years, the cohort is enriched with cancer cases.

In this work we utilized the UKB cohort to study genetic predisposition to cancer across 10 cancer types, including pan-cancer, thereby providing a systematic view of genetic cancer predisposition in the UKB for the first time. We devised a computational pipeline consisting of both GWAS and PWAS, and compared our results to contemporary clinical knowledge about cancer predisposition. We also compared the recovered associations against catalogues of cancer driver genes from the somatic world. Our analysis focused on two neglected aspects of cancer predisposition research. First, we utilized the uniformity of the UKB cohort to perform an unbiased comparative analysis across cancer types and pan-cancer. Second, we took advantage of the capacity of PWAS to model dominant and recessive effects to systematically study the importance of these heritability modes in cancer predisposition across the human coding genome.

## Results

### Genomic and proteomic analysis of cancer predisposition in the UK Biobank

From the UKB, we derived a study cohort of 274,830 unrelated Caucasian individuals (Fig. 1). Based on their medical records, we determined that 56,634 of those individuals had a history of cancer. In addition to this pan-cancer view, we also considered nine distinct cancer types: melanoma, leukemia, breast, ovarian, prostate, lung, skin, pancreatic and colorectal cancer. We studied each of the ten defined cancer types (the nine specific cancers and pan-cancer) using both GWAS and PWAS independently. With GWAS we found 861 significant variant-cancer associations in 97 unique genomic regions, and with PWAS we found 101 significant gene-cancer associations in 65 unique genomic regions. Altogether, the GWAS and PWAS associations mapped to 137 unique genomic regions, available at Supplementary Table S1. The full GWAS and PWAS results across the ten cancer types and all tested variants and genes are available at Supplementary Table S2 and Supplementary Table S3.

**Fig. 1:**
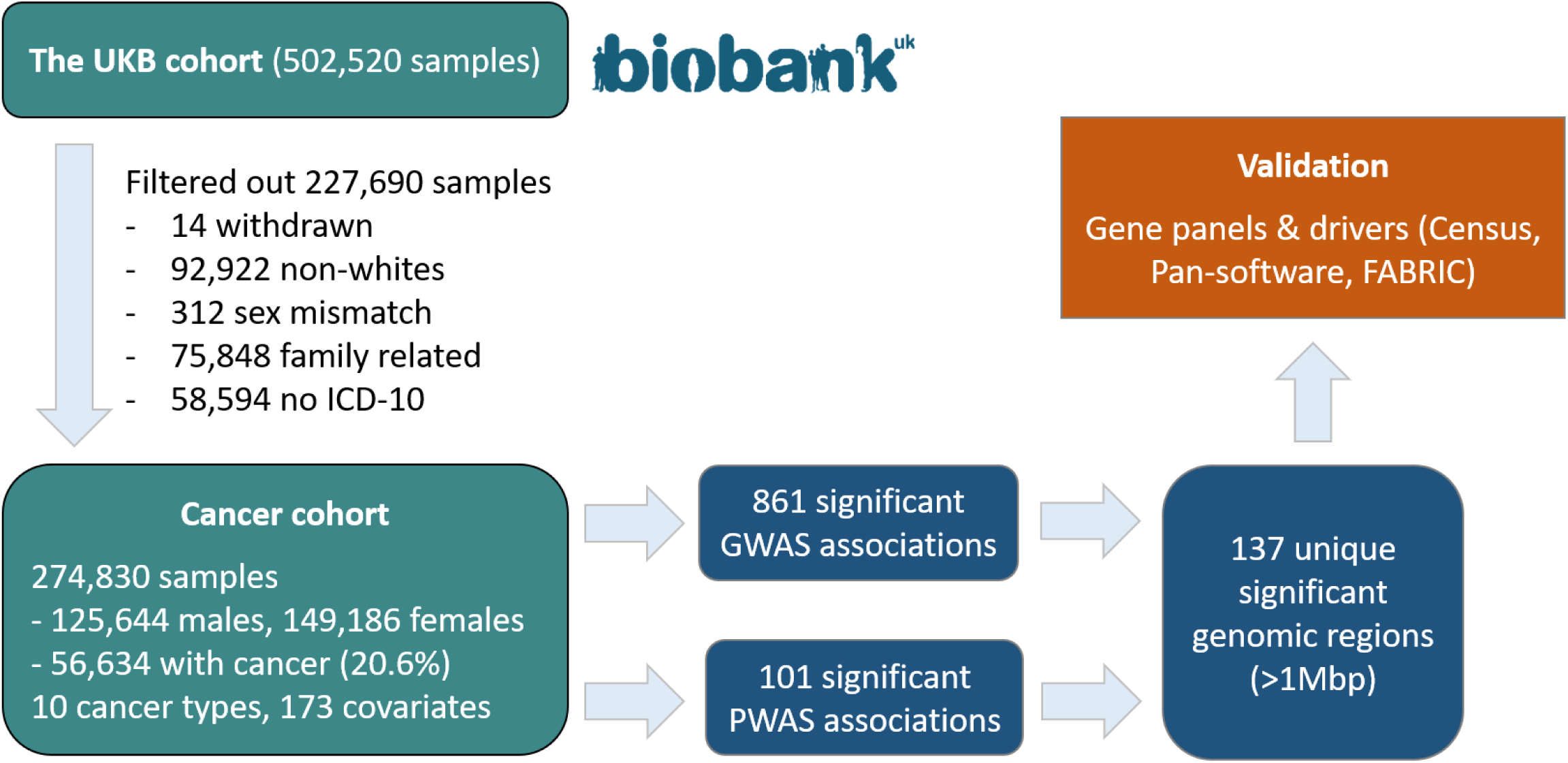
Analysis workflow. From the UK Biobank cohort (∼500K individuals), we derived a study cohort of 274,830 individuals. Based on their medical records, we determined whether each of the individuals had a history of cancer, focusing on nine major cancer types and pan-cancer which combines individuals with any cancer (56,634 overall). We used two statistical methods, standard GWAS and PWAS, over each of the ten cancer types, and found 861 variant-cancer and 101 gene-cancer significant associations, respectively. We merged the significant associations into 137 unique genomic regions (of at least 1,000,000 base pairs each), and validated the regions by comparing them with prominent gene panels used in the clinics and catalogues of cancer driver genes.

We validated our results and compared the discovered genomic regions against prominent cancer predisposition gene panels used in the clinics (CleanPlex TMB 500 and Invitae Multi-Cancer). We also compared our results to three catalogues of cancer driver genes: Census [33], the pan-software catalogue [34] and FABRIC [35]. We find that a majority of the 137 significant cancer regions are indeed supported by external evidence (clinical panels, cancer driver catalogues, or both), but 46 of the 137 regions appear as new discoveries (Fig. 2). We also observe that external validation is slightly stronger for significant regions that are supported by both GWAS and PWAS.

**Fig. 2:**
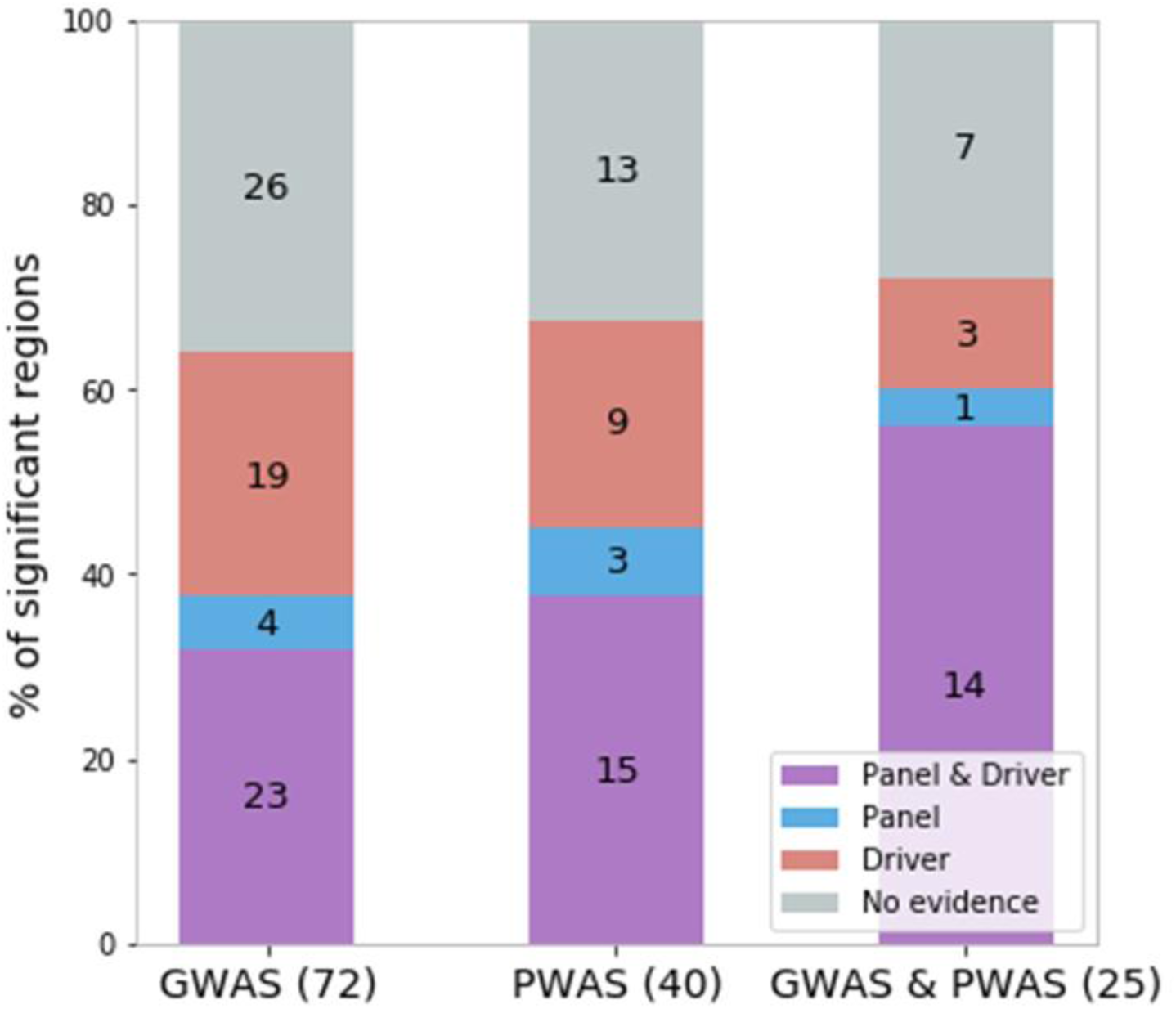
External evidence supporting the cancer predisposition regions. Partitioning of the 137 significant cancer genomic regions according to which method found them (GWAS, PWAS or both) and supporting evidence (overlap with clinical cancer gene panels, driver cancer genes, both, or none).

### Single-gene loci with direct interpretation

To demonstrate the discovered genetic signal of cancer predisposition, we begin by focusing on high-confidence associations with plausible causal mechanisms. One of the benefits of PWAS is its capacity to suggest concrete protein-coding genes underlying genetic associations. However, like GWAS, PWAS is also sensitive to linkage disequilibrium, which can lead to the genetic signal being spread across an extended genomic region and mistakenly attributed to an unrelated nearby gene. For this reason, it is of interest to examine significant associations that are restricted to single genes. We define a significant cancer region to be a soliton region if i) it has exactly one gene significant by PWAS, and ii) the significance is very strong (FDR q-value < 1E-03). Of the 137 cancer regions, 14 are solitons according to this definition (Table 1).

**Table 1:** Soliton cancer genes discovered by PWAS

| Gene | Chromosome bands | Significant cancer types | Lowest PWAS FDR q-value | Lowest GWAS p-value | PWAS heritability | External evidence (out of the 2 gene panels and 3 cancer driver catalogues) |
| --- | --- | --- | --- | --- | --- | --- |
| MUTYH | 1p34.1 | colorectal | $2E-7$ | $1E-3$ | recessive | CleanPlex, Invitae, Census |
| CTLA4 | 2q33.2-q33.3 | skin | $4E-7$ | $1E-12^*$ | both | CleanPlex |
| MITF | 3p14.1-p13 | melanoma | $1E-8$ | $4E-4$ | both | CleanPlex, Invitae, Census |
| SLC45A2 | 5p13.3-p13.2 | skin, melanoma, P-C | $5E-25$ | $3E-26^*$ | recessive | None |
| CCDC170 | 6q25.1-q25.2 | breast | $5E-5$ | $1E-13^*$ | dominant | None |
| POU5F1B | 8q24.21 | prostate, P-C, colorectal, breast | $1E-22$ | $4E-42^*$ | both | None |
| MTAP | 9p21.3 | skin, melanoma, P-C | $1E-4$ | $2E-19^*$ | both | None |
| STN1 | 10q24.33-q25.1 | P-C, melanoma | $9E-6$ | $3E-10^*$ | recessive | None |
| UBE3A | 15q11.2-q12 | pancreatic | $8E-4$ | $3E-4$ | recessive | None |
| HOXB13 | 17q21.32 | prostate, P-C | 2E-31 | 6E-40* | dominant | CleanPlex, Invitae |
| IRF3 | 19q13.33 | skin | 8E-5 | 2E-6 | dominant | None |
| KLK3 | 19q13.33-q13.41 | prostate | 4E-9 | 3E-25* | dominant | None |
| AVP | 20p13 | melanoma | 9E-4 | 2E-4 | recessive | None |
| CHEK2 | 22q12.1-q12.2 | breast, P-C, prostate | 5E-5 | 3E-6 | dominant | CleanPlex, Invitae, Census, PanSoftware, FABRIC |

\*Below the GWAS exome-wide significance threshold (5E-07); P-C, Pan-cancer.

5 of the 14 soliton genes are supported by gene panels (CleanPlex or Invitae), and 3 of them are supported by Census and other cancer driver catalogues. Importantly, the criteria of external evidence for the solitons listed in Table 1 are stricter than those used for the genomic regions in Fig. 2. Whereas in Fig. 2 we considered evidence for any gene within the same region (which may include up to 138 genes; see Supplementary Table S1), in Table 1 we only consider evidence matching the exact soliton gene.

8 of the 14 soliton genes are also found by standard GWAS, whereas the 6 other soliton genes are below the exome-wide significance threshold (5E-07). IRF3 (interferon regulatory factor 3), for example, is associated with skin cancer according to PWAS with overwhelming significance (FDR q-value = 8E-5), but not according to GWAS (p = 2E-6). IRF3 is part of the innate immune system, whose one of its major roles is to recognize and destroy infected and tumorous cells. In-vivo and expression studies have implicated IRF3 in melanoma [36, 37] and other cancer types [38, 39].

The list of solitons also includes novel discoveries. For example, UBE3A (ubiquitin protein ligase E3A; also known as E6AP) is known for resulting in the ubiquitylation and degradation of the p53 tumor suppressor in papillomavirus-positive cervical cancer [40], and it has also been implicated with prostate and other cancers [41–43]. However, E6AP hasn’t been linked to pancreatic cancer, an association that we observe here (PWAS q-value = 8E-04). Another example is AVP (arginine vasopressin; also known as ADH), a gene producing a hormone involved in water balance homeostasis and complex behavioral traits [44]. Despite vast literature about the hormone, there doesn’t appear to be any direct evidence linking the gene to cancer or specifically to melanoma, as observed here (PWAS q-value = 9E-04).

### Predisposition genomic loci are mostly cancer-type specific

The uniformity of the UKB as a prospective cohort that covers all major cancer types provides a unique opportunity to perform a comparative analysis between cancer types. We start by considering the number of significant genomic regions recovered in each cancer type, according to GWAS, PWAS or both (Table 2). We observe that the number of recovered regions vary substantially across cancer types, ranging from 2 in epithelial ovarian cancer to 49 in skin cancer. As statistical power is determined by sample sizes, it is not surprising that cancers with fewer cases tend to recover fewer significant regions. However, this is not the only factor. Chronic lymphocytic leukemia, for example, has the fewest cases (764), yet it recovers 20 significant regions, more than many other cancers. This could be explained by a strong polygenic signal associated with leukemia, or it could be a spurious effect of mistaking somatic mutations as germline variants in blood cancers (as UKB genotypes are based on blood samples). Colorectal cancer includes 3,351 cases but only 4 recovered regions, compared for example to melanoma with 14 recovered regions, despite its slightly lower number of cases (3,122). Interestingly, although pan-cancer has the highest number of cases (by definition), it doesn’t recover the highest number of genomic regions. The major differences in polygenic signal between cancer types are also observed in significance levels (Fig. 3). For example, the 6 significant regions of lung cancer are statistically weaker than the 18 significant regions of breast cancer, although this could simply reflect the different numbers of cases. This cancer-type comparison also reinforces the contribution of both GWAS and PWAS to our analysis, with each of the two methods dominating the discovery in certain cancer types (Table 2).

**Table 2:**
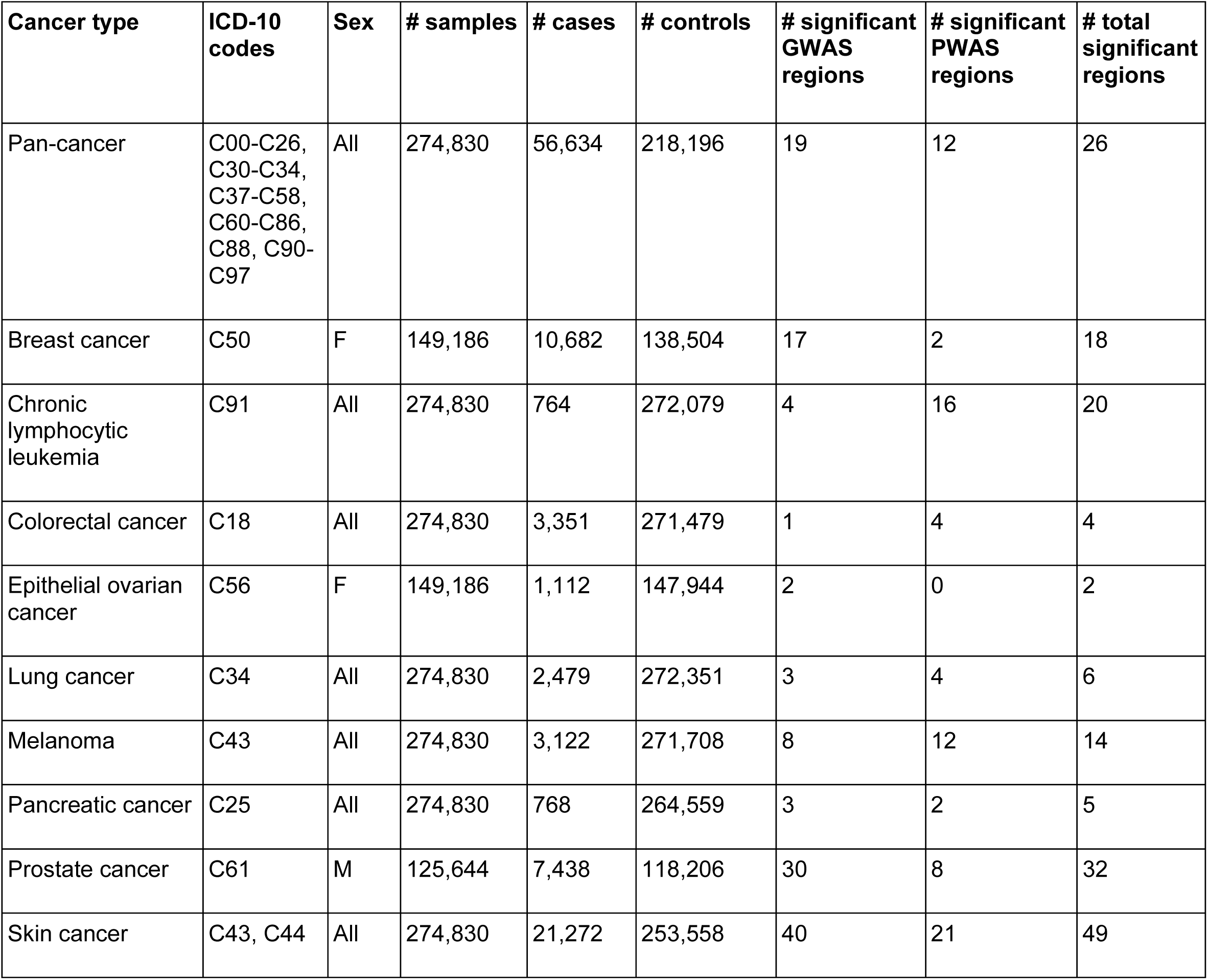
Studied cancer types

**Fig. 3:**
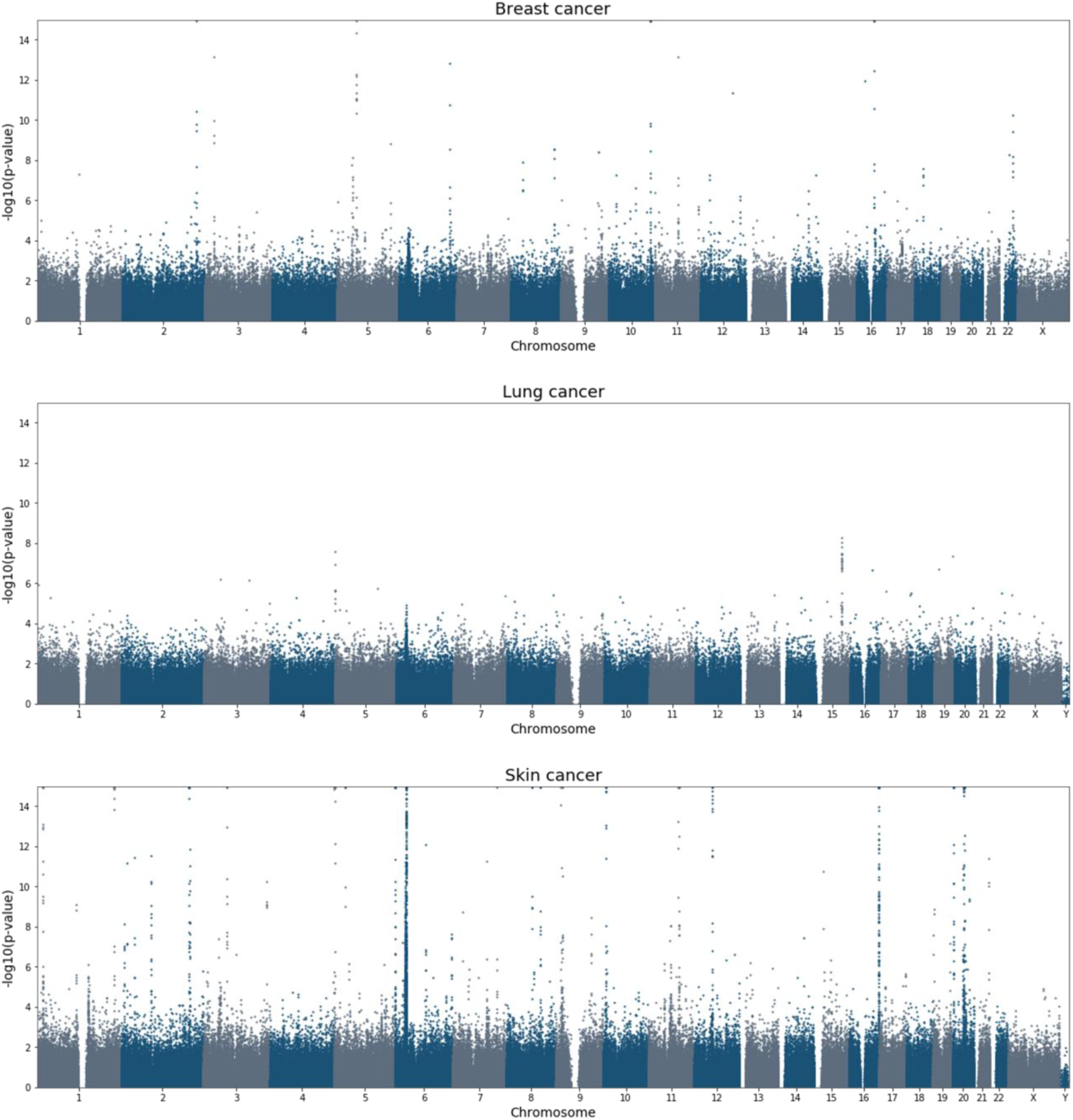
Genome-wide cancer predisposition in breast, lung and skin cancer. Manhattan plots for three selected cancer types. The plots show the significance of all the variants and genes tested with GWAS and PWAS. P-values are capped at 1E-15. Similar Manhattan plots for all ten cancer types (including pan-cancer) are available in Supplementary Fig. S1.

Among the 137 significant cancer predisposition loci, 56 are associated with skin cancer or melanoma. Genes affecting skin pigmentation were implicated with both cancer types. Of all the associations recovered in this study, the most significant is the gene MC1R (Melanocortin 1 receptor; PWAS q-value = 3E-144 in skin cancer), which plays a central role in skin pigmentation, melanin formation and melanocyte differentiation. Many variants in the gene have been associated with pigmentation differences and elevated risk of skin cancer and melanoma [45]. Additional major pigmentation-related genes identified in this study include TYR (tyrosinase; PWAS q-value = 2E-14 in skin cancer), OCA2 (OCA2 melanosomal transmembrane protein; PWAS q-value = 0.05 in skin cancer) and SLC45A2 (solute carrier family 45 member 2; PWAS q-value = 5E-25 in skin cancer) [46].

We also consider the extent to which cancer predisposition genetic signal overlaps between cancers, by counting the number of significant cancer regions shared between cancer types (Fig. 4). Skin cancer and melanoma, for example, share a substantial genetic signal, with 6 of the 14 melanoma regions being also significant in skin cancer. A shared genetic signal between these two cancer types is expected, given their relatedness and the fact that all 3,122 melanoma cases are also among the 21,272 skin-cancer cases (see the ICD-10 definitions of the cancer types in Table 2). More surprising is the fact that 3 of the 18 breast-cancer regions are shared with prostate cancer (with overall 32 regions), while none of the 2 significant ovarian-cancer regions is shared with breast cancer (or any other cancer type). One of the 3 genomic regions shared between breast and prostate cancer is found in 11q13.3. Most significant in this region is the intergenic variant rs7130881 (lowest GWAS p-value = 3E-36). The other 2 regions shared between breast and prostate cancer contain soliton genes listed in Table 1: CHEK2 (22q12.1-q12.2) and POU5F1B (8q24.21). All six associations (the three regions with respect to the two cancer types) have strong support in literature [47–50].

**Fig. 4:**
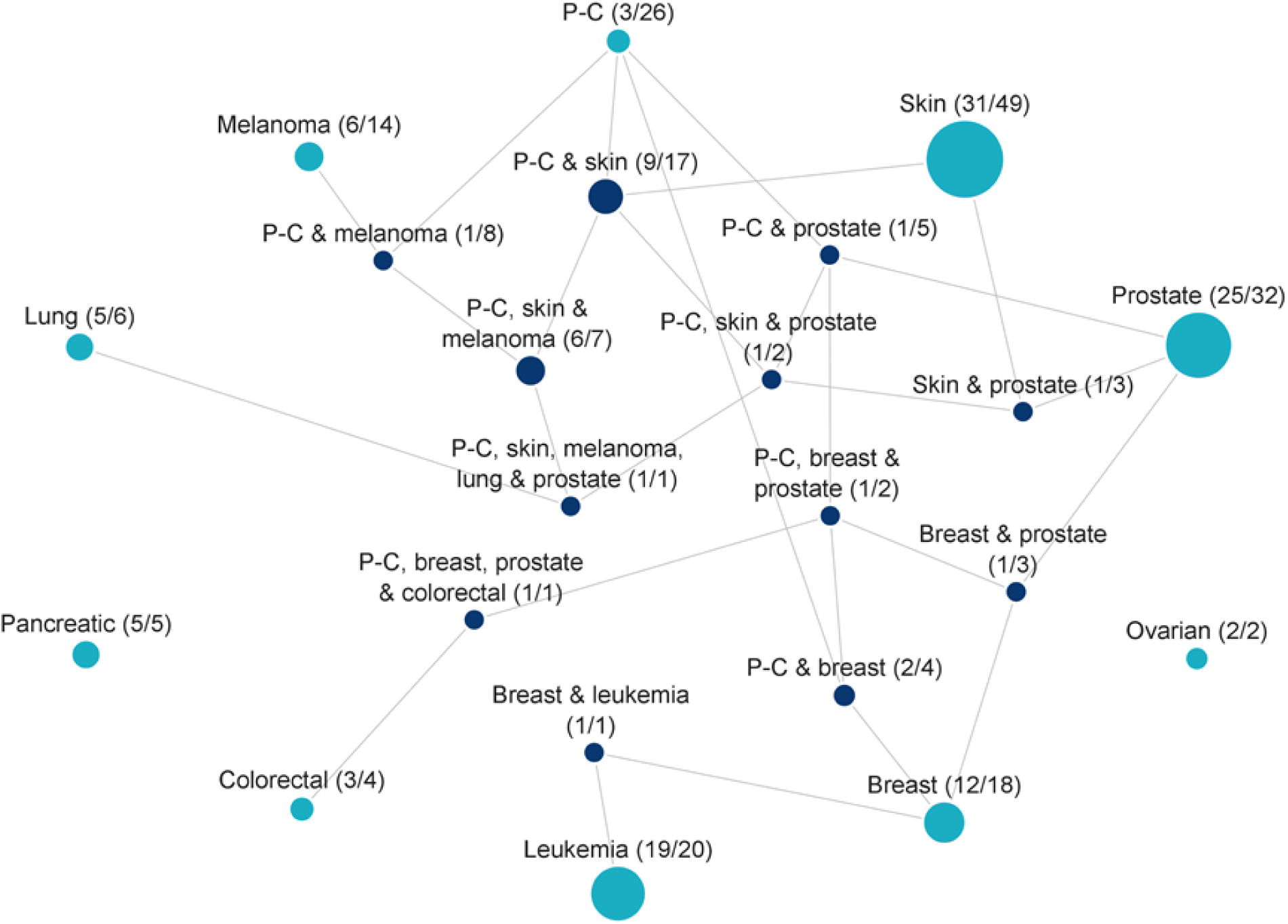
Number of significant regions per cancer-type combination. Partitioning of the 137 significant cancer genomic regions by the combination of cancer types in which they were found significant. Each node in the graph represents a combination of cancer types. Two nodes are connected when one combination is a direct subset of the other. Each node is labeled with two numbers: the number of significant regions matching the exact combination of cancer types represented by the node, and the total number of significant regions matching any combination of cancer types that includes this combination. For example, there are 20 unique genomic regions significantly associated with leukemia, 19 of them are exclusive to leukemia, and 1 also appears in breast cancer. Node sizes reflect the number of significant regions that match the combinations exactly. Nodes representing combinations of more than one cancer type are colored dark blue. Pan-cancer is abbreviated as P-C.

We observe that the pan-cancer phenotype does not substantially add to the recovery of significant genomic regions over the nine specific cancer types. Of the 26 significant pan-cancer regions, only 3 are not recovered by any of the specific cancer types. From a statistical perspective, this stands out given the high number of pan-cancer cases (56,634).

The genomic region with the most dominant pan-cancer signal is 5p15.33, which is also significant in skin cancer, melanoma, prostate cancer and lung cancer. The most significant variant in that region is rs2853677 (lowest GWAS p-value = 8E-20), which occurs in the intronic region of TERT (Telomerase Reverse Transcriptase). TERT is a well-known cancer predisposition gene implicated in numerous cancer types [51]. Mutations in TERT can cause abnormal activities of the telomerase and prevent telomere shortening, thereby leading to cell immortality. rs2853677 has been implicated in various Chinese and Indian populations [52, 53]. We provide evidence for the clinical importance of this variant in the white British population.

Another relatively major genetic overlap is observed between prostate cancer and skin cancer, which share 3 predisposition regions. On top of 5p15.33 (the locus of TERT), prostate and skin cancer also share 6p22.1-p21.31 and 21q22.2-q22.3. The 21q22.2-q22.3 region seems to be a novel discovery of our analysis, without any existing support from the literature. The most significant variant in that region is rs2849691 (lowest GWAS p-value = 4E-12) that occurs in the non-coding lncRNA transcript LOC107985478. The other region, 6p22.1-p21.31, is a 4.7 Mbp genomic region covering the MHC region. Of the 138 genes in that region, 6 are significant in skin cancer according to PWAS: HLA-DPA1, HLA-C, MPIG6B, PPP1R18, CCHCR1 and C6orf15 (Supplementary Table S1).

Other than the handful of shared regions mentioned here, most cancer types show little overlap with other cancers (Fig. 4). Overall, of the 137 cancer predisposition regions, only 26 are shared by multiple cancers. Disregarding pan-cancer, only 13 regions are shared by two or more specific cancer types.

### Recessive effects are prevalent in cancer predisposition

Unlike GWAS, PWAS explicitly models dominant and recessive genetic effects. It therefore provides the opportunity to assess the importance of these effects to cancer predisposition. We observe that recessive effects are indeed common among significant PWAS regions, and especially among the regions that are not captured by GWAS (Fig. 5A). The enrichment of recessive effects in PWAS-exclusive regions is explained by the fact that GWAS models only additive genetic effects. Unlike dominant effects, which can be approximated by an additive model, recessive effects cannot be effectively recovered by GWAS.

**Fig. 5:**
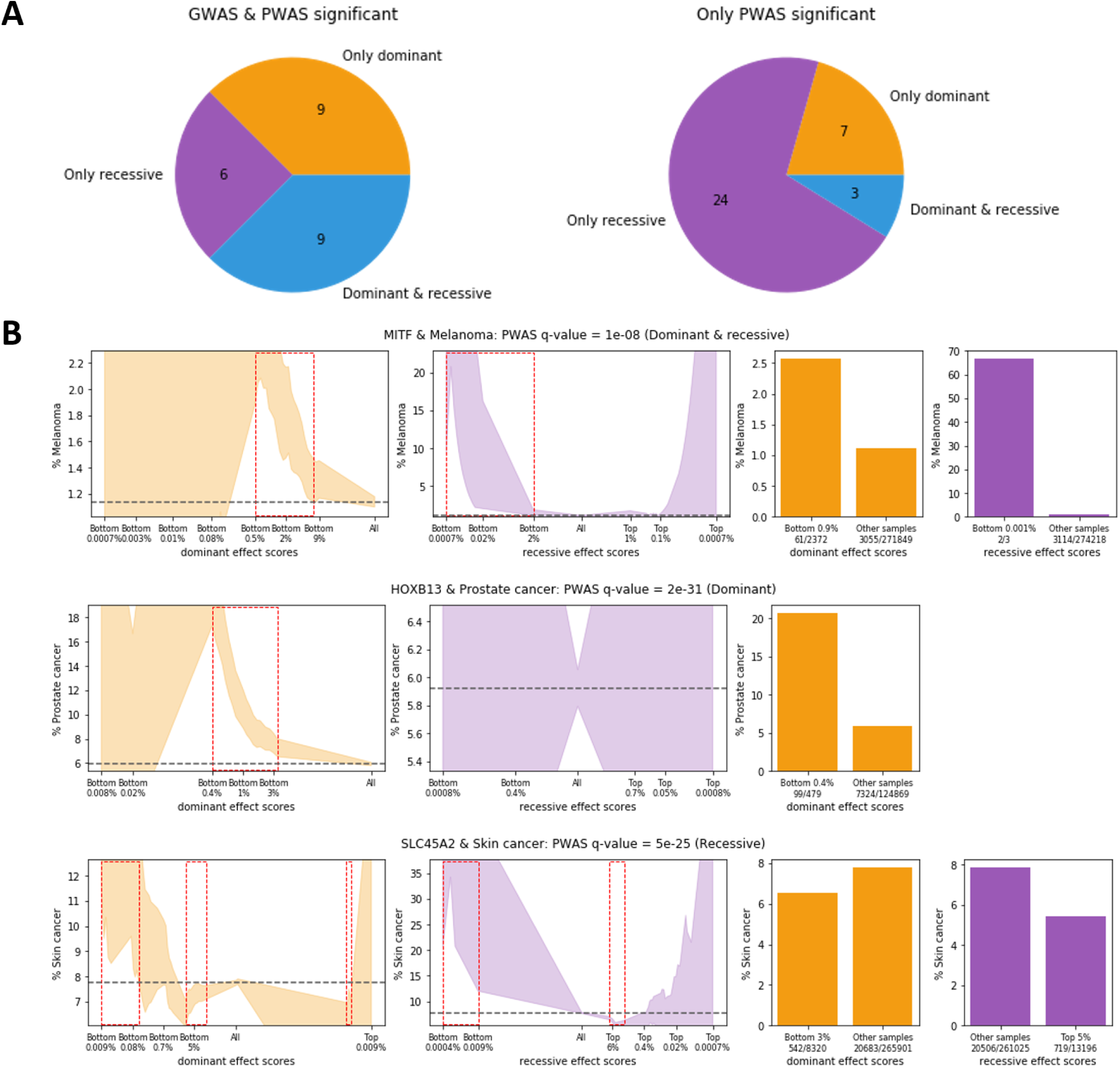
Dominant and recessive gene effects. (**A**) Partitioning of the significant cancer genomic regions that are PWAS-significant according to their inheritance mode (dominant, recessive or both), and whether they are also significant by GWAS. (**B**) Three examples of significant PAWS associations: MITF and melanoma, HOXB13 and prostate cancer, and SLC45A2 and skin cancer. The plots show how risk for the relevant cancer type changes for individuals with varying PWAS effect scores. Dominant effect scores (orange) aim to capture the likelihood of an individual to have at least one damaged copy of the gene, whereas recessive effect scores (purple) aim to capture the likelihood of at least two damaged copies. In both types of PWAS effect scores, lower scores reflect individuals with greater gene damage. The two left columns of plots show a continuous partitioning of the cohort according to the dominant and recessive effect scores. For example, “Bottom 0.5%” on the upper-left plot refers to the 0.5% of the individuals with the lowest dominant effect scores with respect to the MITF gene (i.e. the 0.5% of individuals that are the most likely to have at least one damaged copy of this gene). The color-filled area is stretched between the lower to the higher bound of the 95% confidence interval for the estimated fraction of the selected individuals with a history of the specified cancer type. For example, at least 2% of the individuals with the lowest 0.5% dominant MITF effect scores have a history of melanoma. The dashed horizontal lines show the baseline cancer risks across the entire cohort. The red rectangles indicate partitions whose 95% confidence intervals do not overlap with the baseline rates. The two right columns of bar plots show the most significant partitioning of individuals. For example, among the 0.9% (2,372) of the individuals with the lowest dominant effect scores calculated for the MITF gene, 61 (2.57% of them) have had melanoma, compared to 1.12% rate of melanoma among the other individuals (3,055 melanoma cases among 271,849 individuals). Note that the scales of the plots are not uniform.

To illustrate the dominant and recessive genetic effects captured by PWAS, we selected three soliton associations with strong effects (see Table 1): MITF (melanocyte inducing transcription factor) associated with melanoma (q-value = 1E-08; both dominant and recessive effects are significant), HOXB13 (homeobox B13) associated with prostate cancer (q-value = 2E-31; only dominant effect is significant) and SLC45A2 (solute carrier family 45 member 2) associated with skin cancer (q-value = 5E-25; only recessive effect is significant). For each of the three associations, we plot cancer risk among UKB individuals as a function of PWAS effect scores (Fig. 5B). PWAS effect scores are assigned to each individual in the cohort with respect to each studied protein-coding gene based on their genotype. They reflect the overall estimated functional damage affecting genes at the molecular level, with lower effect scores reflecting more damage. For each individual, each coding gene is assigned two effect scores that aggregate all the variants within the gene that were genotyped or imputed for that individual: a dominant effect score (reflecting the likelihood of at least one damaging hit) and a recessive effect score (reflecting the likelihood of at least two damaging hits). These effect scores are used by PWAS to detect significant genes (if they correlate with a history of cancer), and they can also be used to assess individual-level risk (with respect to specific genes).

In the case of MITF, we observe that among the 2,372 individuals with the lowest dominant effect scores (comprising 0.9% of the cohort), 61 (2.57%) have had melanoma, compared to only 1.12% risk of melanoma in the rest of the cohort (leading to observed risk-ratio of 2.3). When considering recessive effect scores, we find that 2 out of the 3 individuals with the lowest scores have had melanoma. We conclude that individuals with functional damage to the MITF protein have substantially higher rates of melanoma. As the name of the gene suggests, MITF is a transcription factor that controls the development and function of melanocytes, which are pigment-producing cells. The causal connection of MITF to melanoma is well established [54].

Unlike MITF, showing both dominant and recessive effects (which might be interpreted as approximating an additive effect), HOXB13 shows a clear dominant effect according to PWAS. Among the 479 males with the lowest dominant effect scores (0.4% of the male cohort), 99 of them (20.7%) have a history of prostate cancer, compared to 5.87% rate of prostate cancer in the rest of the male cohort (risk-ratio of 3.5). HOXB13 regulates cellular response to androgen, an important male hormone. Its link to prostate cancer and early onset of the disease has been established in both GWAS and family studies [55].

A clear recessive effect is exemplified by SLC45A2 (also known by the names AIM1, MATP and OCA4). Among the 13,196 individuals with highest recessive effect scores, i.e. the 5% of the UKB cohort least likely to have two damaged copies of the gene, 719 of them (5.45%) are skin-cancer cases, compared to 7.86% of cases in the rest of the cohort (risk-ratio of 1.4). SLC45A2 is a transporter protein likely involved in the production of melanin. The gene has indeed been implicated in melanoma and skin cancer through albinism, a disorder known for recessive inheritance [46].

These three examples demonstrate the applicability of PWAS effect scores to clinical risk assessment. Similar plots for all 101 significant PWAS associations are available in Supplementary Fig. S2.

To further examine the recessive associations, we list the 30 genomic regions with only significant recessive effects (Fig. 5A) in Table 3. We find that all 30 recessive regions support exactly one significant association of gene and cancer-type. Notably, 10 of the 30 recessive genes are enzymes (or part of enzyme complexes), which are considered a standard model for recessive effects [56]. Additional 5 genes are transporters associated with the solute carrier family. In fact, of all 101 significant PWAS associations, these genes are the only transporters of that family (namely, all 5 transporters of the solute carrier family that we have recovered are associated with recessive effects). Both SLC45A2 and SLC12A9, which we associate with skin cancer, are implicated with pigmentation according to the Open Targets Platform [57]. We also observe that many of the 30 recessive genes are involved in cellular processes of apoptosis, ubiquitination and immunology. Interestingly, leukemia seems to be particularly enriched with recessive effects. Of the 16 genomic regions associated with leukemia according to PWAS, 12 show exclusive recessive effects and are listed in Table 3. The possibility of mistaking somatic mutations as germline variation in blood cancers should be taken into account when interpreting this finding. For example, the high prevalence of recessive heritability in leukemia might be a reflection of the “second-hit” hypothesis [58]. As an external validation scheme, Table 3 also lists published cancer genetic associations curated by the Open Targets Platform. We find that the cancer types reported by the Open Targets Platform are usually in agreement with our results. Nevertheless, most recessive genes appear to represent novel genetic associations, maybe as a result of recessive genetic studies being largely neglected in case-control cohorts. When considering the external validation used earlier (Fig. 2), only 2 of the 30 recessive genes are supported by clinical panels or catalogues of cancer drivers. Specifically, MUTYH and BIRC3 are both established as cancer drivers, and they both appear in clinical panels. Overall, these 30 recessive genes highlight the importance of modeling recessive inheritance in non-familial (case-control) cancer studies.

**Table 3:** Recessive cancer predisposition genes

| Gene symbol | Gene name | Chromosome bands | Molecular function | Cancer type | Cancer genetic associations according to Open Targets Platform |
| --- | --- | --- | --- | --- | --- |
| PADI1 | Peptidyl Arginine Deiminase 1 | 1p36.13 | enzyme | melanoma | None |
| AKIRIN1 | Akirin 1 | 1p34.3-p34.2 | regulator | lung | None |
| MUTYH | Adenine DNA glycosylase | 1p34.1 | enzyme | colorectal | colorectal and other cancers |
| FCGR1A | High affinity immunoglobulin gamma Fc receptor I | 1q21.2 | receptor | leukemia | None |
| OR6K6 | Olfactory receptor 6K6 | 1q23.1-q23.2 | receptor | pancreatic | None |
| DPT | Dermatopontin | 1q24.2 | ECM | leukemia | leukemia |
| TNFSF18 | Tumor necrosis factor ligand superfamily member 18 | 1q24.3-q25.1 | cytokine | leukemia | None |
| LYZL4 | Lysozyme-like protein 4 | 3p22.1 | enzyme | leukemia | None |
| POLR2H | DNA-directed RNA polymerases I, II, and III subunit RPABC3 | 3q27.1-q27.2 | enzyme | melanoma | None |
| RCHY1 | RING finger and CHY zinc finger domain-containing protein 1 | 4q13.3-q21.1 | enzyme | leukemia | None |
| PCDH10 | Protocadherin-10 | 4q28.3 | cell adhesion | leukemia | None |
| SLC45A2 | Membrane-associated transporter protein | 5p13.3-p13.2 | transporter | skin | skin and melanoma |
| PTPN12 | Tyrosine-protein phosphatase non-receptor type 12 | 7q11.23-q21.11 | enzyme | skin | None |
| LMTK2 | Serine/threonine-protein kinase LMTK2 | 7q21.3-q22.1 | enzyme | prostate | prostate |
| SLC12A9 | Solute carrier family 12 member 9 | 7q22.1 | transporter | melanoma | skin |
| STN1 | CST complex subunit STN1 | 10q24.33-q25.1 | DNA binding | pan-cancer | Many cancers |
| BIRC3 | Baculoviral IAP repeat-containing protein 3 | 11q22.1-q22.2 | regulator | leukemia | leukemia and other blood cancers |
| SLC38A4 | Sodium-coupled neutral amino acid transporter 4 | 12q13.11 | transporter | skin | None |
| UBE3A | Ubiquitin-protein ligase E3A | 15q11.2-q12 | enzyme | pancreatic | leukemia |
| SLC12A6 | Solute carrier family 12 member 6 | 15q14 | transporter | lung | None |
| ATP8B4 | Probable phospholipid-transporting ATPase IM | 15q21.2 | enzyme complex | skin | leukemia |
| STX8 | Syntaxin-8 | 17p13.1 | receptor | melanoma | None |
| ATPAF2 | ATP synthase mitochondrial F1 complex assembly factor 2 | 17p11.2 | enzyme complex | lung | None |
| NLRP9 | NACHT, LRR and PYD domains-containing protein 9 | 19q13.42-q13.43 | regulator | leukemia | None |
| AVP | Vasopressin-neurophysin 2-copeptin | 20p13 | hormone | melanoma | None |
| SLC2A4RG | SLC2A4 regulator | 20q13.33 | regulator | prostate | prostate and other cancers |
| CCT8 | T-complex protein 1 subunit theta | 21q21.3 | chaperone | leukemia | None |
| SGSM3 | Small G protein signaling modulator 3 | 22q13.1-q13.2 | regulator | leukemia | breast |
| RBMX2 | RNA-binding motif protein, X-linked 2 | Xq25-q26.1 | RNA binding | leukemia | None |
| ZNF449 | Zinc finger protein 449 | Xq26.3 | DNA binding | leukemia | None |

## Discussion

We performed a comprehensive analysis of cancer predisposition in the UKB cohort and identified 137 significant genomic regions combining the discoveries of both GWAS and PWAS. The two methods complemented each other. While GWAS identifies variants with additive genetic effects, PWAS identifies coding genes with dominant or recessive effects (or a combination of these effects).

The capacity of PWAS to detect non-additive effects has proven substantial in our analysis. We detected 30 genomic regions with only recessive effects, indicating that recessive inheritance is substantial for cancer predisposition also outside of family studies. Many of the discovered genes with significant recessive effects had not been previously implicated with cancer predisposition (Table 3), indicating that non-additive effects are neglected in contemporary genetic association studies. The main reason for the scarcity of studies on recessive effects is that GWAS and current variant-level methods are not designed to capture such genetic effects, especially when compound heterozygosity is involved. By exposing the importance of recessive inheritance to cancer predisposition and presenting a methodology for recovering it, we aim to motivate future investigation of this underexplored source of cancer risk.

Generally, many of the genomic loci recovered in this study are supported by external evidence (clinical panels and cancer drivers), while many are novel associations (Fig. 2, Table 3). While proving definite causal links is beyond the scope of this work, many of our novel discoveries appear as promising candidates. In particular, genes defined as solitons (Table 1) are likely less sensitive to linkage-disequilibrium and are therefore more likely to prove causal.

Many of the reported genes are associated with substantial cancer risk (Fig. 5B). For example, we show that individuals in the male cohort with substantial damage to HOXB13 have an increased risk of prostate cancer (20.7%, compared to 5.9% in the rest of the male cohort). We provide a similar cancer risk analysis for all 101 PWAS gene associations (Supplementary Fig. S2). The capacity for risk assessment at the resolution of whole genes is enabled by PWAS through the aggregation of all variants affecting an individual. Integration of such data into the clinics could have a substantial impact on individual and familial cancer risk evaluation. Even when variants of unknown significance are observed within an individual, PWAS can still estimate gene damage and compare it to the population distribution, thereby providing risk assessment.

The presented analysis was conducted over nine specific cancer types and pan-cancer. We observe that most cancer predisposition loci are cancer-type specific. Only 13 of the 137 loci are shared by two or more specific cancer types, with 6 of them shared only by skin cancer and melanoma. Only 2 loci are shared by three or more specific cancer types (Fig. 4). The pan-cancer analysis recovered only 3 unique associations not discovered in specific cancers. This observation is in stark contrast to somatic tumor genomics, where most discovered cancer drivers are pan-cancer [35]. Our results suggest that cancer predisposition of germline genetic variation, unlike positive selection of somatic mutations in tumors, are mostly cancer-type specific. The cancer type with the highest number of recovered loci is skin cancer, with 49 of the 137 genomic regions associated with that cancer type. While this could be explained by the high number of skin cancer cases, it highlights a substantial polygenic signal for nonmelanoma skin cancer. Such strong polygenicity is compatible with the substantial role of environmental factors such as UV radiation and viral infection in this cancer [59].

An important limitation of our analysis is its restriction to the white British population, as the UKB cohort does not include a substantial number of non-white individuals required for sufficient statistical power. This leads to a lack of validation of the discovered associations on other populations, as well as oversight of additional associations only present in other ethnic groups. Another limitation of the UKB is the lack of sequencing data for the vast majority of the cohort. In this analysis we relied solely on genotyping of ∼800K predefined genetic markers and ∼600K variants in coding regions imputed from the original set of markers [32]. This limitation is particularly relevant for PWAS, which underestimates genetic damage when non-genotyped variants are involved, leading to diminished statistical power (but, critically, not to false discoveries) [29].

This study demonstrates the power of exhaustive genomic and proteomic analysis over large biobanks for expanding the discovery of cancer predisposition loci beyond family studies. In particular, we have demonstrated that recessive effects are also prevalent in case-control cohorts when looked for by suitable methodology. We have also demonstrated the benefit of cohorts with diverse cancer-type annotations for cross-cancer comparisons. Finally, our results show promise towards individual clinical risk assessment of cancer.

## Methods

### Study cohort

From the entire UK Biobank (UKB) cohort of 502,520 individuals (application ID 26664) [31, 32], we filtered out 227,690 individuals (see Fig. 1) for the following reasons: i) 14 individuals had asked to withdraw from the UKB, ii) 92,922 where non-whites (according to self-reported ethnicity or their genetics), iii) 312 had self-reported sex mismatching their genetics, iv) 75,848 were family-related to other individuals in the cohort (from each group of related individuals we chose only one representative), v) and 58,594 didn’t have any recorded ICD-10 codes (which were required to determine cancer history).

Following these filtrations, we remained with a study cohort of 274,830 individuals, 56,634 (20.06%) of them had a history of cancer according to their ICD-10 codes (the number of cases and controls per cancer type are listed in Table 2).

On top of cancer status, we also extracted from the UKB the following set of covariates: sex, year of birth, 40 genetic principal components, genotyping batch (105 categories) and UKB assessment center (25 categories). Altogether, 173 covariates (including a constant intercept) were included in the GWAS and PWAS analyses.

The computational pipeline for processing the UK Biobank data used in this work is an open-source project available at https://github.com/nadavbra/ukbb_parser.

### GWAS

To run GWAS, we used the PLINK software [60] (https://www.cog-genomics.org/plink/2.0/; version v2.00a2.3LM 64-bit Intel, 24 Jan 2020). For example, to run GWAS for breast cancer over chromosome 1, we ran the following command:

*plink2 --bed ukb_cal_chr1_v2.bed --bim ukb_snp_chr1_v2.bim --fam ukb2666_cal_chr1_v2_s488366.fam --pheno breast_cancer.txt --covar covariates.txt --out chr1_breast_cancer --1 --glm hide-covar no-x-sex --mac 20 --covar-variance-standardize --freq --threads 32 --memory 100000*

Where *ukb_cal_chr1_v2.bed, ukb_snp_chr1_v2.bim* and *ukb2666_cal_chr1_v2_s488366.fam* are the files with the genetic data of chr1, provided by the UKB. The files *breast_cancer.txt* and *covariates.txt* are PLINK-formatted tab-separated data files with the cancer phenotype (case-control status) and covariates of each of the samples in the study cohort, respectively.

We ran GWAS independently over ten cancer phenotypes (nine specific cancer types and pan-cancer) and over 26 chromosomes (22 autosomal chromosomes, chrX, chrY, chrXY and chrMY, as provided by the UKB). The complete GWAS results with all the summary statistics, including 861 variant-cancer associations below the exome-wide significance threshold (5E-07), are available per cancer-type in Supplementary Table S2.

### PWAS

We used the PWAS software [29] (version 1.0.4; available at https://github.com/nadavbra/pwas) and executed the standard PWAS pipeline (specified in the GitHub page). We derived the dominant and recessive PWAS effect scores for the entire UKB cohort (which depend solely on the genetic information) and tested associations between 18,053 protein-coding genes and the ten cancer phenotypes. The complete PWAS results with the full summary statistics, which include 101 FDR-significant gene-cancer associations, are available per cancer-type in Supplementary Table S3.

### Merging the significant associations into unique genomic regions

We merged the 861 significant GWAS associations and 101 significant PWAS associations into 137 unique genomic regions by extending each association by 500,000 base-pairs on both sides and iteratively merging overlapping regions. GWAS associations started as 1bp and were extended into 1,000,001bp regions (the position of the variant, plus 500,000 bp on each side) while PWAS associations started as the region of the gene’s coding-sequence (i.e. from the genomic coordinate of the start codon to the last before the stop codon). This merging procedure resulted in 137 non-overlapping genomic loci of sizes of at least 1Mbp, available in Supplementary Table S1. The largest obtained locus was chr6:30,145,049-34,892,998 (of size 4.7 Mbp), which covers the MHC region. This region contains 7,049 variants genotyped by the UKB (207 of which are GWAS-significant) and 138 protein-coding genes tested by PWAS (6 of them are PWAS-significant).

### External cancer evidence

We validated each of the 137 cancer predisposition regions against three catalogues of cancer drivers (Census [33], pan-software [34] and FABRIC [35]) and two gene panels used in the clinics (CleanPlex TMB 500 and Invitae Multi-Cancer). The “Cancer Gene Census” was downloaded as a CSV file (cancer_gene_census.csv, from https://cancer.sanger.ac.uk/cosmic/download) containing 723 genes. Each of the Census genes is identified as a tumor-suppressor gene (TSG), oncogene or a fusion gene (or any combination of these three categories). The list of 515 genes in the CleanPlex TMB 500 panel was downloaded from https://www.paragongenomics.com/wp-content/uploads/2019/04/PS1008_CleanPlex_TMB-500-Panel_Gene-List.txt, and the list of 83 genes in the Invitae Multi-Cancer panel was downloaded from https://www.invitae.com/en/physician/tests/01101/#info-panel-assay_information. The list of 83 Invitae Multi-Cancer genes was further extended by 50 extra genes, contributed by Vardiela Meiner from the Center for Clinical Genetics in Hadassah Medical Center (Supplementary Table S4). The Census gene list was downloaded in May 2020, and the two panel gene lists were downloaded in June 2020. The list of 299 pan-software driver genes was downloaded from Supplementary Table S1 of the publication [34]. The list of 593 FABRIC driver genes was downloaded from Supplementary Table S1 of the publication [35]. We mapped the obtained lists of genes onto our 18,053 analyzed genes according to their UniProt ID (in the case of FABRIC) or gene symbols (in the case of the other resources). For each of the 137 genomic regions listed in Supplementary Table S1, we include all the genes in the region that are supported by each of these external catalogues.

In Fig. 2, we consider a genomic locus to be supported by clinical panels or driver catalogues if it contains at least one gene listed in those external resources. In Table 1 and Table 3, we consider the external evidence to be relevant only if it matches the exact gene listed on the table.

In Table 3 we also include cancer associations of the listed genes according to the Open Targets Platform [57]. We consider a reported association to be relevant only if i) it is related to cancer or neoplasm, and ii) it is supported by genetic association (as opposed to the overall association score reported by the platform, which also includes other sources of evidence, including somatic mutations).

### Cancer risk as a function of PWAS effect scores

In Fig. 5B and Supplementary Fig. S2 we illustrate the clinical significance of PWAS associations by plotting the risk of the relevant cancer type as a function of dominant and recessive effect scores (which capture the overall amount of gene damage to the relevant gene, per individual). The underlying data for these figures (cancer case-control status and PWAS effect scores) was taken directly from the study cohort (see the relevant sections in the Methods). In the continuous plots (the two left columns in these figures), we examine the entire distribution of PWAS effect scores by considering all possible cutoffs of the relevant cohort, i.e. all subsets with effect scores that are either below or above a certain threshold. Since the precise effect-score values are not so informative, we describe the relevant sub-populations by percentiles (e.g. “bottom 0.9%”, referring to the 0.9% of the cohort with the lowest scores, namely the 9% with the greatest estimated functional damage to the gene, or “top 5%”, referring to the 5% of the cohort with the highest scores, namely the 5% with the least estimated functional damage to the gene). Each of the continuous plots also includes a reference to the entire cohort, marked on the x-axis as “All”. Cutoffs of bottom percentiles are shown to the left of the entire-cohort reference point, and cutoffs of top percentiles are shown to its right (both in log scale). 95% confidence intervals for the fraction of cases in each subpopulation were calculated using Wilson score interval’s method for binomial proportion.

In addition to the continuous plots that capture the entire distribution of PWAS effect scores, we also illustrate the associations by choosing specific cutoffs showing the strongest differences. Specifically, we tested each cutoff by Fisher’s exact test for the association between the effect-score cutoff to cancer case-control status, selecting the cutoff with the lowest p-value (and display the plot only if it’s lower than 1E-03).

Note that these plots do not account for covariates, and should therefore not be considered as rigorous statistical evidence for the associations (unlike the summary p-values reported by PWAS, which account for covariates and consider the entire distribution of effect scores).

## Supporting information

Supplementary Table S1 - Significant Regions

Supplementary Table S4 - Extra 50 Panel Genes

Supplementary Table S3 - Full PWAS Results

## Acknowledgements

We would like to thank Vardiela Meiner from the Center for Clinical Genetics in Hadassah Medical Center for fruitful discussion and valuable feedback on our work. We would also like to thank Roni Rasnic from the Hebrew University for important comments on our manuscript.

## Supplementary Materials

**Supplementary Table S1**: The full list of 137 significant cancer predisposition genomic regions.

**Supplementary Table S2**: Full GWAS results (PLINK’s summary statistics across all ten cancer types and 770,783 genetic markers).

**Supplementary Table S3**: Full PWAS results (summary statistics across all ten cancer types and 18,053 analyzed protein-coding genes).

**Supplementary Table S4**: A list of 50 extra genes supplementing the Invitae Multi-Cancer clinical gene panel.

**Supplementary Fig. S1**: Genome-wide cancer predisposition in all ten cancer-types (an extension of Fig. 3).

**Supplementary Fig. S2**: Cancer risk across all 101 PWAS-significant genes (an extension of Fig. 5B).

