## Supplementary Figures for "Recessive effects in cancer predisposition exposed by genome-wide and proteome-wide association studies"

**Contents:**

Supplementary Fig. S1: Genome-wide cancer predisposition in all ten cancer-types (an extension of Fig. 3 in the main text; page 2).

Supplementary Fig. S2: Cancer risk across all 101 PWAS-significant genes (an extension of Fig. 5B in the main text; page 6).

Supplementary Fig. S1: Genome-wide cancer predisposition in all ten cancer-types

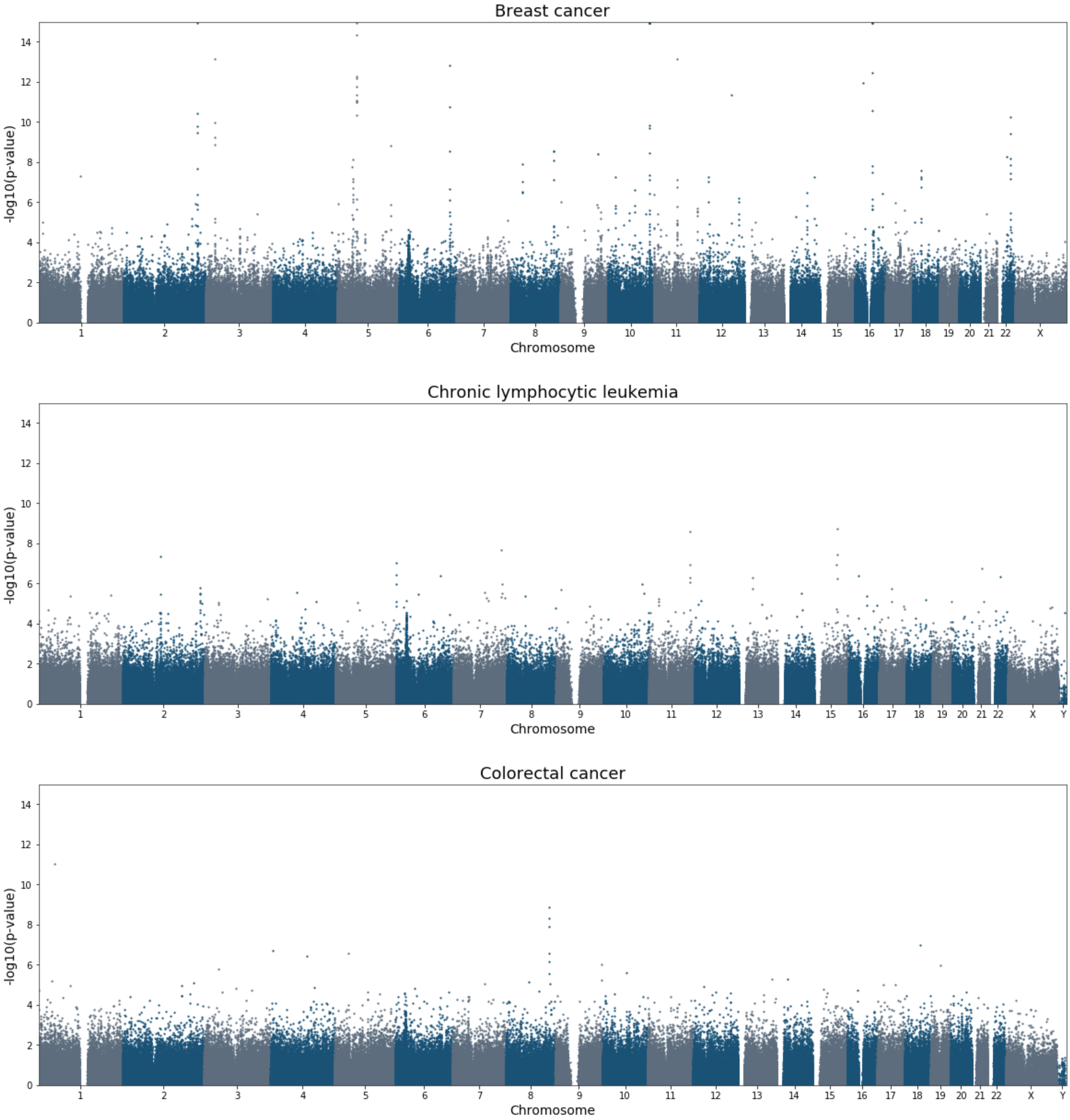

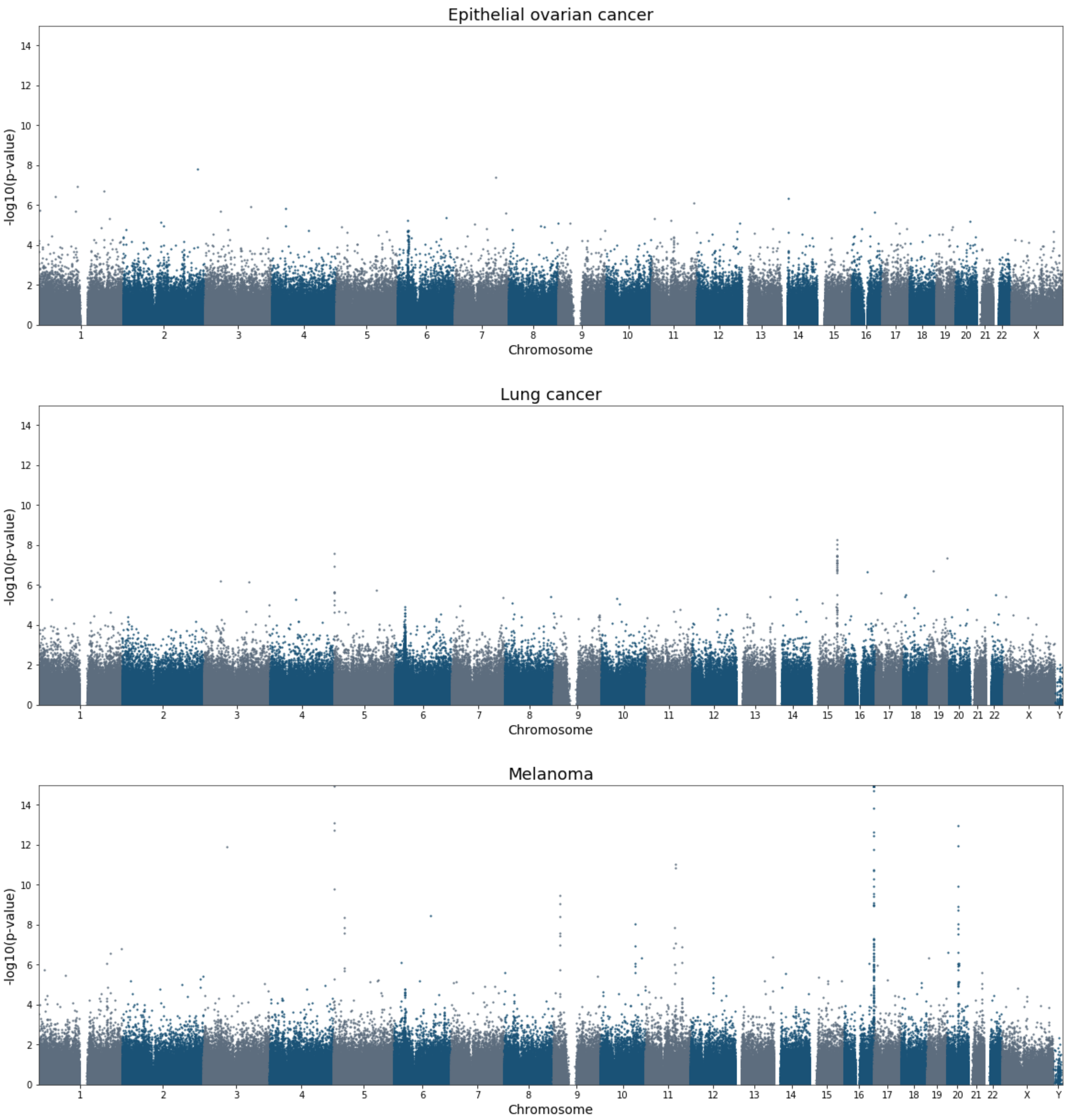
Epithelial ovarian cancer
-log10(p-value)
Chromosome
Lung cancer
-log10(p-value)
Chromosome
Melanoma
-log10(p-value)
Chromosome

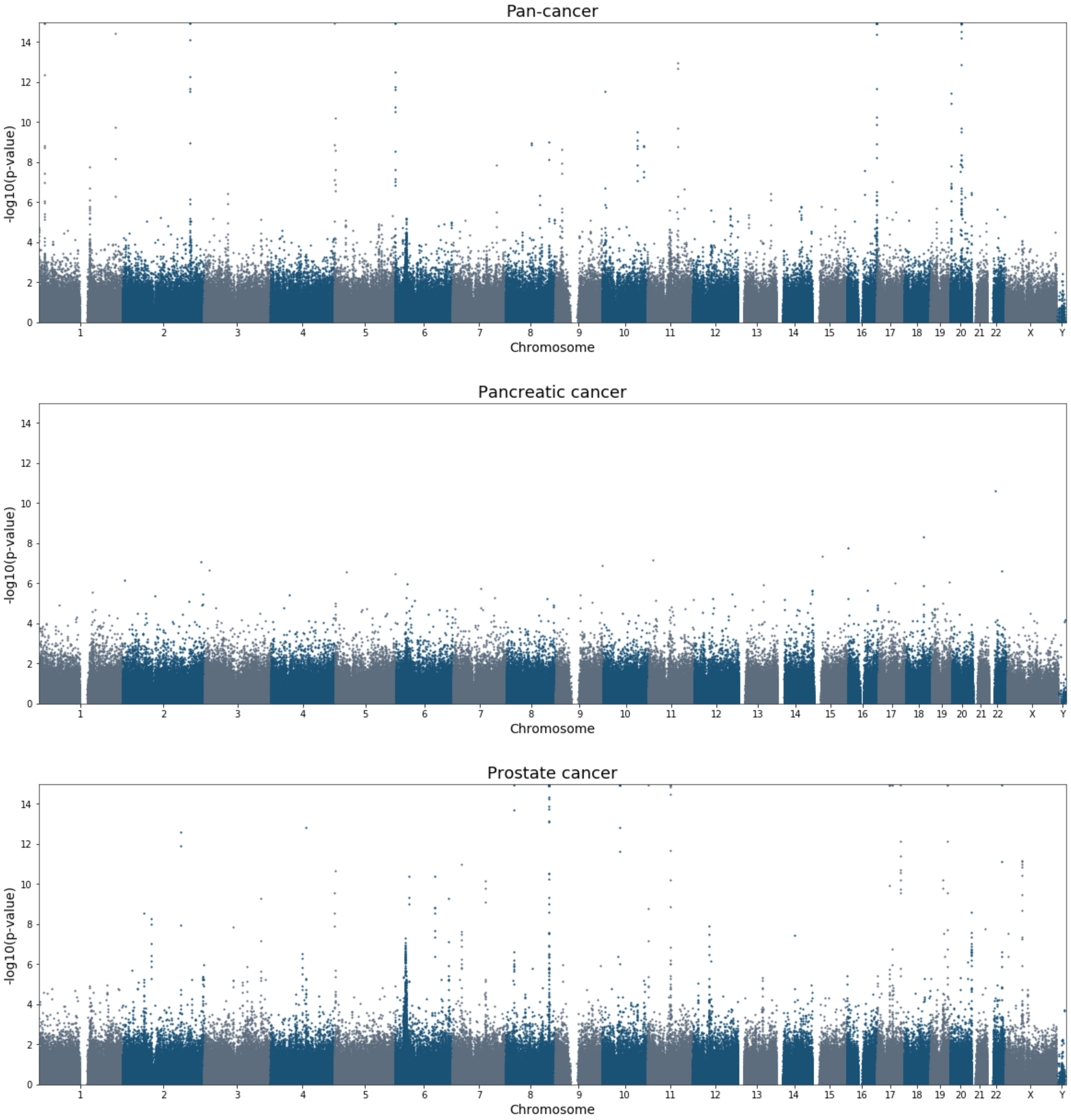
Pan-cancer
Pancreatic cancer
Prostate cancer
-log10(p-value)
Chromosome
0
2
4
6
8
10
12
14
X
Y

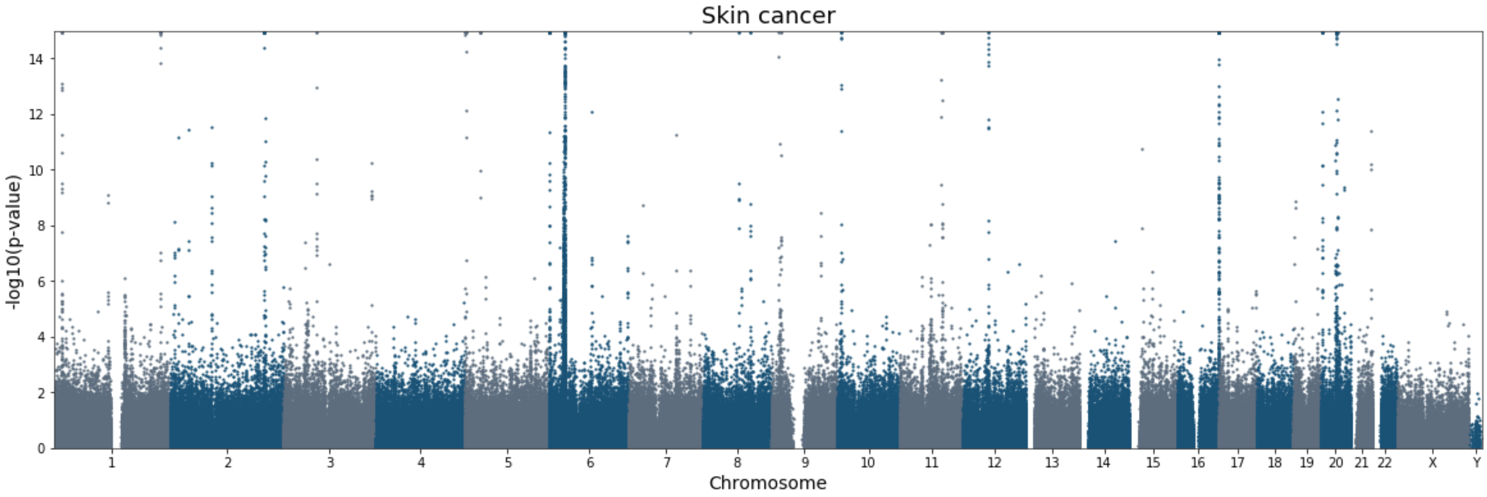
Skin cancer
-log10(p-value)
Chromosome
0
2
4
6
8
10
12
14
X
Y

### Supplementary Fig. S2: Cancer risk across all 101 PWAS-significant genes

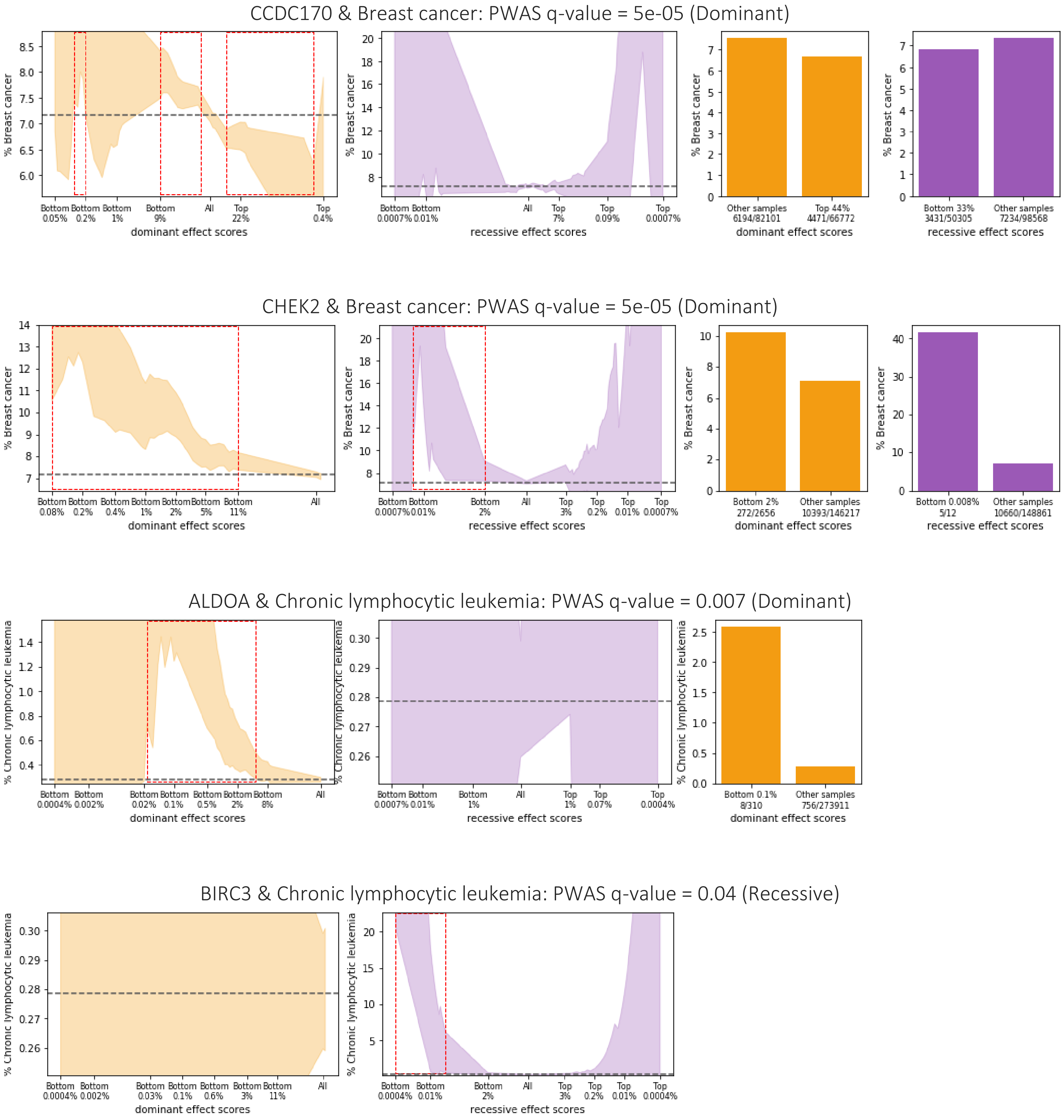

CCT8 & Chronic lymphocytic leukemia: PWAS q-value = 0.02 (Recessive)

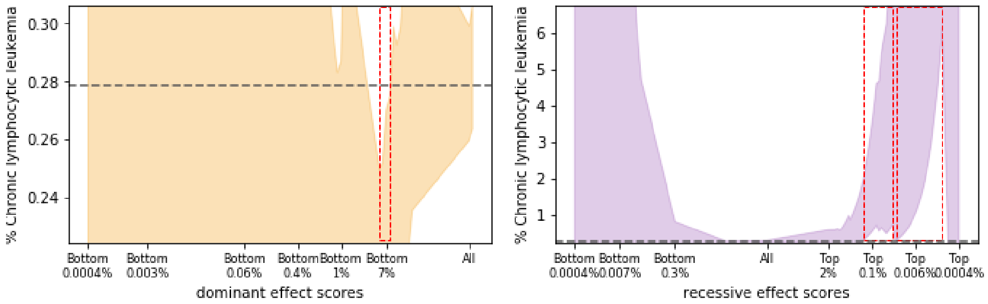

CTNNAL1 & Chronic lymphocytic leukemia: PWAS q-value = 0.05 (Dominant)

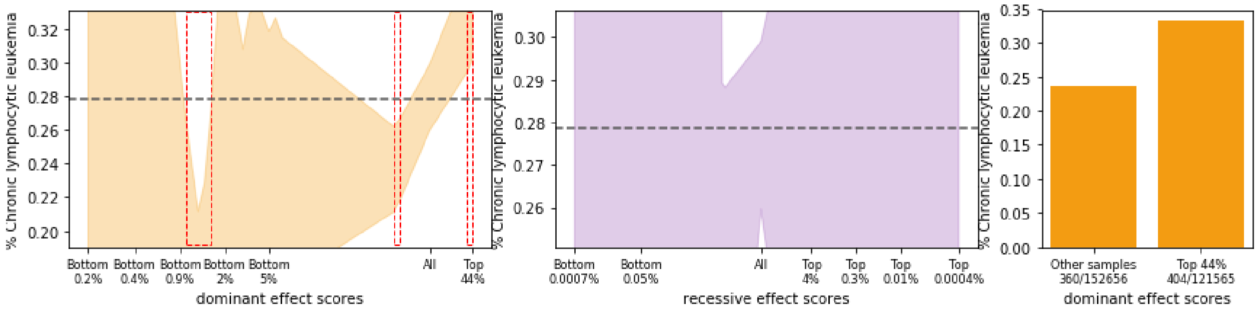

DPT & Chronic lymphocytic leukemia: PWAS q-value = 0.04 (Recessive)

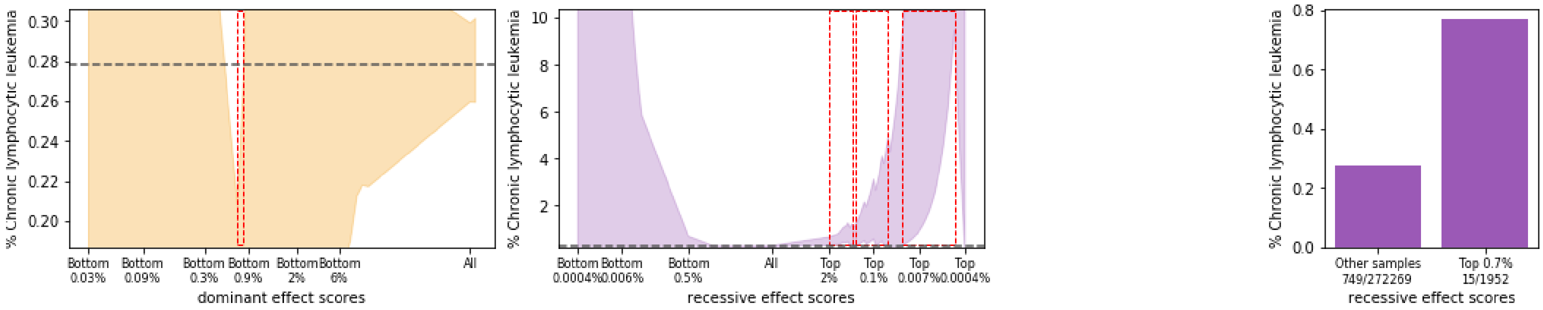

FAM160B1 & Chronic lymphocytic leukemia: PWAS q-value = 0.009 (None)

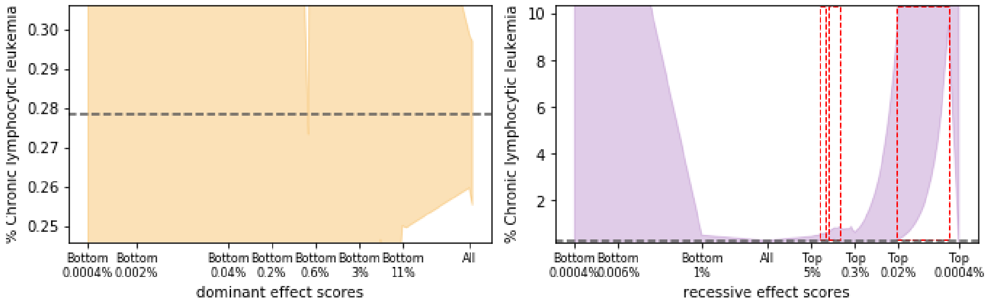

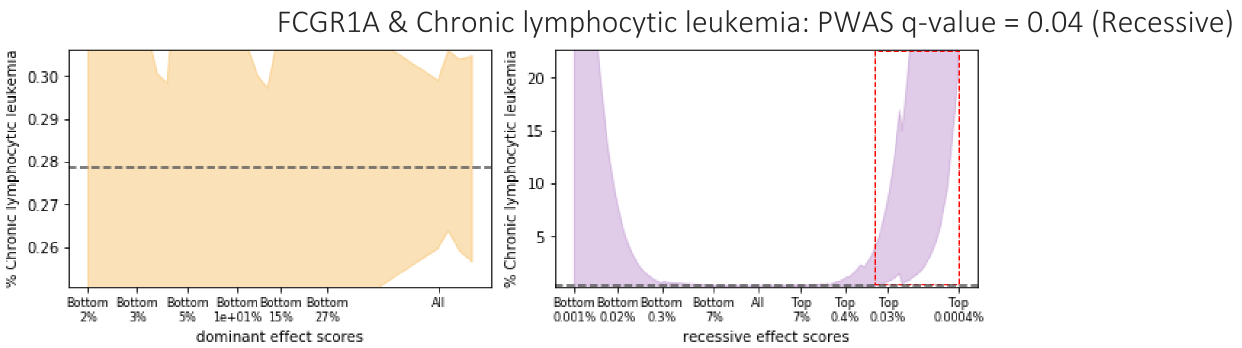
FCGR1A & Chronic lymphocytic leukemia: PWAS q-value = 0.04 (Recessive)
% Chronic lymphocytic leukemia
dominant effect scores
Bottom 2%
Bottom 3%
Bottom 5%
Bottom 1e+01%
Bottom 15%
Bottom 27%
All
% Chronic lymphocytic leukemia
recessive effect scores
Bottom 0.001%
Bottom 0.02%
Bottom 0.3%
Bottom 7%
All
Top 7%
Top 0.4%
Top 0.03%
Top 0.0004%

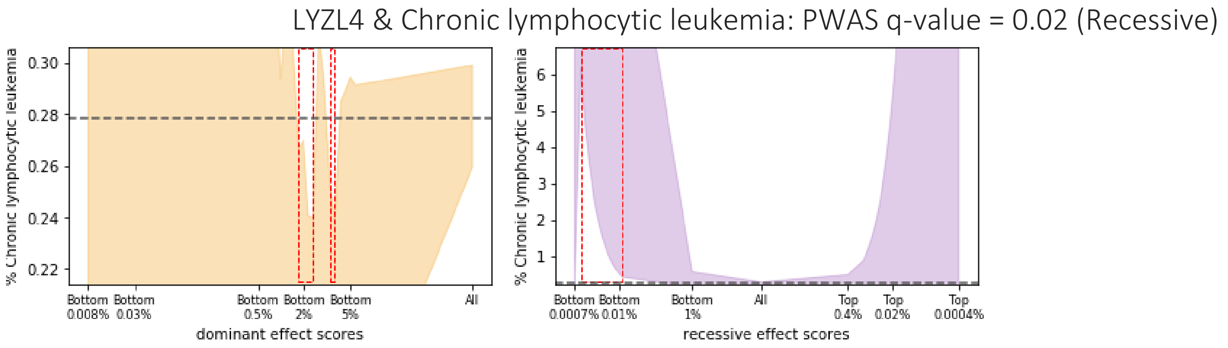
LYZL4 & Chronic lymphocytic leukemia: PWAS q-value = 0.02 (Recessive)
% Chronic lymphocytic leukemia
dominant effect scores
Bottom 0.008%
Bottom 0.03%
Bottom 0.5%
Bottom 2%
Bottom 5%
All
% Chronic lymphocytic leukemia
recessive effect scores
Bottom 0.0007%
Bottom 0.01%
Bottom 1%
All
Top 0.4%
Top 0.02%
Top 0.0004%

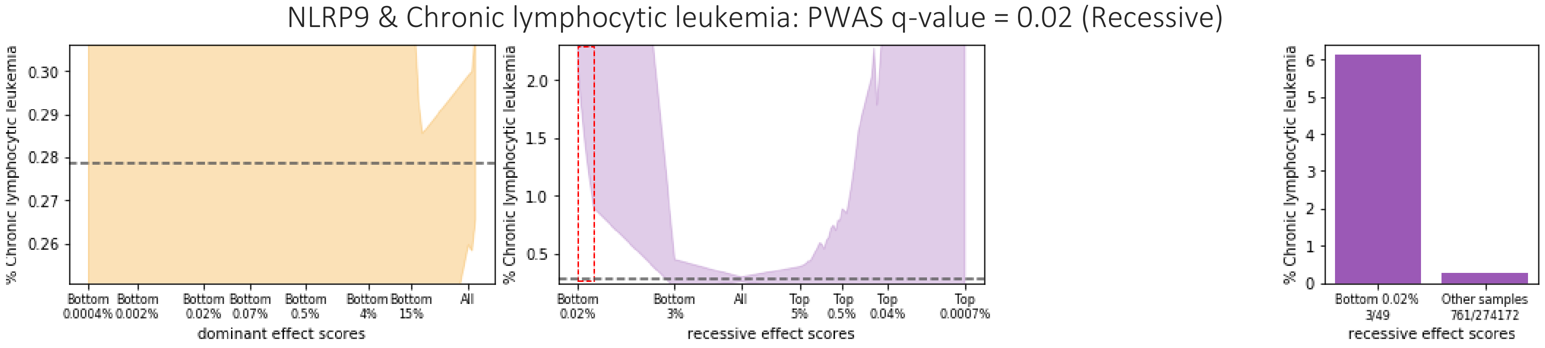
NLRP9 & Chronic lymphocytic leukemia: PWAS q-value = 0.02 (Recessive)
% Chronic lymphocytic leukemia
dominant effect scores
Bottom 0.0004%
Bottom 0.002%
Bottom 0.02%
Bottom 0.07%
Bottom 0.5%
Bottom 4%
Bottom 15%
All
% Chronic lymphocytic leukemia
recessive effect scores
Bottom 0.02%
Bottom 3%
All
Top 5%
Top 0.5%
Top 0.04%
Top 0.0007%
% Chronic lymphocytic leukemia
Bottom 0.02%
3/49
Other samples
761/274172
recessive effect scores

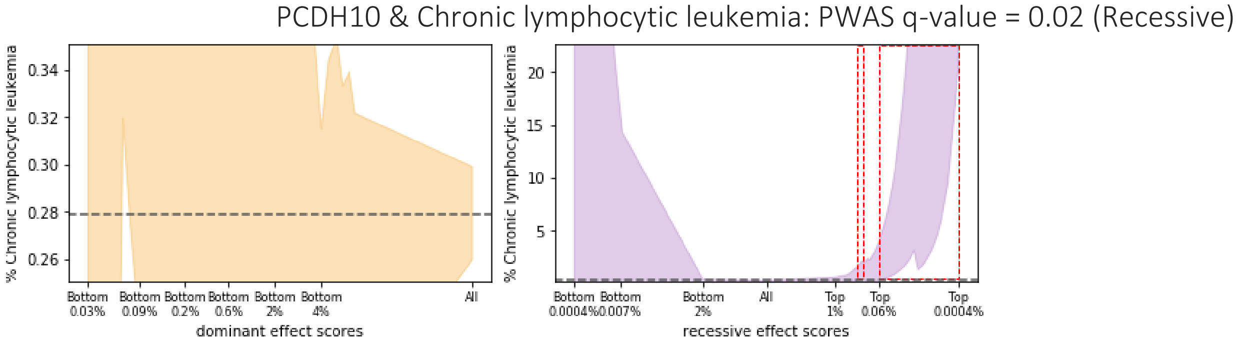
PCDH10 & Chronic lymphocytic leukemia: PWAS q-value = 0.02 (Recessive)
% Chronic lymphocytic leukemia
dominant effect scores
Bottom 0.03%
Bottom 0.09%
Bottom 0.2%
Bottom 0.6%
Bottom 2%
Bottom 4%
All
% Chronic lymphocytic leukemia
recessive effect scores
Bottom 0.0004%
Bottom 0.007%
Bottom 2%
All
Top 1%
Top 0.06%
Top 0.0004%

RBMX2 & Chronic lymphocytic leukemia: PWAS q-value = 0.03 (Recessive)

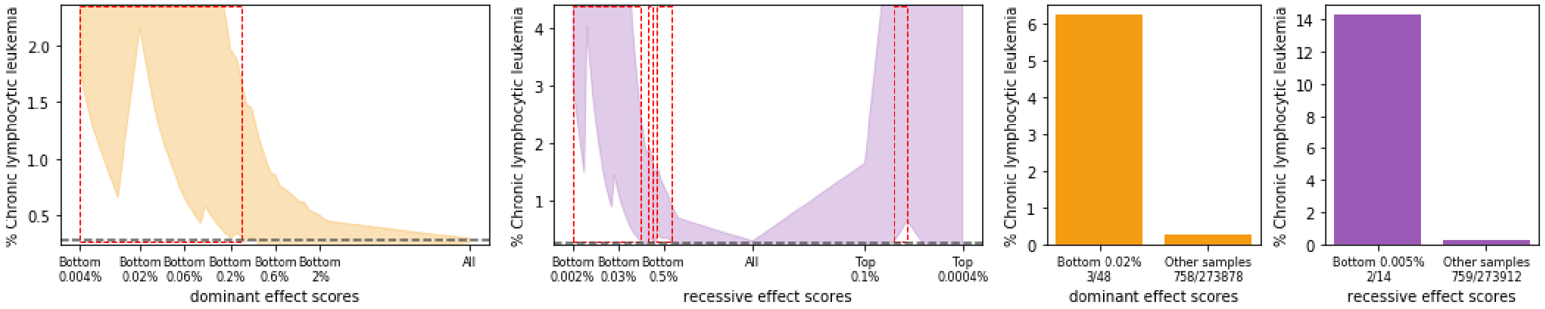

RCHY1 & Chronic lymphocytic leukemia: PWAS q-value = 0.01 (Recessive)

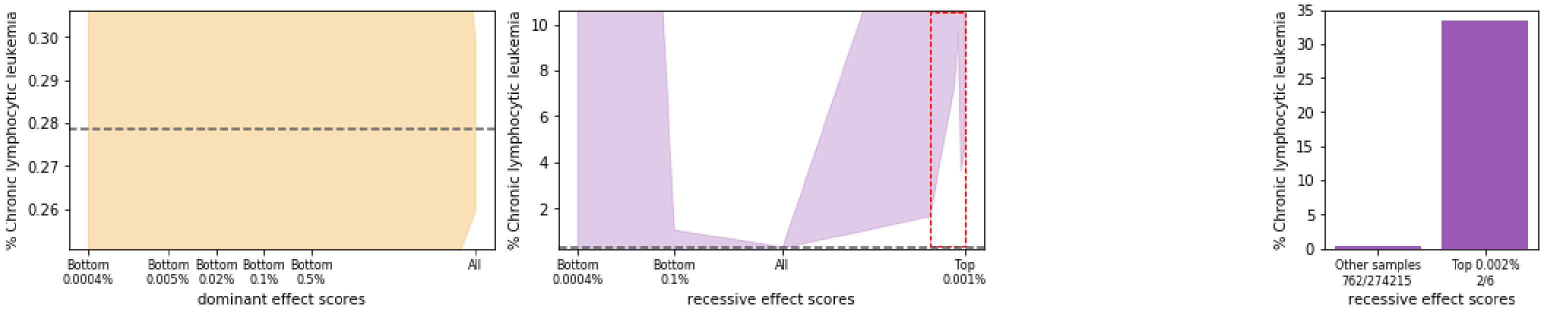

SGSM3 & Chronic lymphocytic leukemia: PWAS q-value = 0.05 (Recessive)

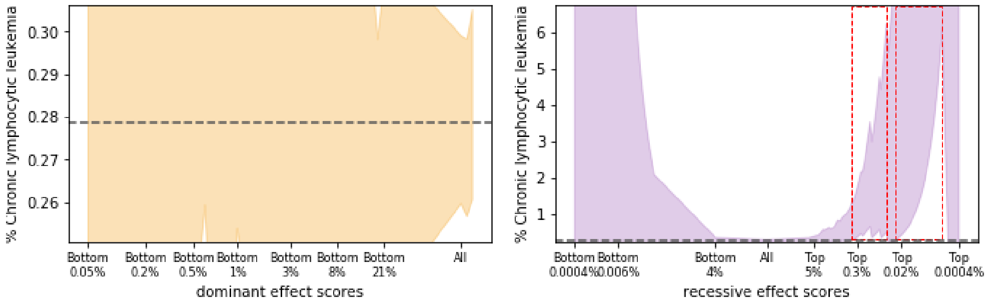

SRSF5 & Chronic lymphocytic leukemia: PWAS q-value = 0.01 (Dominant)

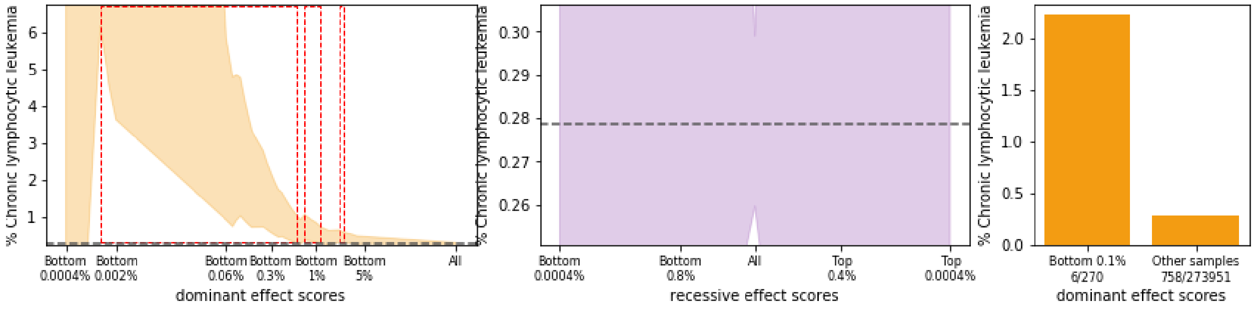

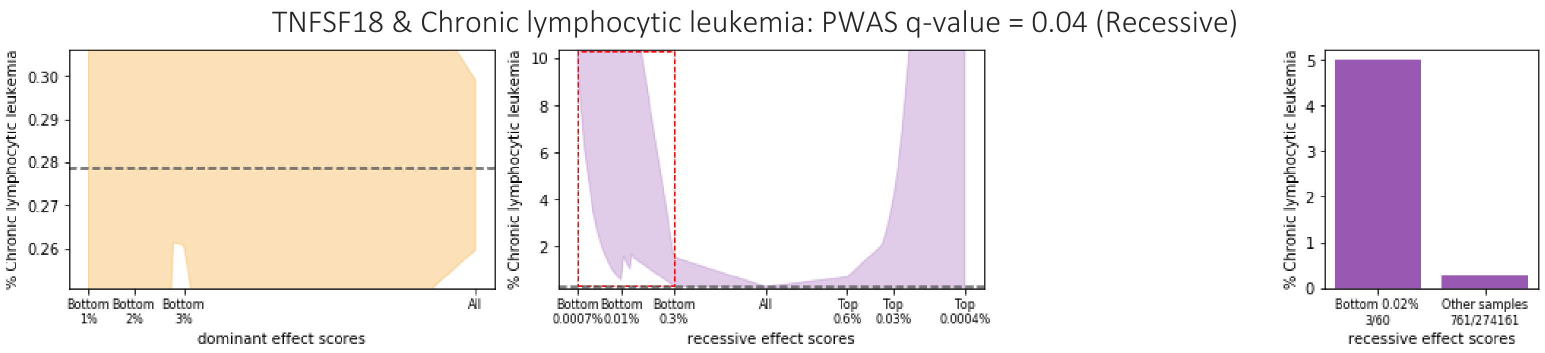
TNFSF18 & Chronic lymphocytic leukemia: PWAS q-value = 0.04 (Recessive)
% Chronic lymphocytic leukemia
dominant effect scores
recessive effect scores
Bottom 0.02% 3/60
Other samples 761/274161

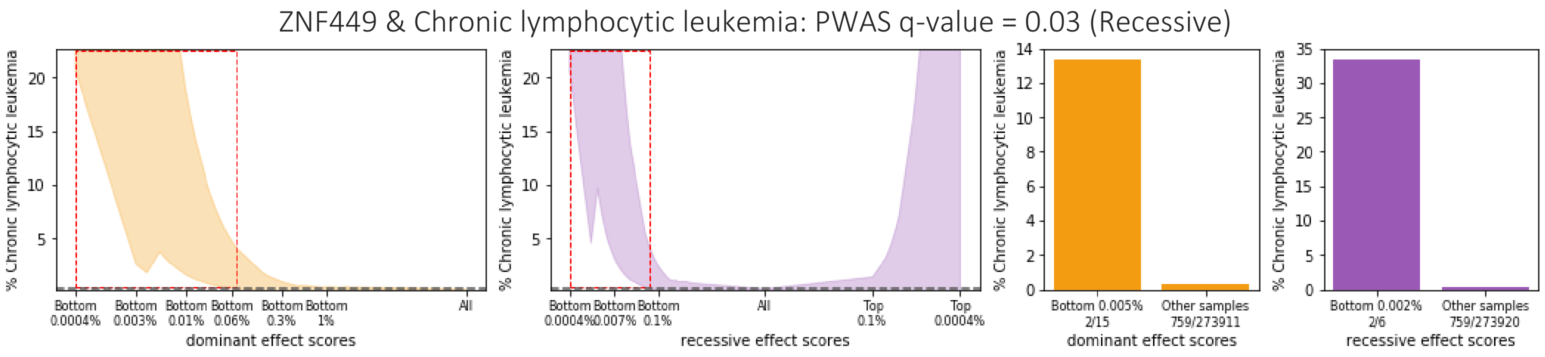
ZNF449 & Chronic lymphocytic leukemia: PWAS q-value = 0.03 (Recessive)
% Chronic lymphocytic leukemia
dominant effect scores
recessive effect scores
Bottom 0.005% 2/15
Other samples 759/273911
Bottom 0.002% 2/6
Other samples 759/273920

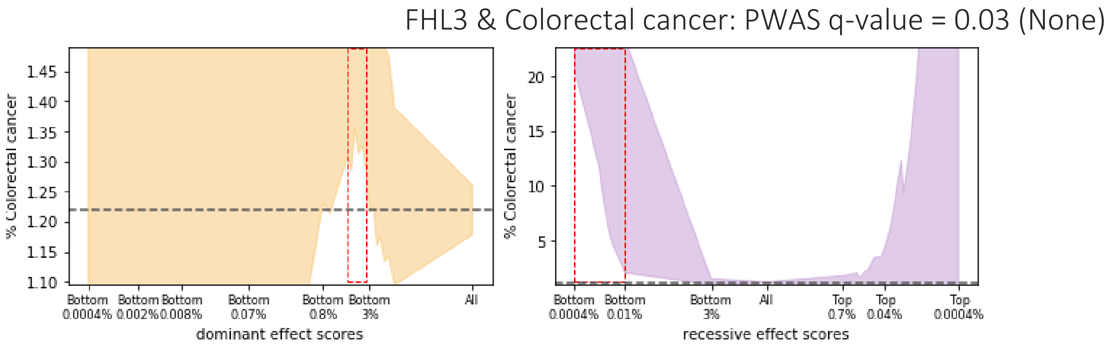
FHL3 & Colorectal cancer: PWAS q-value = 0.03 (None)
% Colorectal cancer
dominant effect scores
recessive effect scores

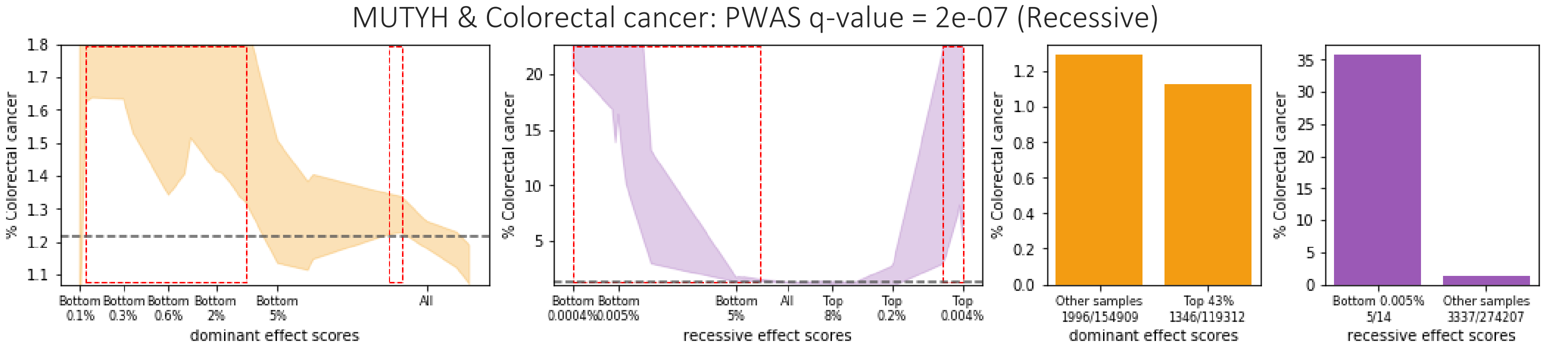
MUTYH & Colorectal cancer: PWAS q-value = 2e-07 (Recessive)
% Colorectal cancer
dominant effect scores
recessive effect scores
Other samples 1996/154909
Top 43% 1346/119312
Bottom 0.005% 5/14
Other samples 3337/274207

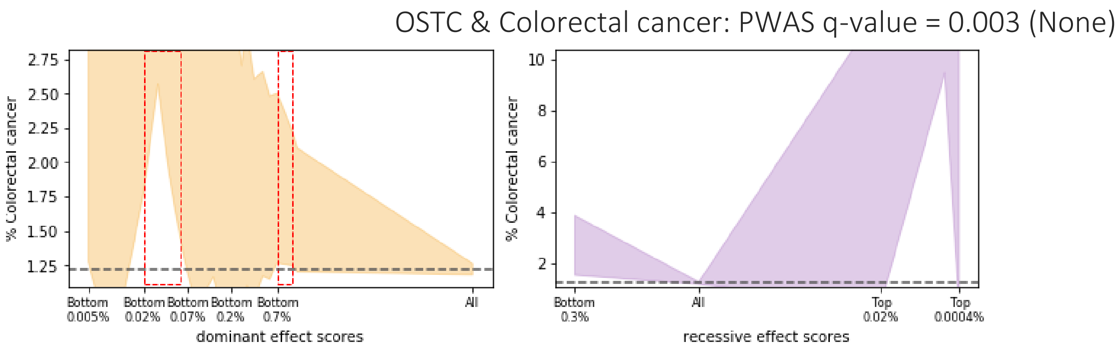
OSTC & Colorectal cancer: PWAS q-value = 0.003 (None)
% Colorectal cancer
Bottom 0.005%
Bottom 0.02%
Bottom 0.07%
Bottom 0.2%
Bottom 0.7%
All
dominant effect scores
Bottom 0.3%
All
Top 0.02%
Top 0.0004%
recessive effect scores

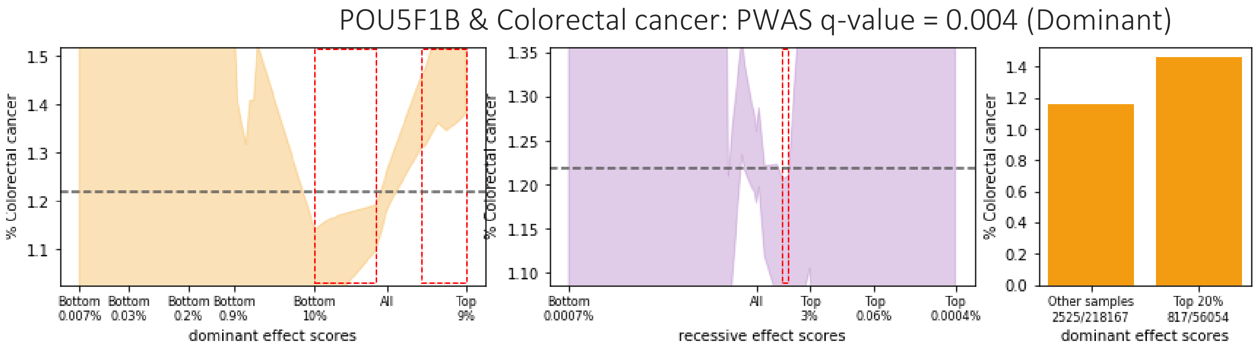
POU5F1B & Colorectal cancer: PWAS q-value = 0.004 (Dominant)
% Colorectal cancer
Bottom 0.007%
Bottom 0.03%
Bottom 0.2%
Bottom 0.9%
Bottom 10%
All
Top 9%
dominant effect scores
Bottom 0.0007%
All
Top 3%
Top 0.06%
Top 0.0004%
recessive effect scores
Other samples 2525/218167
Top 20% 817/56054
dominant effect scores

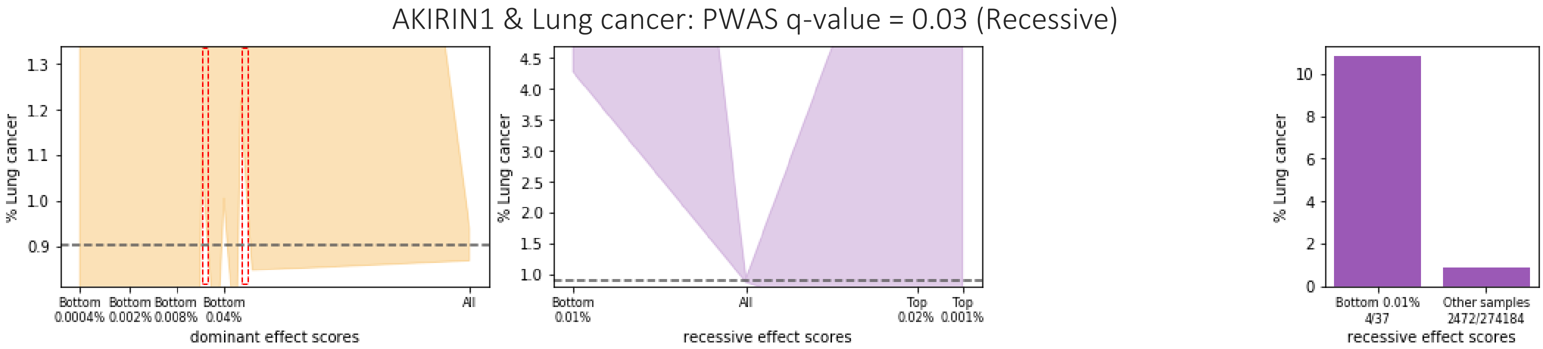
AKIRIN1 & Lung cancer: PWAS q-value = 0.03 (Recessive)
% Lung cancer
Bottom 0.0004%
Bottom 0.002%
Bottom 0.008%
Bottom 0.04%
All
dominant effect scores
Bottom 0.01%
All
Top 0.02%
Top 0.001%
recessive effect scores
Bottom 0.01% 4/37
Other samples 2472/274184
recessive effect scores

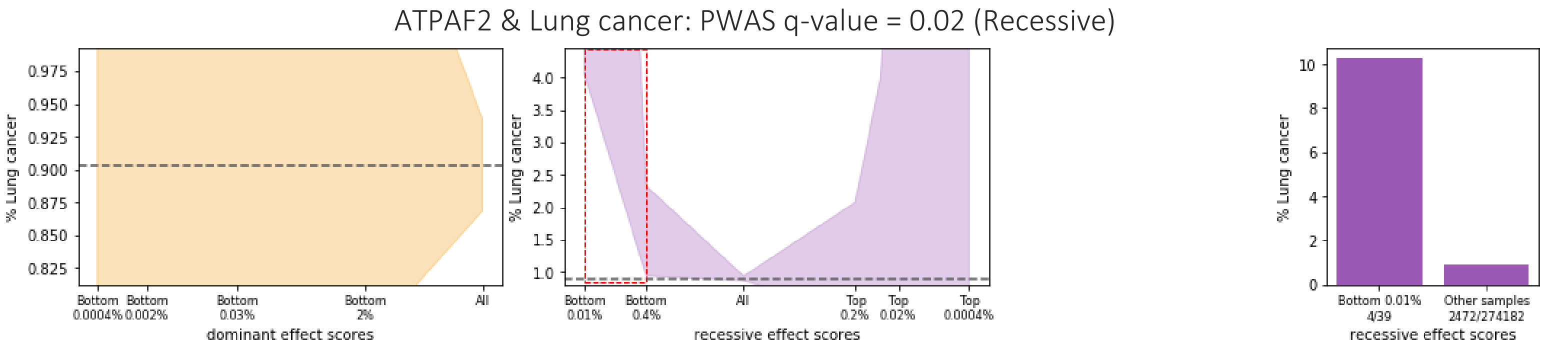
ATPAF2 & Lung cancer: PWAS q-value = 0.02 (Recessive)
% Lung cancer
Bottom 0.0004%
Bottom 0.002%
Bottom 0.03%
Bottom 2%
All
dominant effect scores
Bottom 0.01%
Bottom 0.4%
All
Top 0.2%
Top 0.02%
Top 0.0004%
recessive effect scores
Bottom 0.01% 4/39
Other samples 2472/274182
recessive effect scores

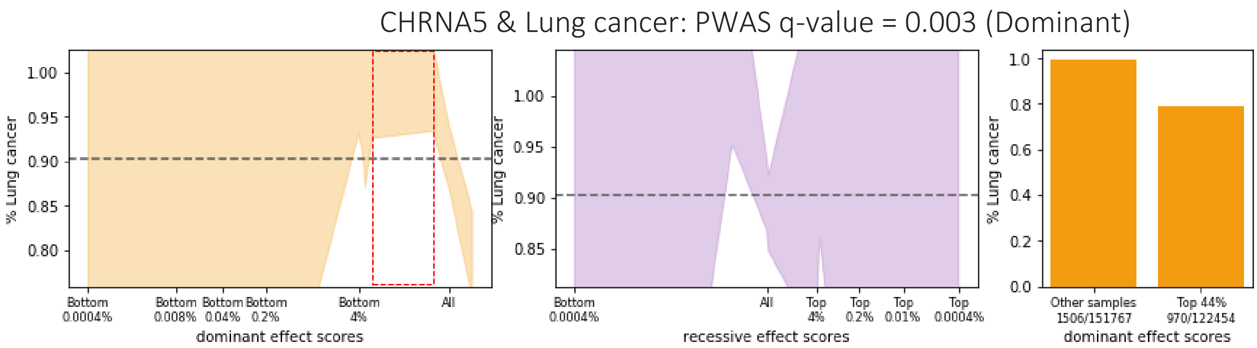
CHRNA5 & Lung cancer: PWAS q-value = 0.003 (Dominant)
% Lung cancer
Bottom 0.0004%
Bottom 0.008%
Bottom 0.04%
Bottom 0.2%
Bottom 4%
All
dominant effect scores
Bottom 0.0004%
All
Top 4%
Top 0.2%
Top 0.01%
Top 0.0004%
recessive effect scores
Other samples 1506/151767
Top 44% 970/122454
dominant effect scores

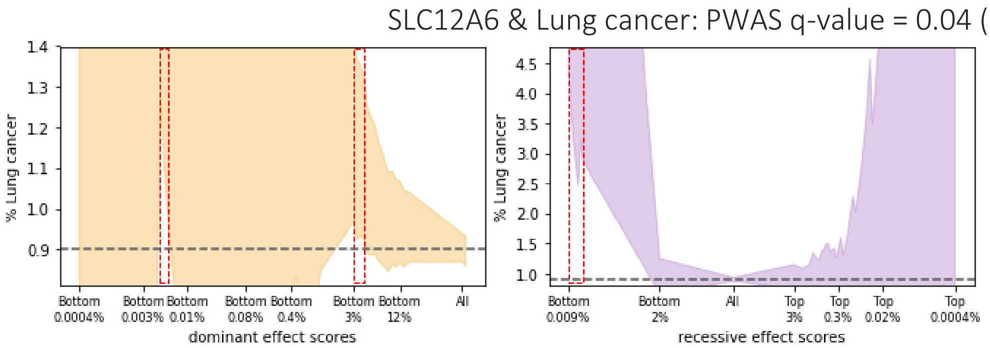
SLC12A6 & Lung cancer: PWAS q-value = 0.04 (Recessive)
% Lung cancer
Bottom 0.0004%
Bottom 0.003%
Bottom 0.01%
Bottom 0.08%
Bottom 0.4%
Bottom 3%
Bottom 12%
All
dominant effect scores
Bottom 0.009%
Bottom 2%
All
Top 3%
Top 0.3%
Top 0.02%
Top 0.0004%
recessive effect scores

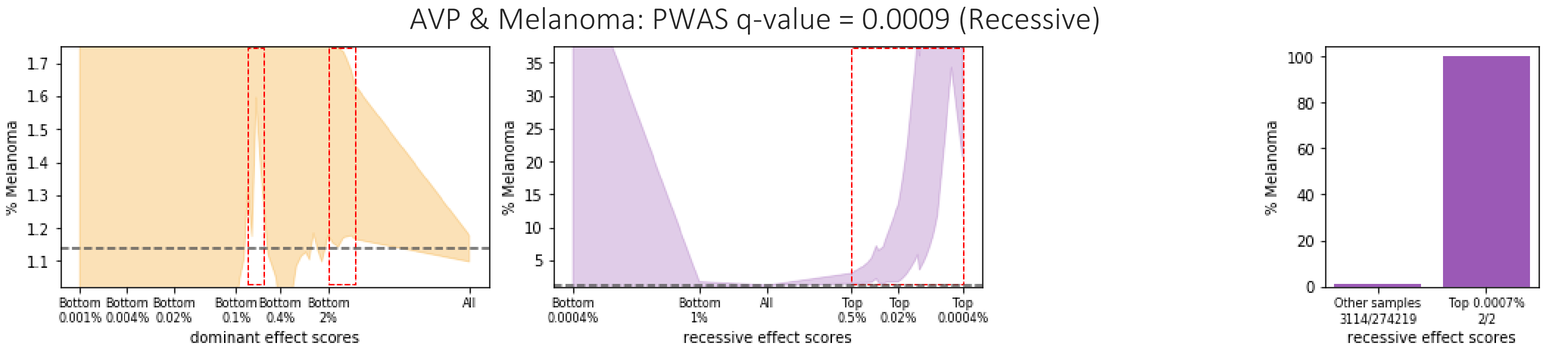
AVP & Melanoma: PWAS q-value = 0.0009 (Recessive)
% Melanoma
Bottom 0.001%
Bottom 0.004%
Bottom 0.02%
Bottom 0.1%
Bottom 0.4%
Bottom 2%
All
dominant effect scores
Bottom 0.0004%
Bottom 1%
All
Top 0.5%
Top 0.02%
Top 0.0004%
recessive effect scores
Other samples 3114/274219
Top 0.0007% 2/2
recessive effect scores

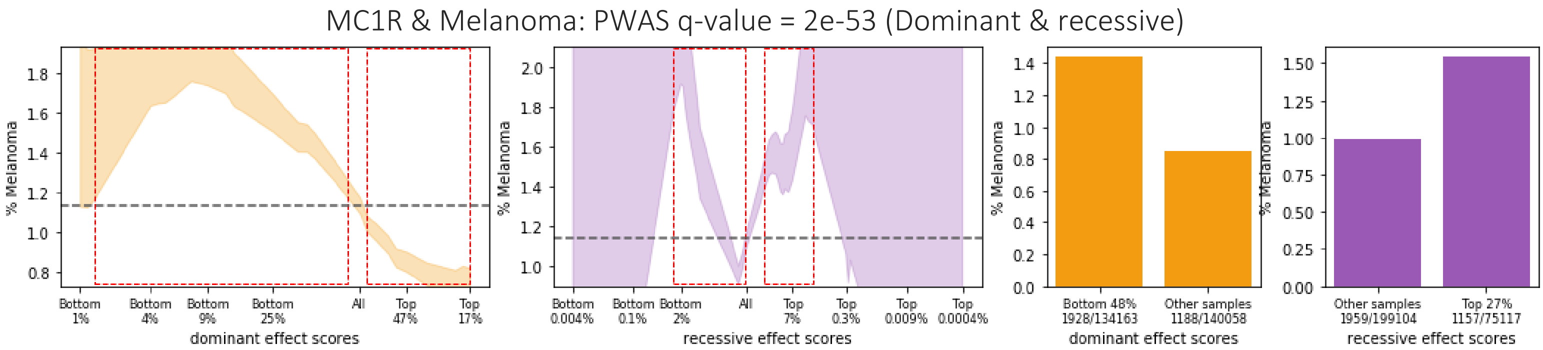
MC1R & Melanoma: PWAS q-value = 2e-53 (Dominant & recessive)
% Melanoma
Bottom 1%
Bottom 4%
Bottom 9%
Bottom 25%
All
Top 47%
Top 17%
dominant effect scores
Bottom 0.004%
Bottom 0.1%
Bottom 2%
All
Top 7%
Top 0.3%
Top 0.009%
Top 0.0004%
recessive effect scores
Bottom 48% 1928/134163
Other samples 1188/140058
dominant effect scores
Other samples 1959/199104
Top 27% 1157/75117
recessive effect scores

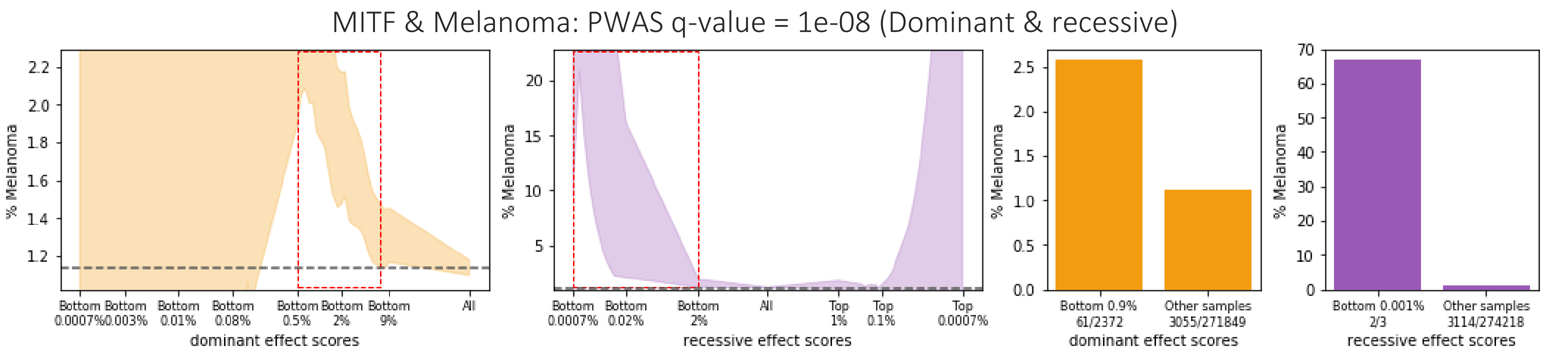
MITF & Melanoma: PWAS q-value = 1e-08 (Dominant & recessive)
% Melanoma
dominant effect scores
recessive effect scores
Bottom 0.9% 61/2372
Other samples 3055/271849
Bottom 0.001% 2/3
Other samples 3114/274218

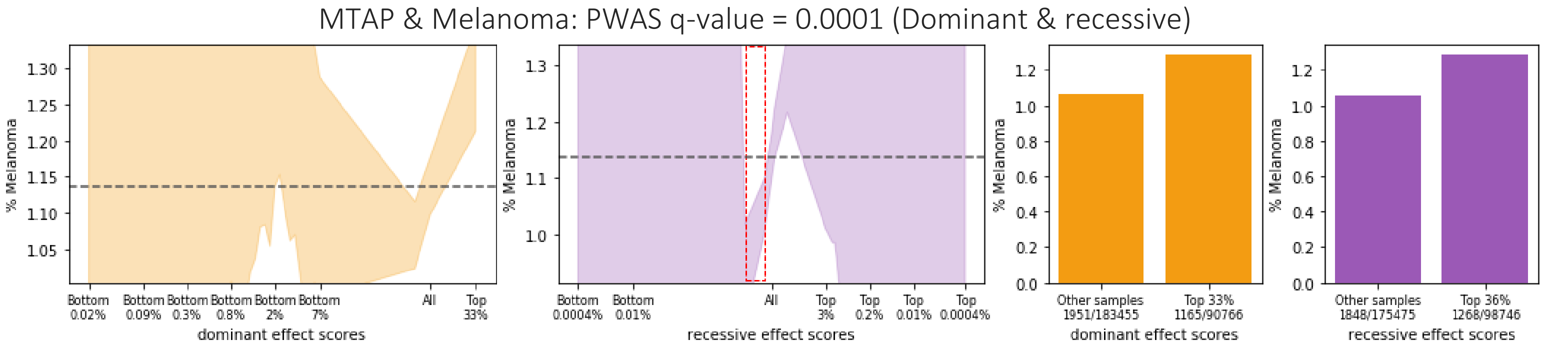
MTAP & Melanoma: PWAS q-value = 0.0001 (Dominant & recessive)
% Melanoma
dominant effect scores
recessive effect scores
Other samples 1951/183455
Top 33% 1165/90766
Other samples 1848/175475
Top 36% 1268/98746

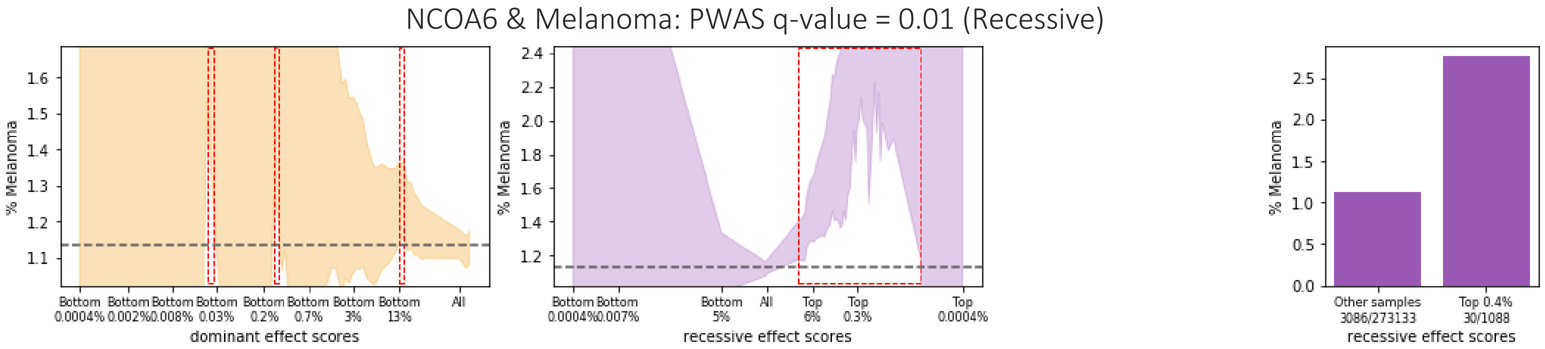
NCOA6 & Melanoma: PWAS q-value = 0.01 (Recessive)
% Melanoma
dominant effect scores
recessive effect scores
Other samples 3086/273133
Top 0.4% 30/1088

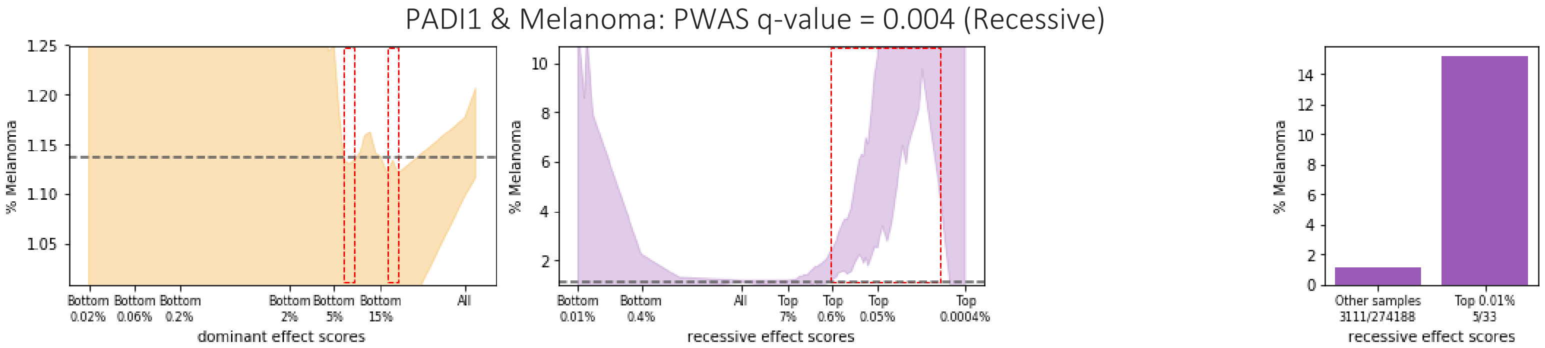
PADI1 & Melanoma: PWAS q-value = 0.004 (Recessive)
% Melanoma
dominant effect scores
recessive effect scores
Other samples 3111/274188
Top 0.01% 5/33

POLR2H & Melanoma: PWAS q-value = 0.01 (Recessive)

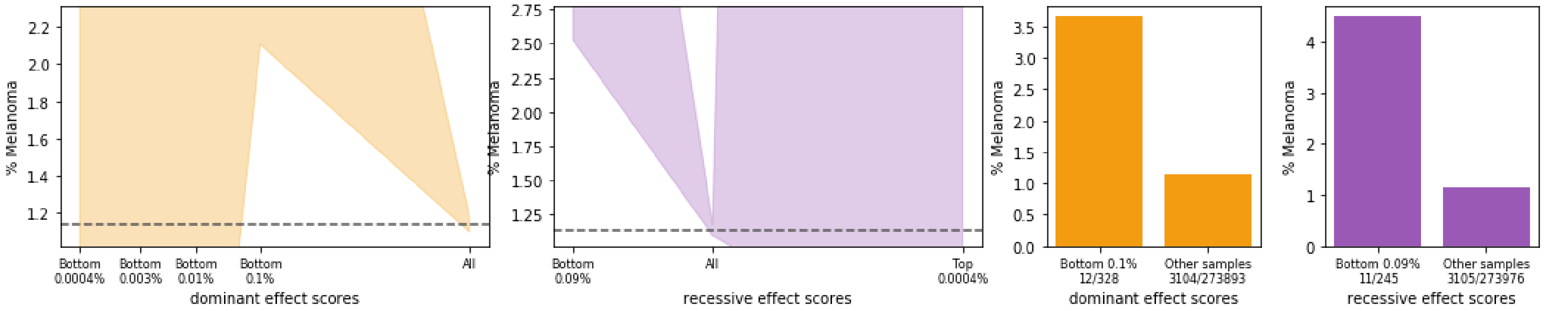

SLC12A9 & Melanoma: PWAS q-value = 0.02 (Recessive)

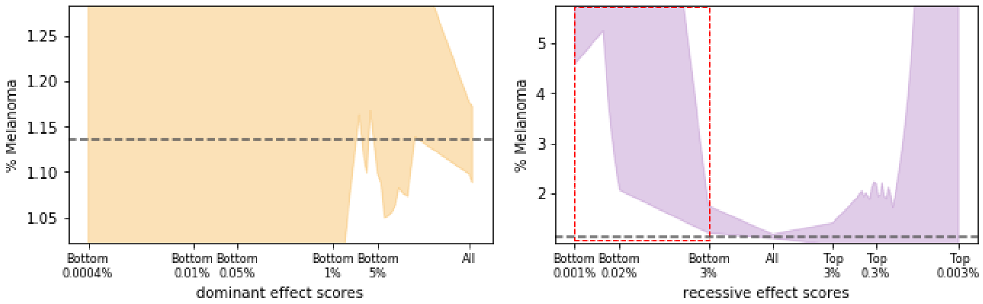

SLC45A2 & Melanoma: PWAS q-value = 8e-05 (Recessive)

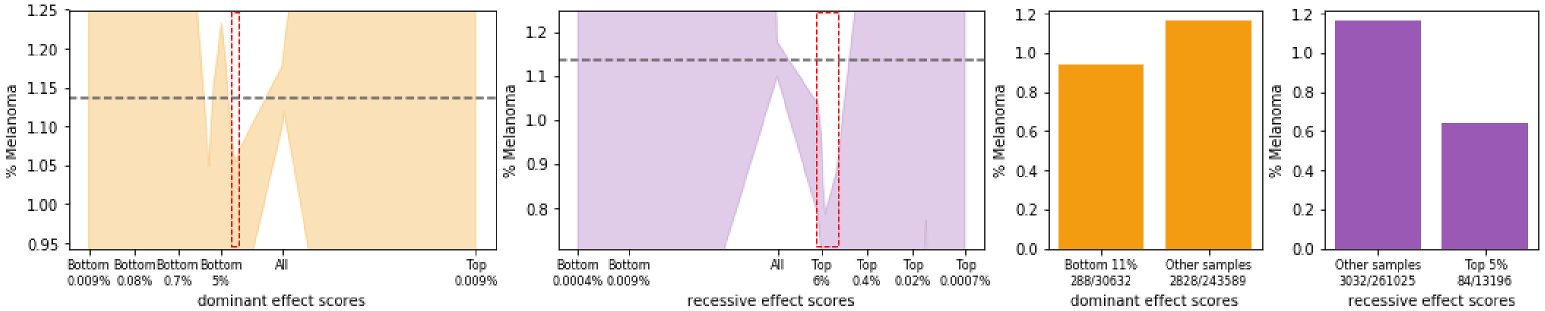

STN1 & Melanoma: PWAS q-value = 0.005 (Recessive)

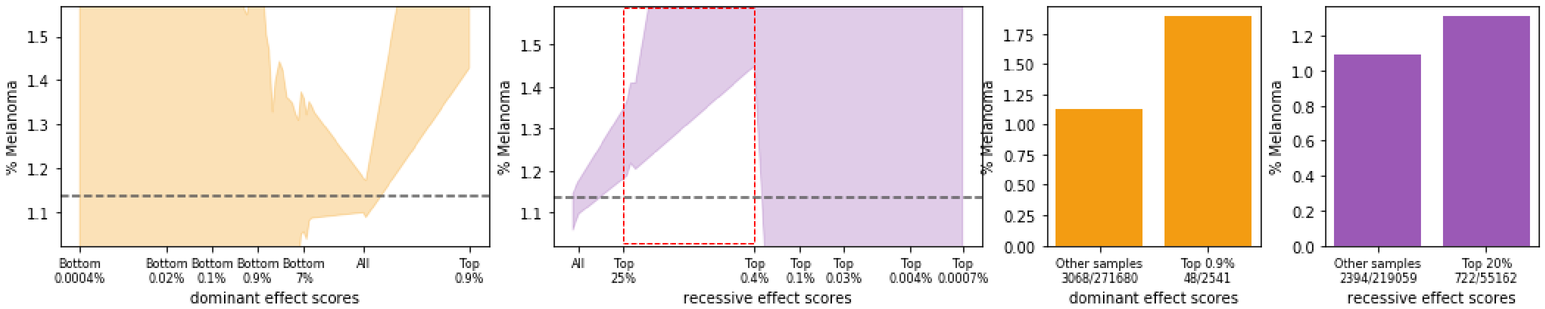

STX8 & Melanoma: PWAS q-value = 0.003 (Recessive)

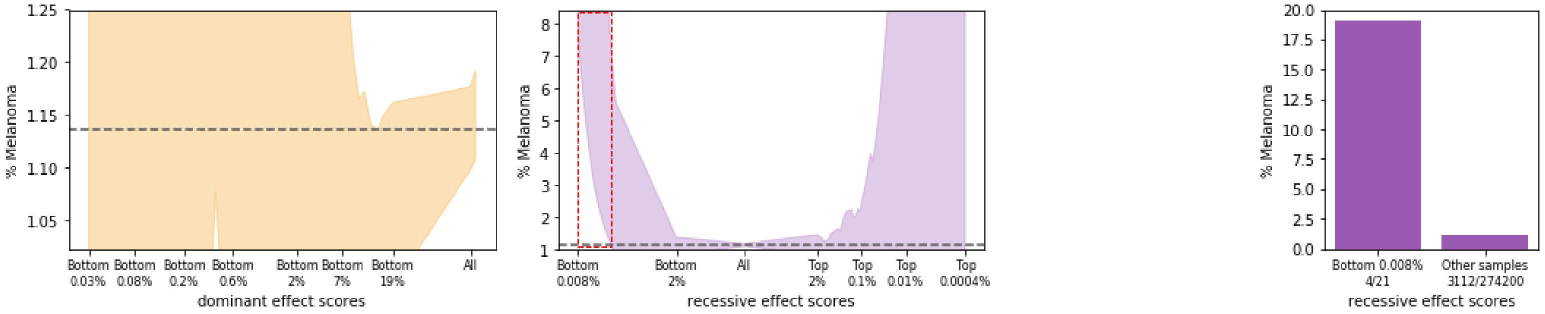

TYR & Melanoma: PWAS q-value = 0.005 (Recessive)

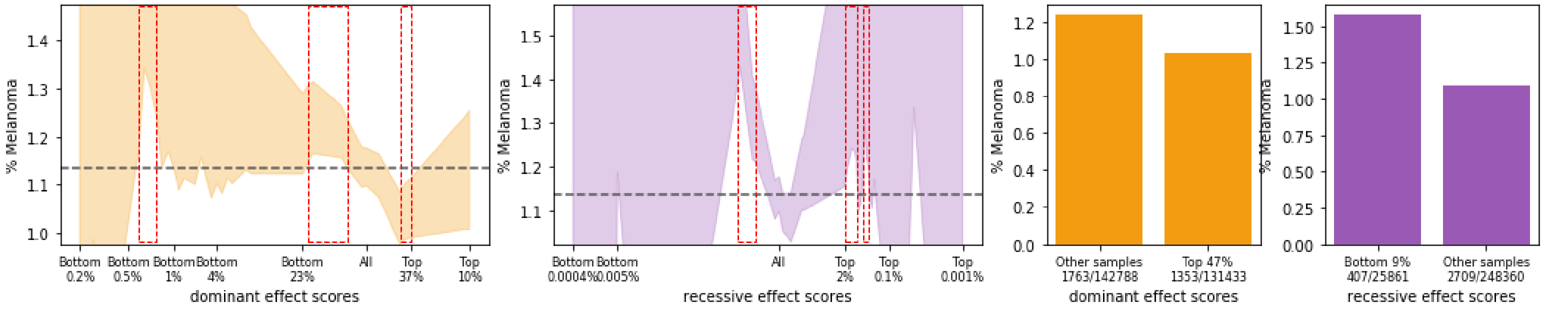

ZNF276 & Melanoma: PWAS q-value = 0.02 (Recessive)

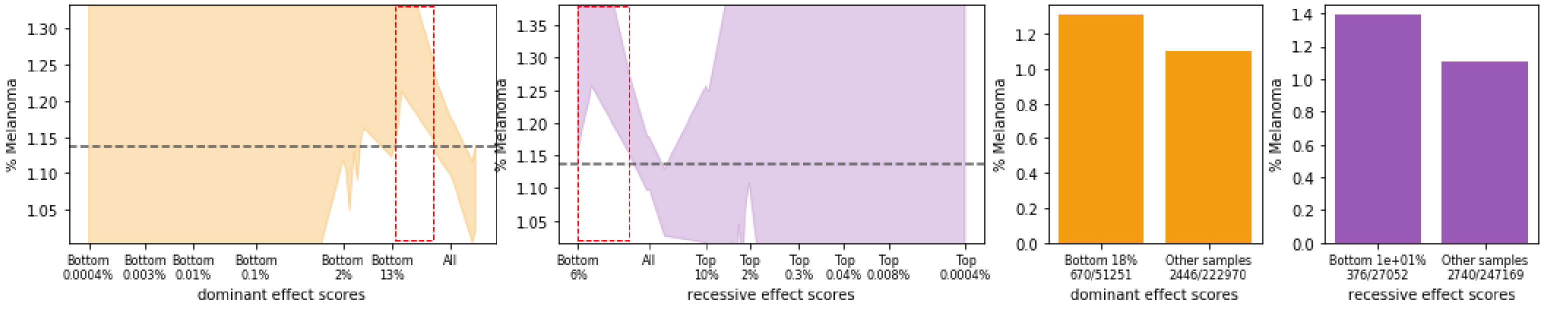

ALS2CR12 & Pan-cancer: PWAS q-value = 5e-09 (Dominant & recessive)

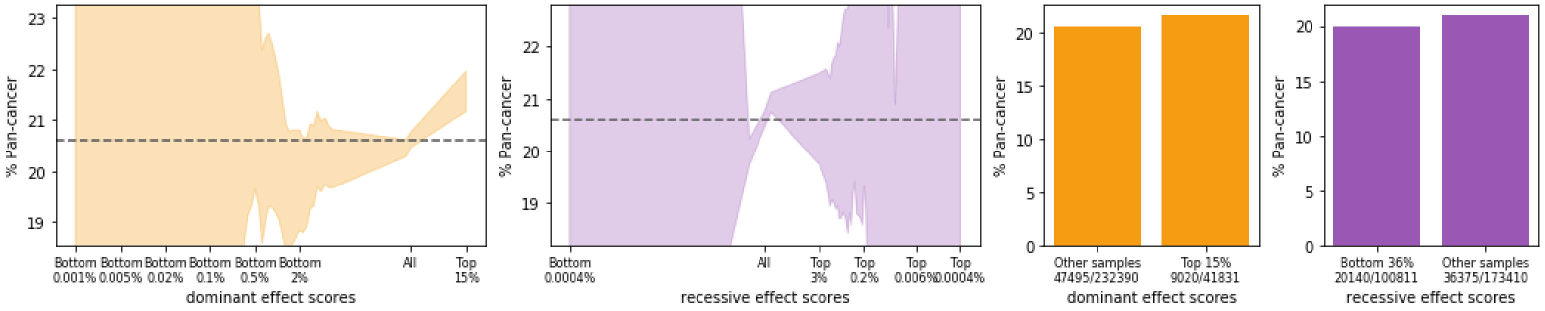

BNIPL & Pan-cancer: PWAS q-value = 0.01 (Dominant)

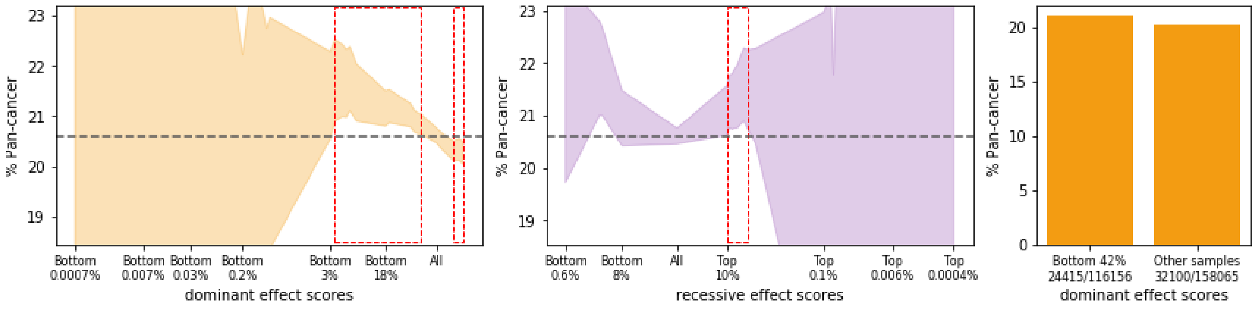

C1orf52 & Pan-cancer: PWAS q-value = 0.04 (Dominant)

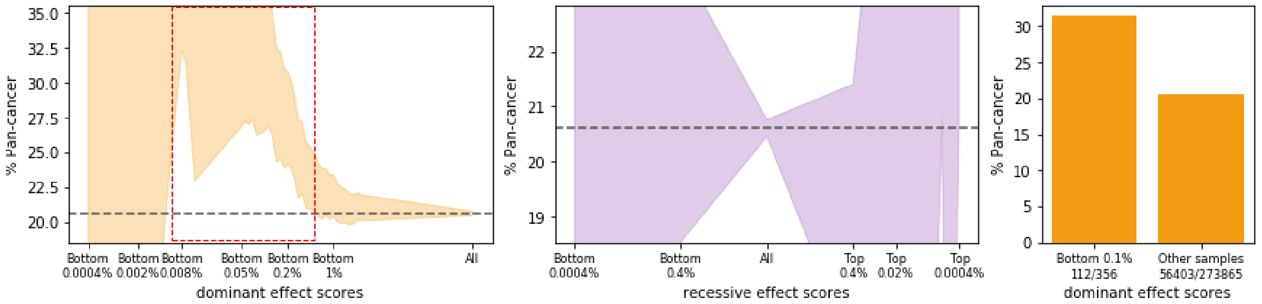

CERS2 & Pan-cancer: PWAS q-value = 0.006 (Dominant)

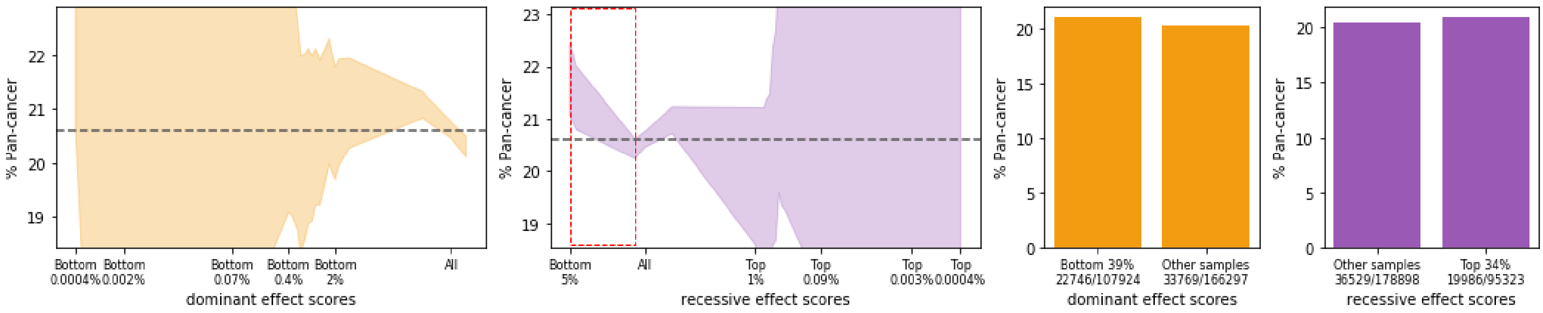

CHEK2 & Pan-cancer: PWAS q-value = 0.007 (Dominant)

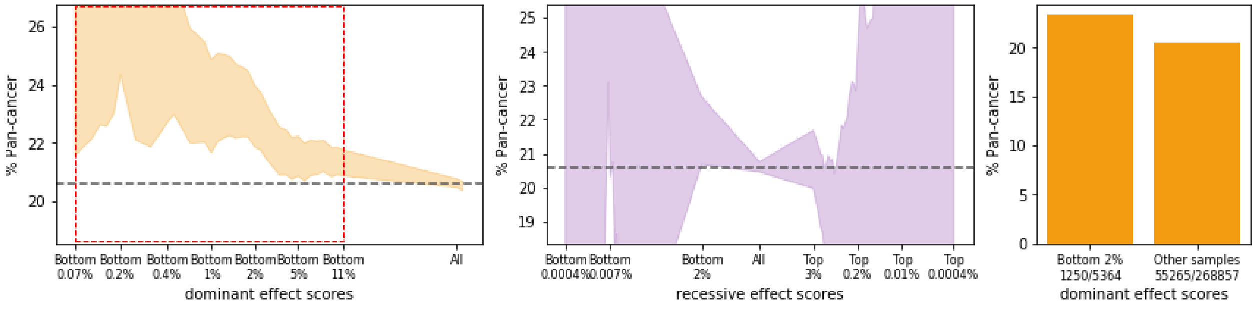

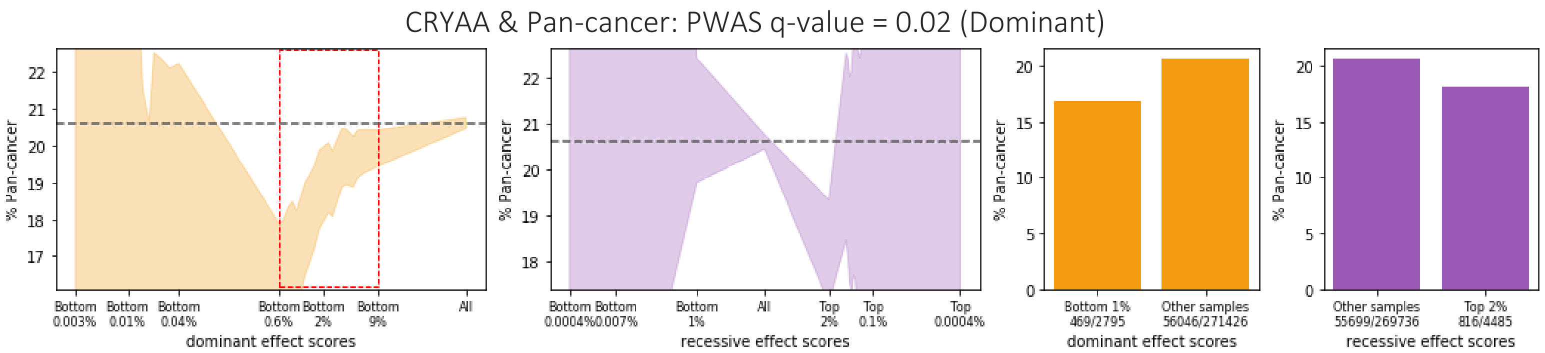
CRYAA & Pan-cancer: PWAS q-value = 0.02 (Dominant)
% Pan-cancer
Bottom 0.003%
Bottom 0.01%
Bottom 0.04%
Bottom 0.6%
Bottom 2%
Bottom 9%
All
dominant effect scores
Bottom 0.0004%
Bottom 0.007%
Bottom 1%
All
Top 2%
Top 0.1%
Top 0.0004%
recessive effect scores
Bottom 1% 469/2795
Other samples 56046/271426
dominant effect scores
Other samples 55699/269736
Top 2% 816/4485
recessive effect scores

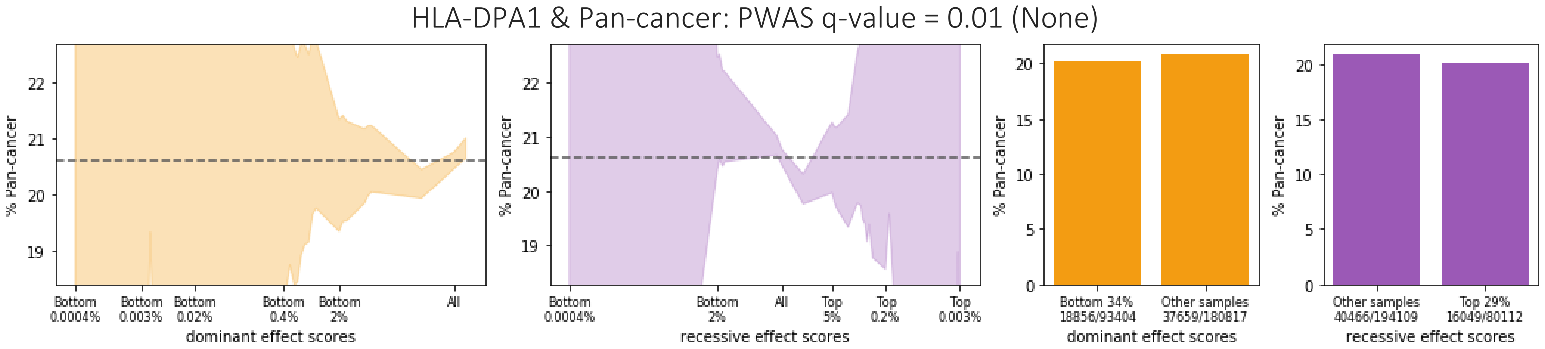
HLA-DPA1 & Pan-cancer: PWAS q-value = 0.01 (None)
% Pan-cancer
Bottom 0.0004%
Bottom 0.003%
Bottom 0.02%
Bottom 0.4%
Bottom 2%
All
dominant effect scores
Bottom 0.0004%
Bottom 2%
All
Top 5%
Top 0.2%
Top 0.003%
recessive effect scores
Bottom 34% 18856/93404
Other samples 37659/180817
dominant effect scores
Other samples 40466/194109
Top 29% 16049/80112
recessive effect scores

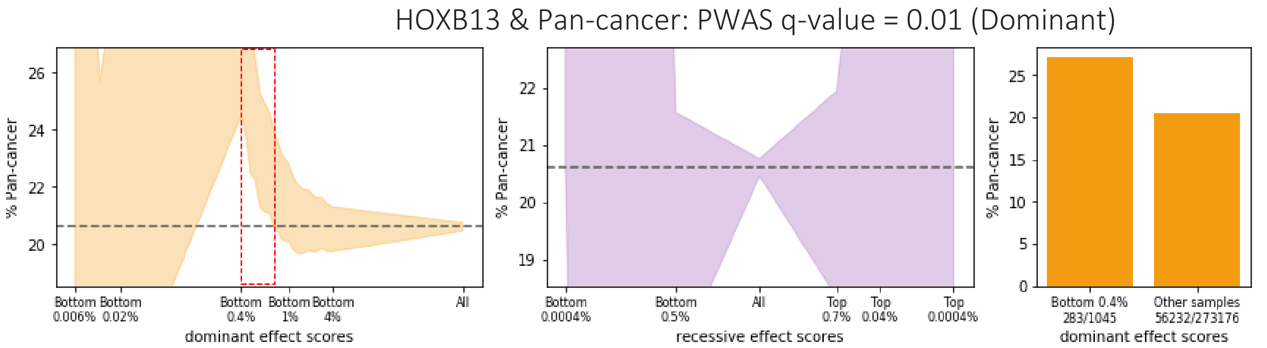
HOXB13 & Pan-cancer: PWAS q-value = 0.01 (Dominant)
% Pan-cancer
Bottom 0.006%
Bottom 0.02%
Bottom 0.4%
Bottom 1%
Bottom 4%
All
dominant effect scores
Bottom 0.0004%
Bottom 0.5%
All
Top 0.7%
Top 0.04%
Top 0.0004%
recessive effect scores
Bottom 0.4% 283/1045
Other samples 56232/273176
dominant effect scores

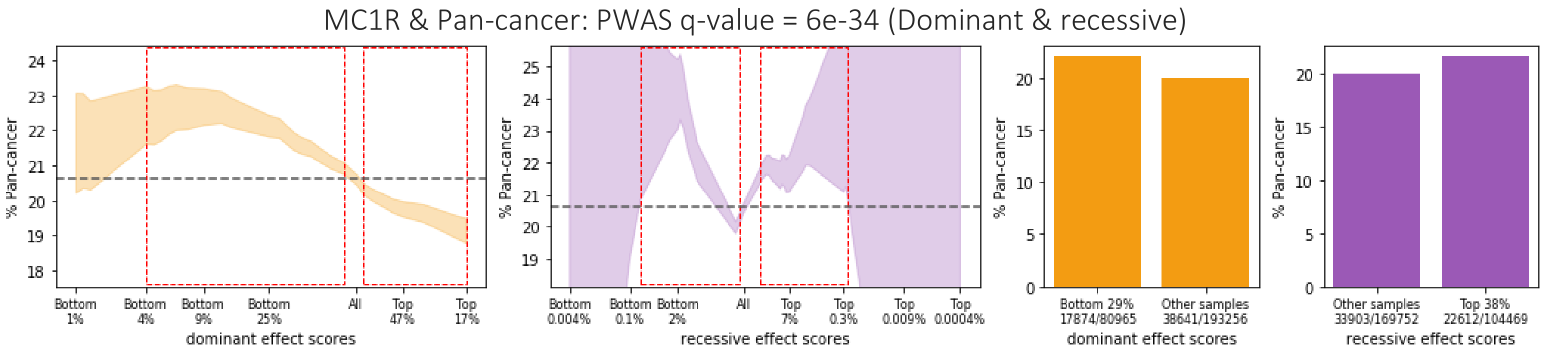
MC1R & Pan-cancer: PWAS q-value = 6e-34 (Dominant & recessive)
% Pan-cancer
Bottom 1%
Bottom 4%
Bottom 9%
Bottom 25%
All
Top 47%
Top 17%
dominant effect scores
Bottom 0.004%
Bottom 0.1%
Bottom 2%
All
Top 7%
Top 0.3%
Top 0.009%
Top 0.0004%
recessive effect scores
Bottom 29% 17874/80965
Other samples 38641/193256
dominant effect scores
Other samples 33903/169752
Top 38% 22612/104469
recessive effect scores

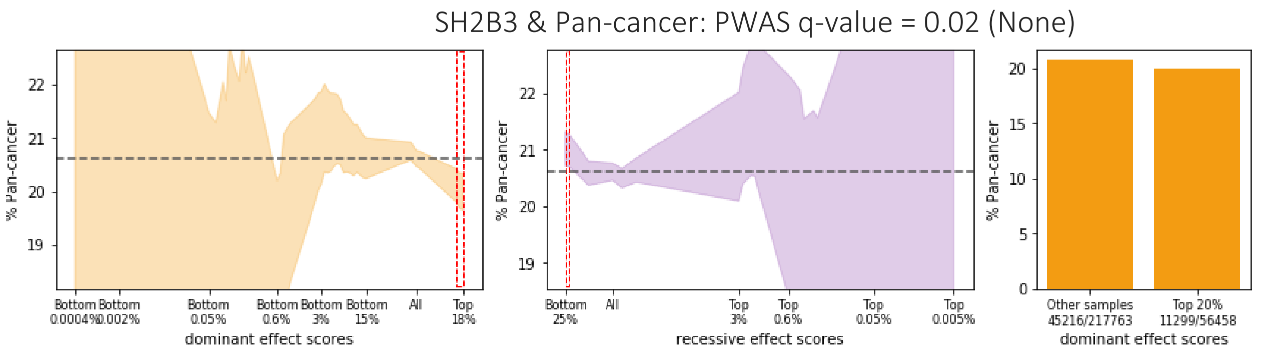
SH2B3 & Pan-cancer: PWAS q-value = 0.02 (None)
% Pan-cancer
dominant effect scores
recessive effect scores
dominant effect scores

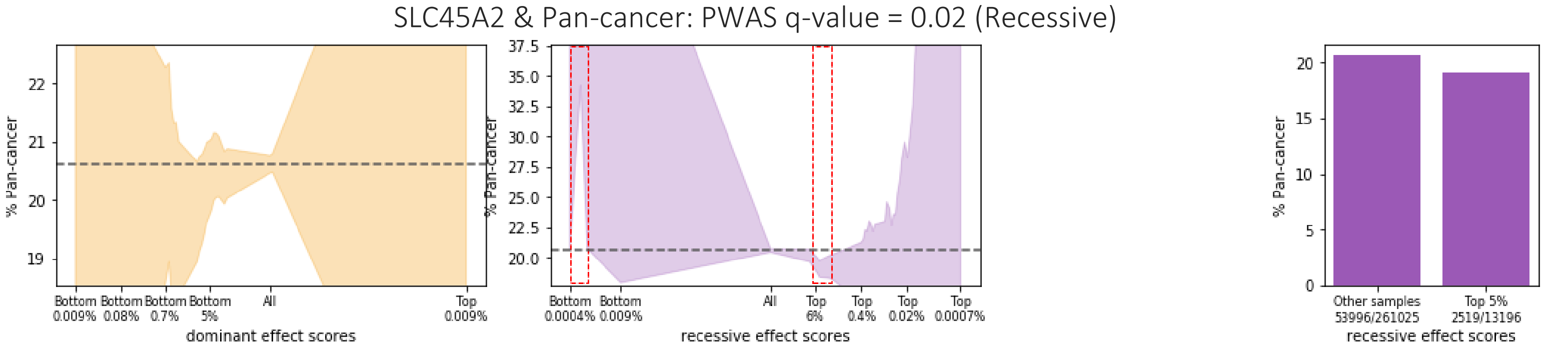
SLC45A2 & Pan-cancer: PWAS q-value = 0.02 (Recessive)
% Pan-cancer
dominant effect scores
recessive effect scores
recessive effect scores

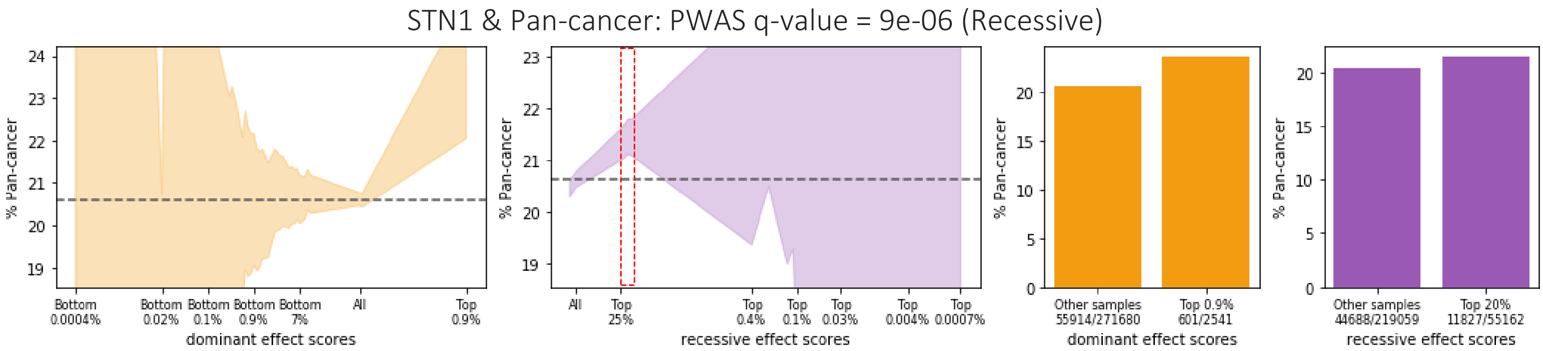
STN1 & Pan-cancer: PWAS q-value = 9e-06 (Recessive)
% Pan-cancer
dominant effect scores
recessive effect scores
dominant effect scores
recessive effect scores

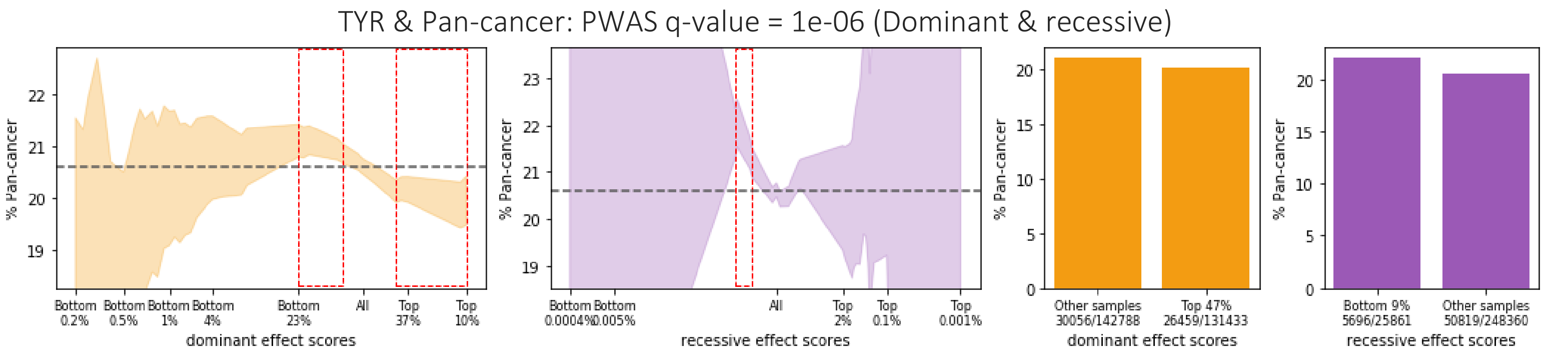
TYR & Pan-cancer: PWAS q-value = 1e-06 (Dominant & recessive)
% Pan-cancer
dominant effect scores
recessive effect scores
dominant effect scores
recessive effect scores

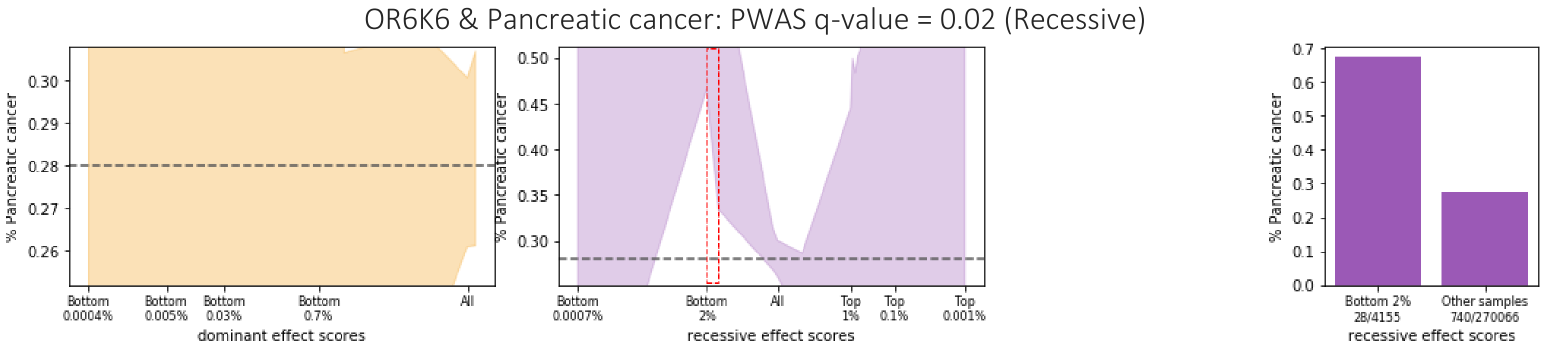
OR6K6 & Pancreatic cancer: PWAS q-value = 0.02 (Recessive)
% Pancreatic cancer
dominant effect scores
Bottom 0.0004%
Bottom 0.005%
Bottom 0.03%
Bottom 0.7%
All
recessive effect scores
Bottom 0.0007%
Bottom 2%
All
Top 1%
Top 0.1%
Top 0.001%
Bottom 2% 28/4155
Other samples 740/270066
recessive effect scores

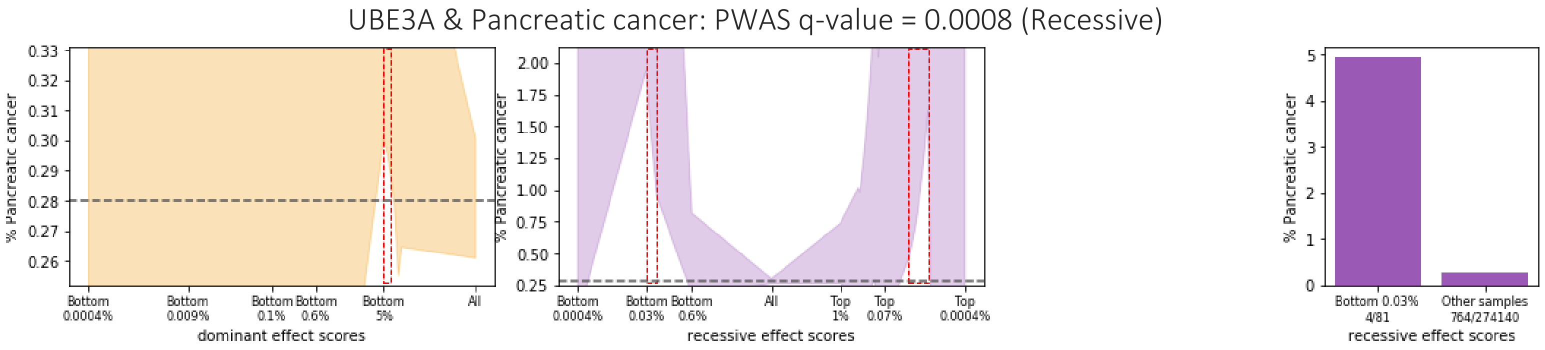
UBE3A & Pancreatic cancer: PWAS q-value = 0.0008 (Recessive)
% Pancreatic cancer
dominant effect scores
Bottom 0.0004%
Bottom 0.009%
Bottom 0.1%
Bottom 0.6%
Bottom 5%
All
recessive effect scores
Bottom 0.0004%
Bottom 0.03%
Bottom 0.6%
All
Top 1%
Top 0.07%
Top 0.0004%
Bottom 0.03% 4/81
Other samples 764/274140
recessive effect scores

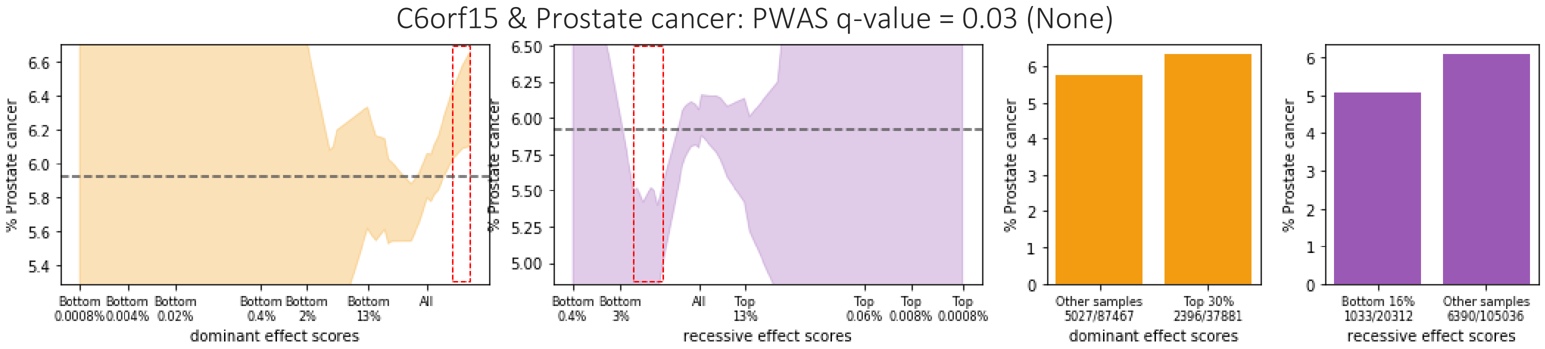
C6orf15 & Prostate cancer: PWAS q-value = 0.03 (None)
% Prostate cancer
dominant effect scores
Bottom 0.0008%
Bottom 0.004%
Bottom 0.02%
Bottom 0.4%
Bottom 2%
Bottom 13%
All
recessive effect scores
Bottom 0.4%
Bottom 3%
All
Top 13%
Top 0.06%
Top 0.008%
Top 0.0008%
Other samples 5027/87467
Top 30% 2396/37881
dominant effect scores
Bottom 16% 1033/20312
Other samples 6390/105036
recessive effect scores

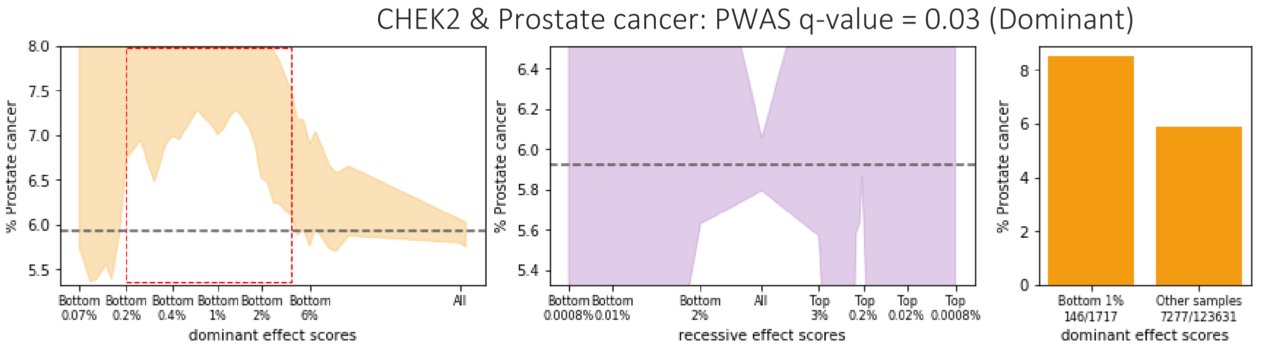
CHEK2 & Prostate cancer: PWAS q-value = 0.03 (Dominant)
% Prostate cancer
dominant effect scores
Bottom 0.07%
Bottom 0.2%
Bottom 0.4%
Bottom 1%
Bottom 2%
Bottom 6%
All
recessive effect scores
Bottom 0.0008%
Bottom 0.01%
Bottom 2%
All
Top 3%
Top 0.2%
Top 0.02%
Top 0.0008%
Bottom 1% 146/1717
Other samples 7277/123631
dominant effect scores

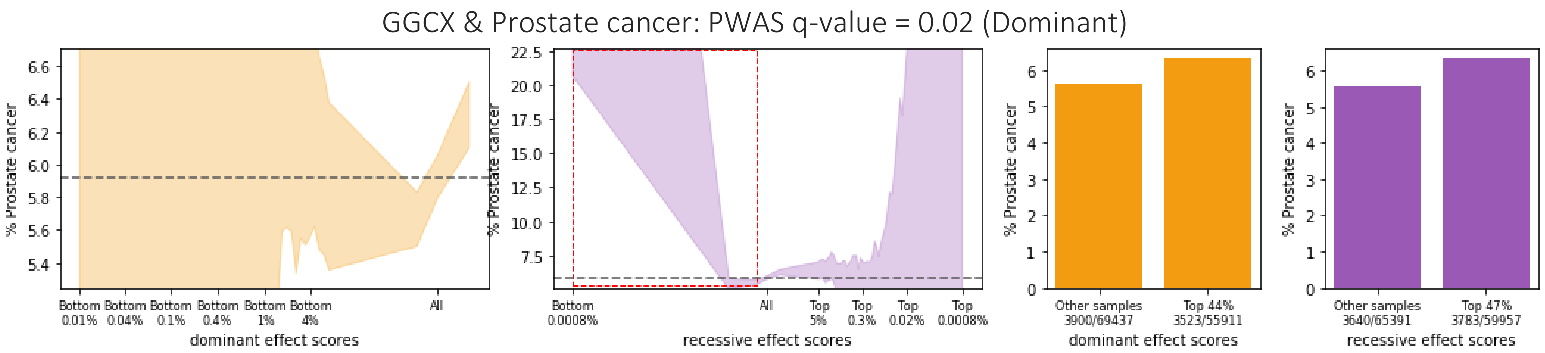
GGCX & Prostate cancer: PWAS q-value = 0.02 (Dominant)
% Prostate cancer
dominant effect scores
recessive effect scores
Other samples 3900/69437
Top 44% 3523/55911
Other samples 3640/65391
Top 47% 3783/59957

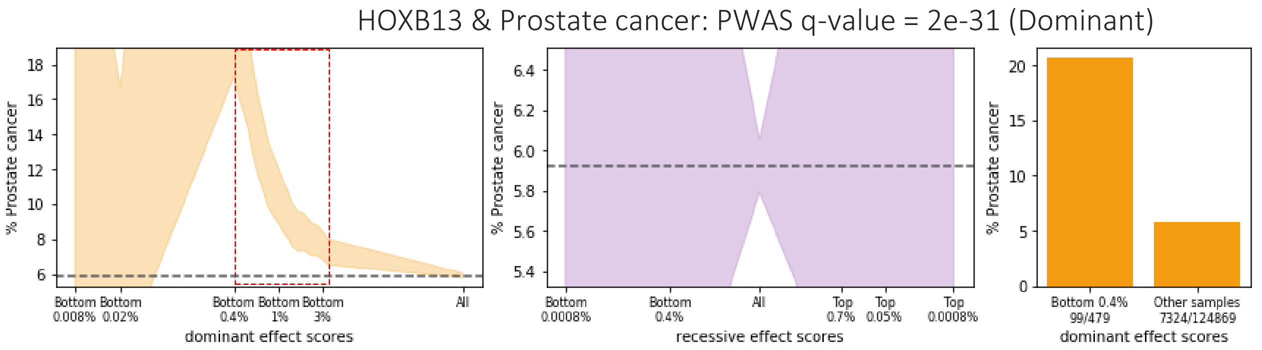
HOXB13 & Prostate cancer: PWAS q-value = 2e-31 (Dominant)
% Prostate cancer
dominant effect scores
recessive effect scores
Bottom 0.4% 99/479
Other samples 7324/124869

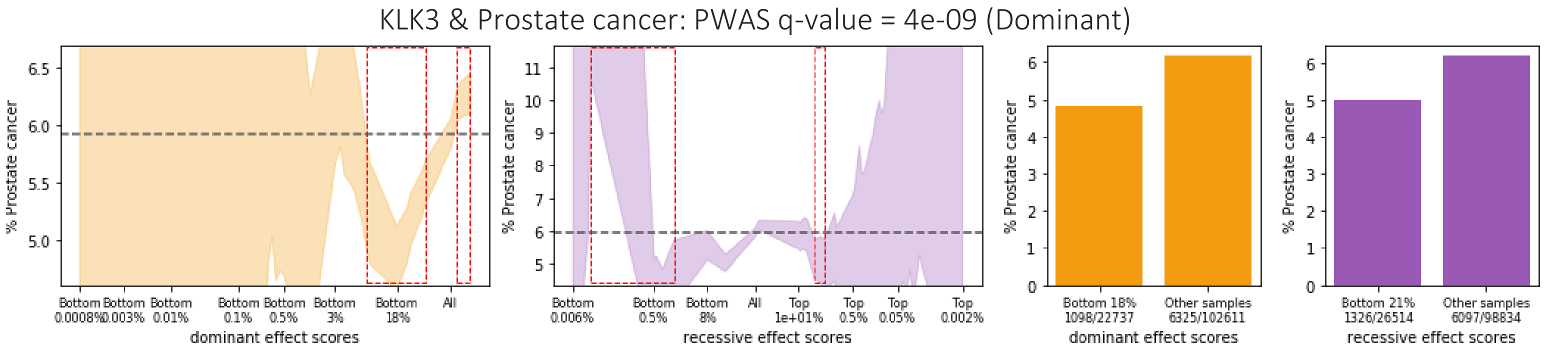
KLK3 & Prostate cancer: PWAS q-value = 4e-09 (Dominant)
% Prostate cancer
dominant effect scores
recessive effect scores
Bottom 18% 1098/22737
Other samples 6325/102611
Bottom 21% 1326/26514
Other samples 6097/98834

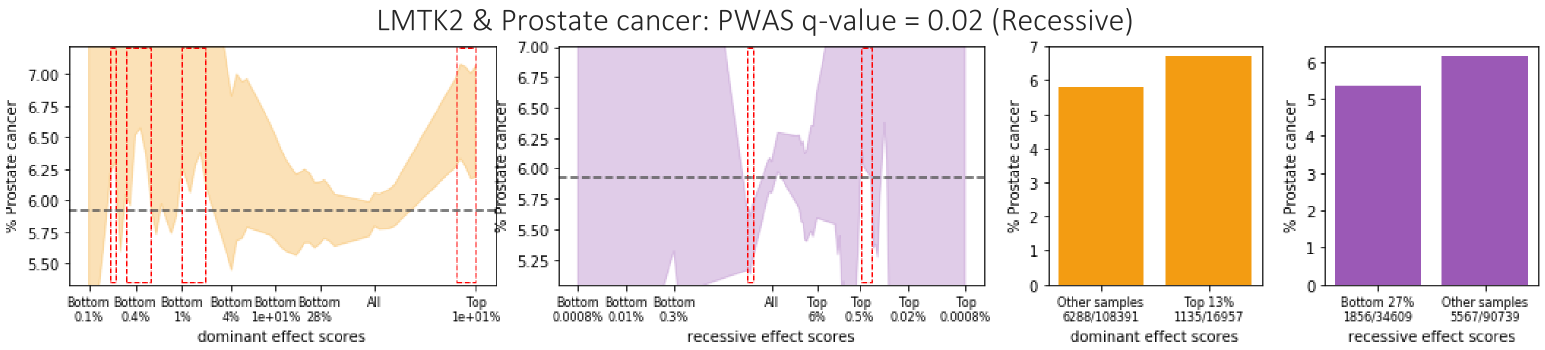
LMTK2 & Prostate cancer: PWAS q-value = 0.02 (Recessive)
% Prostate cancer
dominant effect scores
recessive effect scores
Other samples 6288/108391
Top 13% 1135/16957
Bottom 27% 1856/34609
Other samples 5567/90739

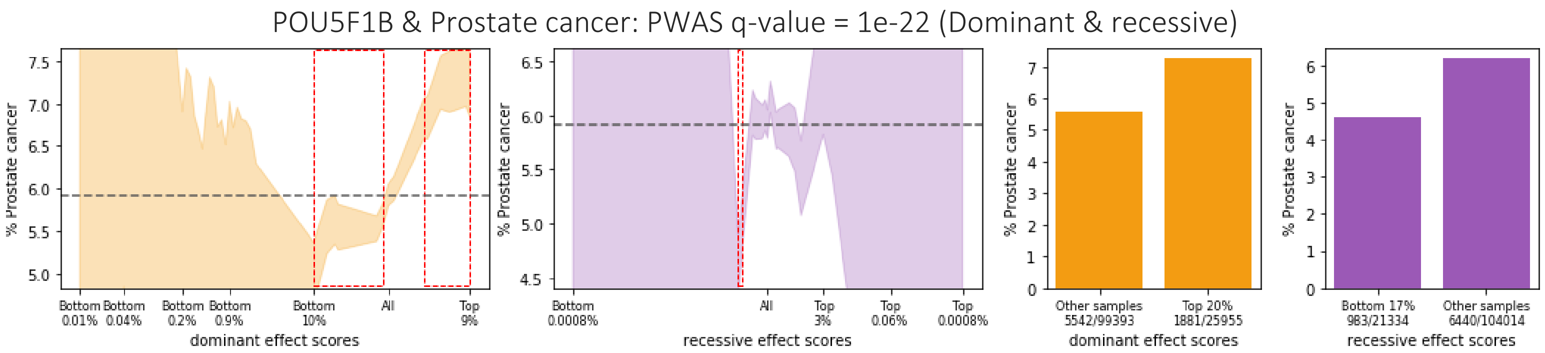
POU5F1B & Prostate cancer: PWAS q-value = 1e-22 (Dominant & recessive)
% Prostate cancer
Bottom 0.01%
Bottom 0.04%
Bottom 0.2%
Bottom 0.9%
Bottom 10%
All
Top 9%
dominant effect scores
Bottom 0.0008%
All
Top 3%
Top 0.06%
Top 0.0008%
recessive effect scores
Other samples 5542/99393
Top 20% 1881/25955
dominant effect scores
Bottom 17% 983/21334
Other samples 6440/104014
recessive effect scores

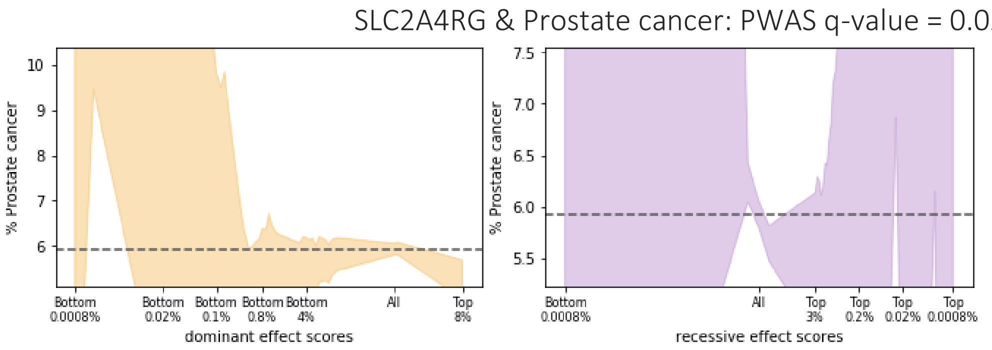
SLC2A4RG & Prostate cancer: PWAS q-value = 0.02 (Recessive)
% Prostate cancer
Bottom 0.0008%
Bottom 0.02%
Bottom 0.1%
Bottom 0.8%
Bottom 4%
All
Top 8%
dominant effect scores
Bottom 0.0008%
All
Top 3%
Top 0.2%
Top 0.02%
Top 0.0008%
recessive effect scores

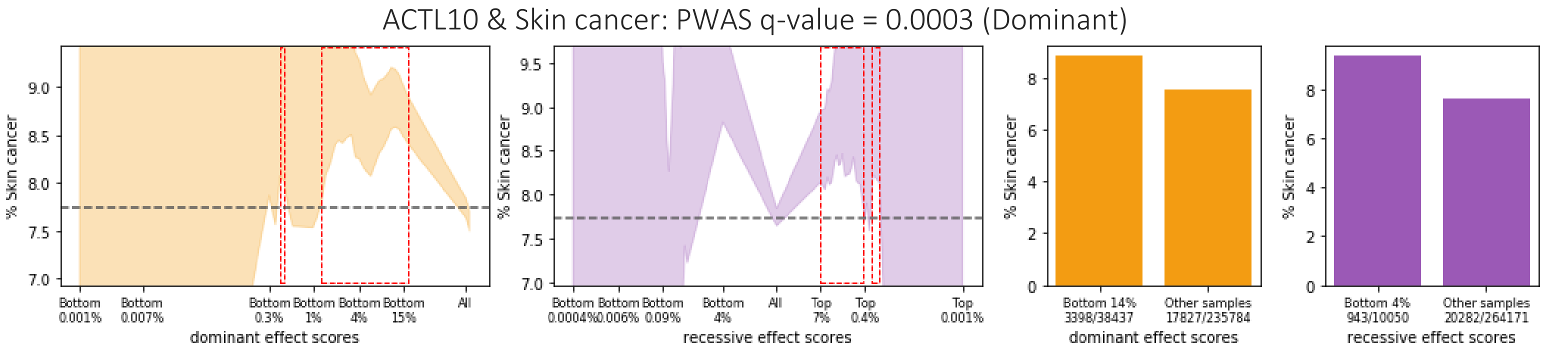
ACTL10 & Skin cancer: PWAS q-value = 0.0003 (Dominant)
% Skin cancer
Bottom 0.001%
Bottom 0.007%
Bottom 0.3%
Bottom 1%
Bottom 4%
Bottom 15%
All
dominant effect scores
Bottom 0.0004%
Bottom 0.006%
Bottom 0.09%
Bottom 4%
All
Top 7%
Top 0.4%
Top 0.001%
recessive effect scores
Bottom 14% 3398/38437
Other samples 17827/235784
dominant effect scores
Bottom 4% 943/10050
Other samples 20282/264171
recessive effect scores

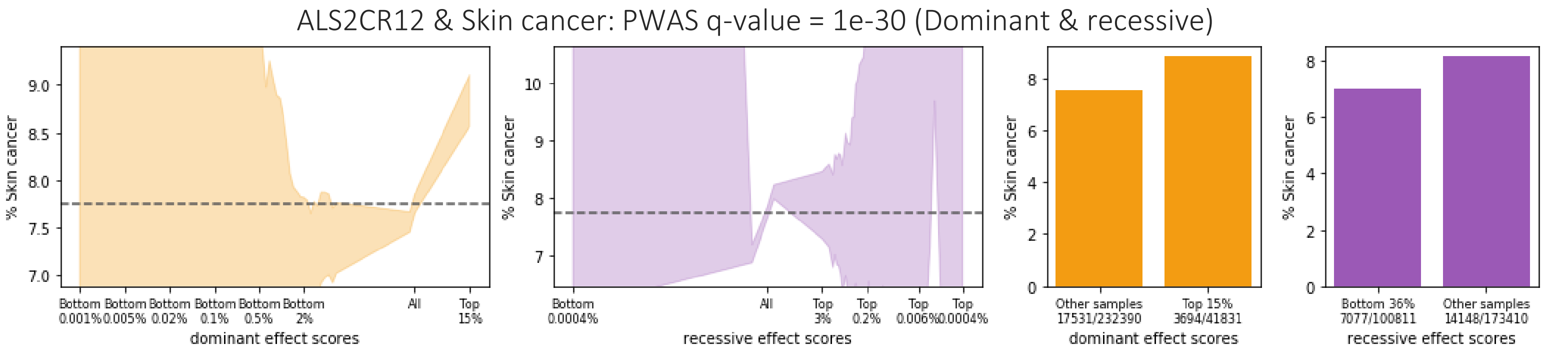
ALS2CR12 & Skin cancer: PWAS q-value = 1e-30 (Dominant & recessive)
% Skin cancer
Bottom 0.001%
Bottom 0.005%
Bottom 0.02%
Bottom 0.1%
Bottom 0.5%
Bottom 2%
All
Top 15%
dominant effect scores
Bottom 0.0004%
All
Top 3%
Top 0.2%
Top 0.006%
Top 0.0004%
recessive effect scores
Other samples 17531/232390
Top 15% 3694/41831
dominant effect scores
Bottom 36% 7077/100811
Other samples 14148/173410
recessive effect scores

ATP8B4 & Skin cancer: PWAS q-value = 0.03 (Recessive)

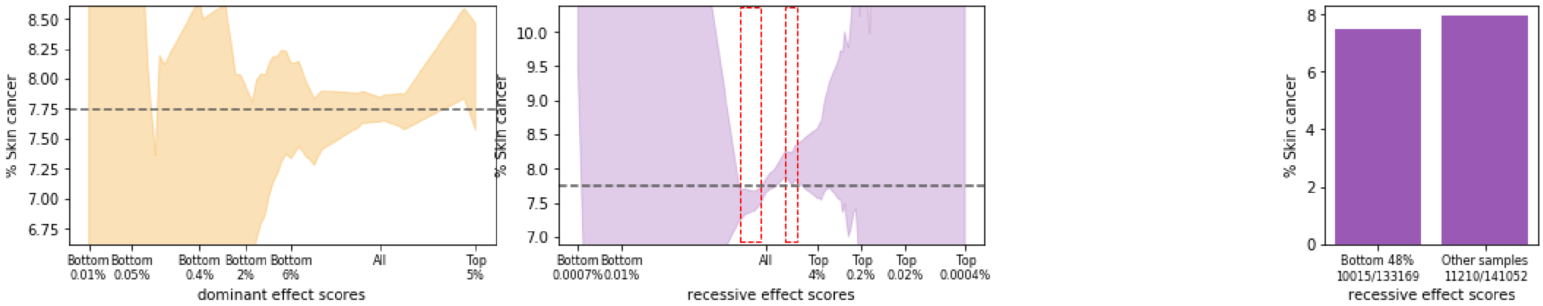

BCAS4 & Skin cancer: PWAS q-value = 0.03 (Dominant)

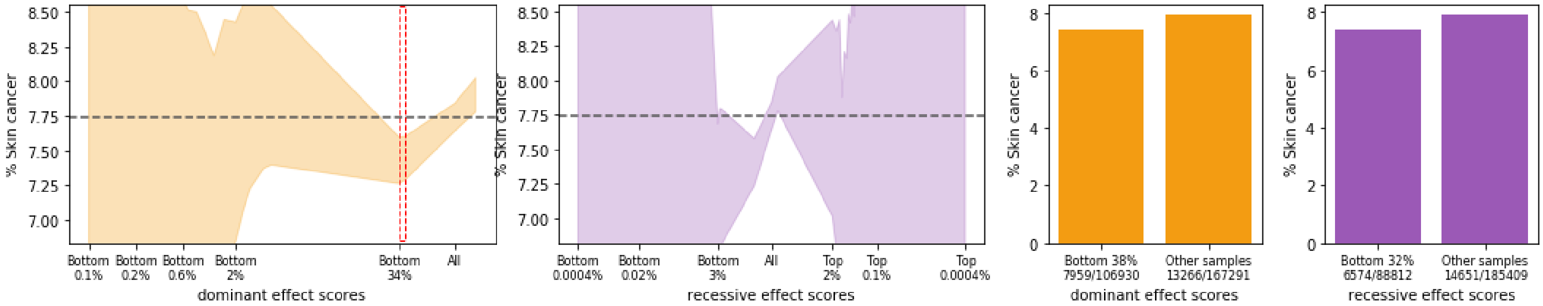

BNIPL & Skin cancer: PWAS q-value = 0.02 (None)

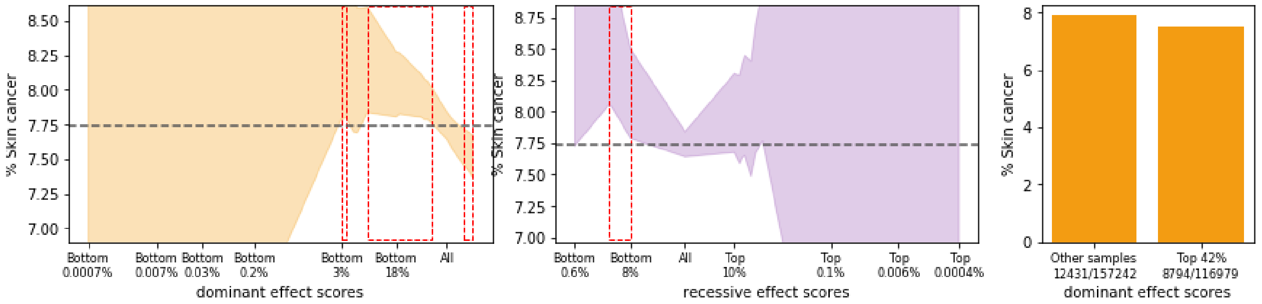

BPIFB3 & Skin cancer: PWAS q-value = 0.0005 (Dominant)

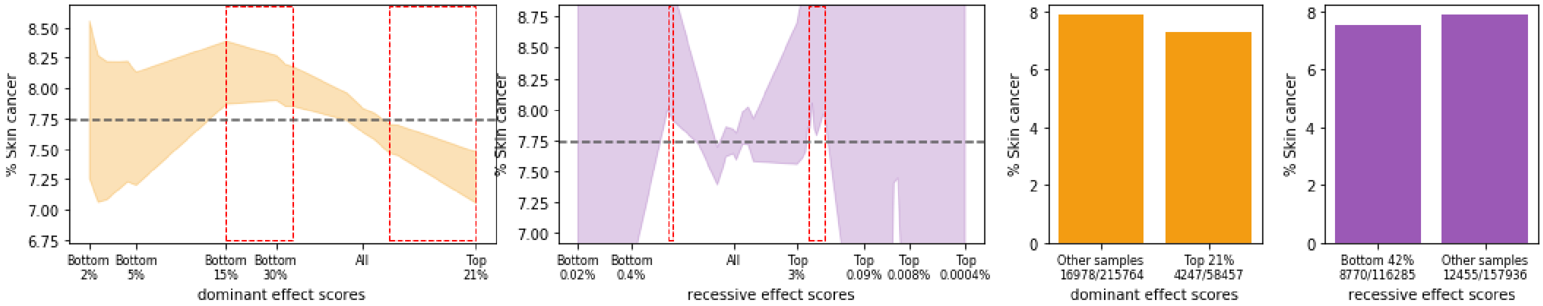

C6orf15 & Skin cancer: PWAS q-value = 0.02 (None)

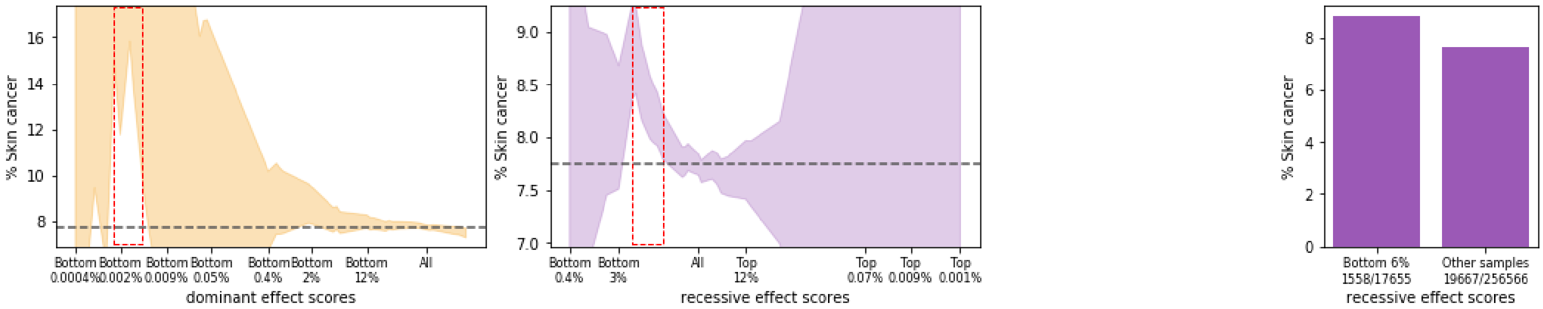

CCHCR1 & Skin cancer: PWAS q-value = 0.01 (Recessive)

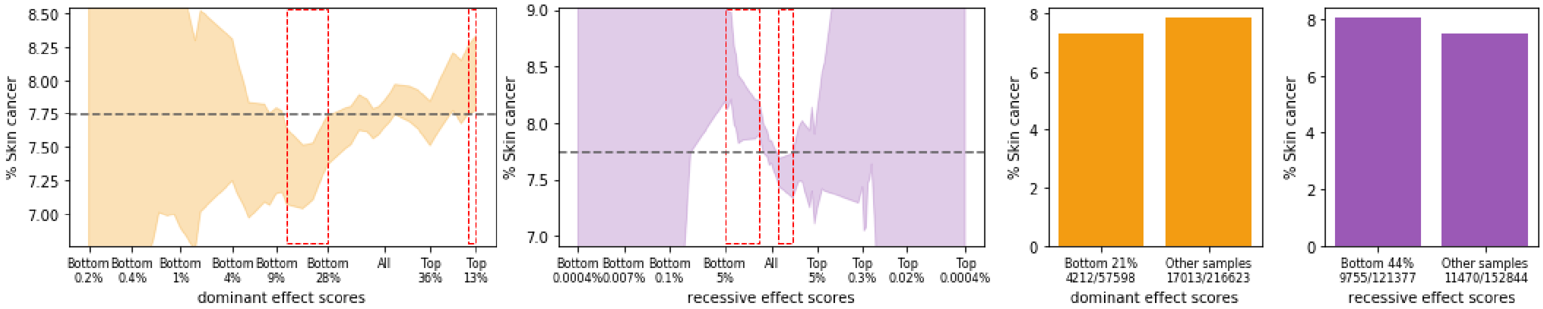

CDHR4 & Skin cancer: PWAS q-value = 0.03 (Dominant)

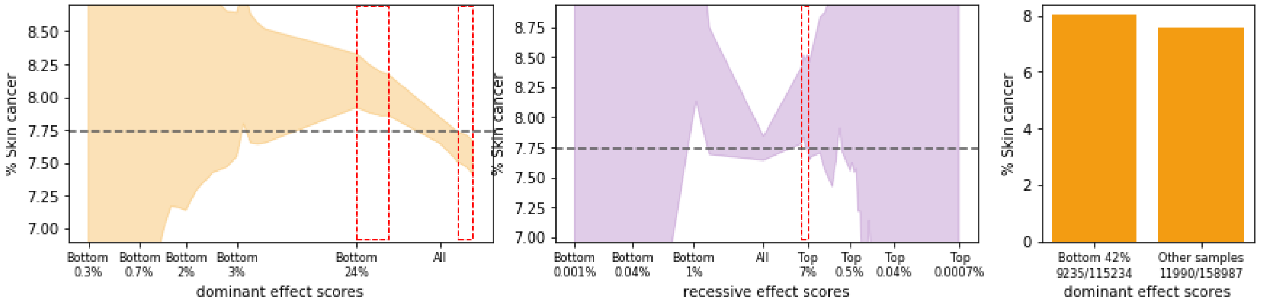

CTLA4 & Skin cancer: PWAS q-value = 4e-07 (Dominant & recessive)

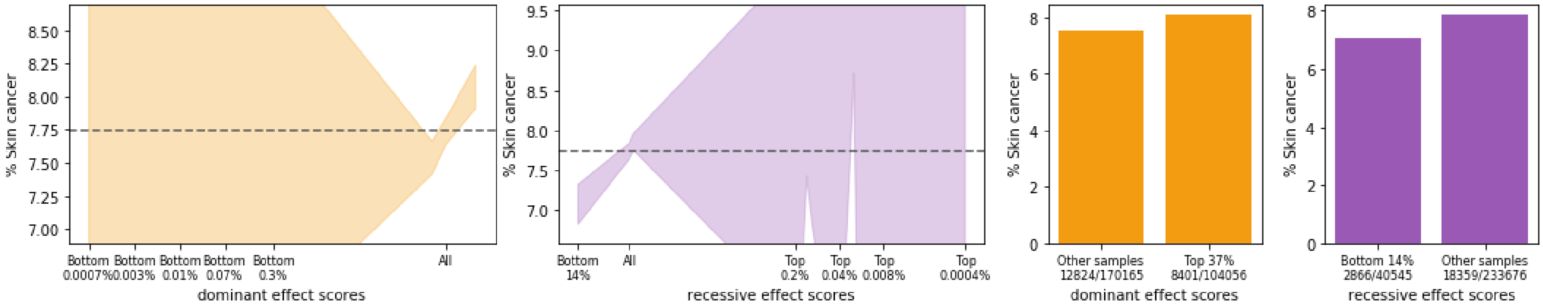

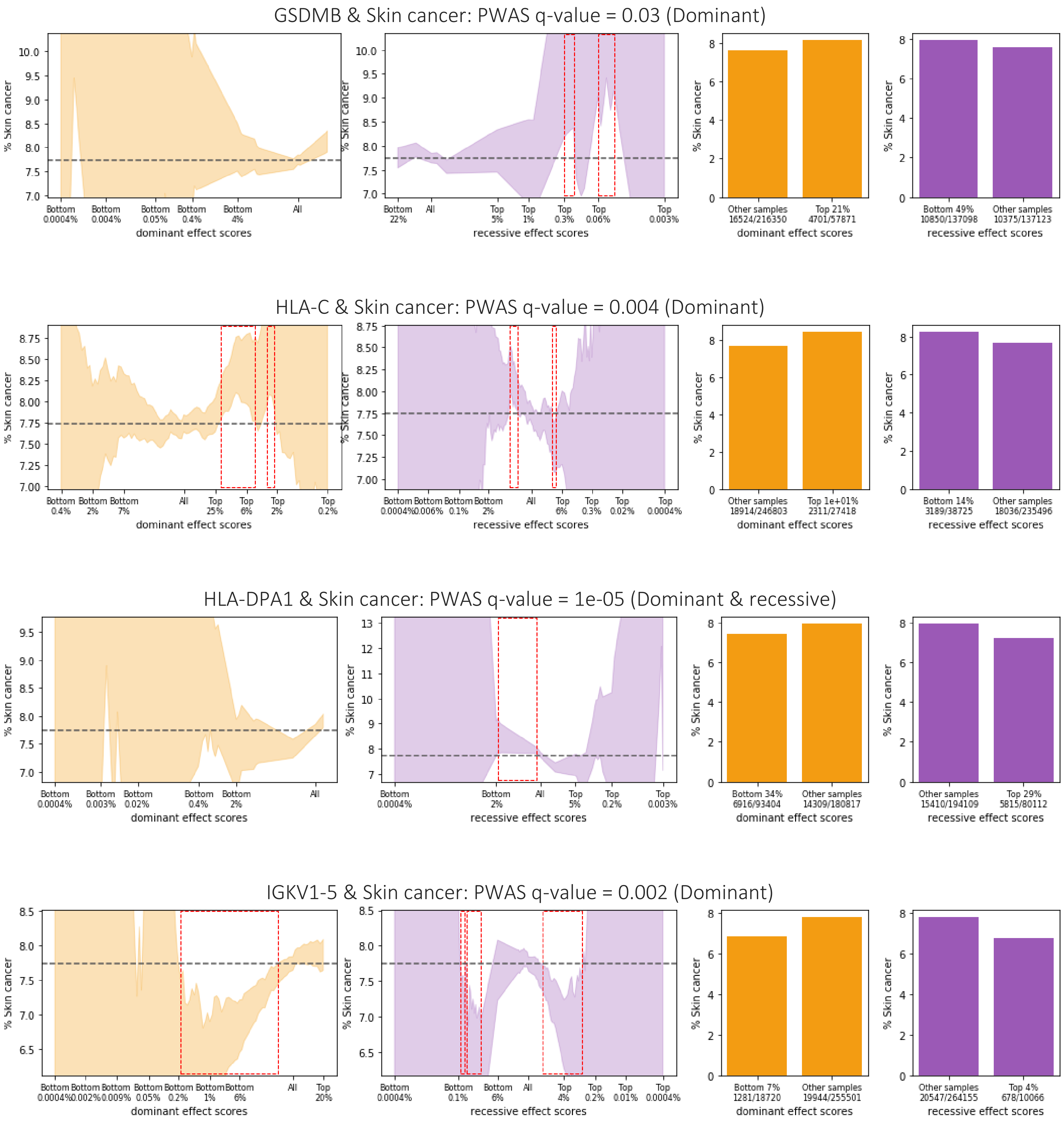
GSDMB & Skin cancer: PWAS q-value = 0.03 (Dominant)
% Skin cancer
dominant effect scores
recessive effect scores
Other samples 16524/216350
Top 21% 4701/57871
Bottom 49% 10850/137098
Other samples 10375/137123
HLA-C & Skin cancer: PWAS q-value = 0.004 (Dominant)
Other samples 18914/246803
Top 1e+01% 2311/27418
Bottom 14% 3189/38725
Other samples 18036/235496
HLA-DPA1 & Skin cancer: PWAS q-value = 1e-05 (Dominant & recessive)
Bottom 34% 6916/93404
Other samples 14309/180817
Other samples 15410/194109
Top 29% 5815/80112
IGKV1-5 & Skin cancer: PWAS q-value = 0.002 (Dominant)
Bottom 7% 1281/18720
Other samples 19944/255501
Other samples 20547/264155
Top 4% 678/10066

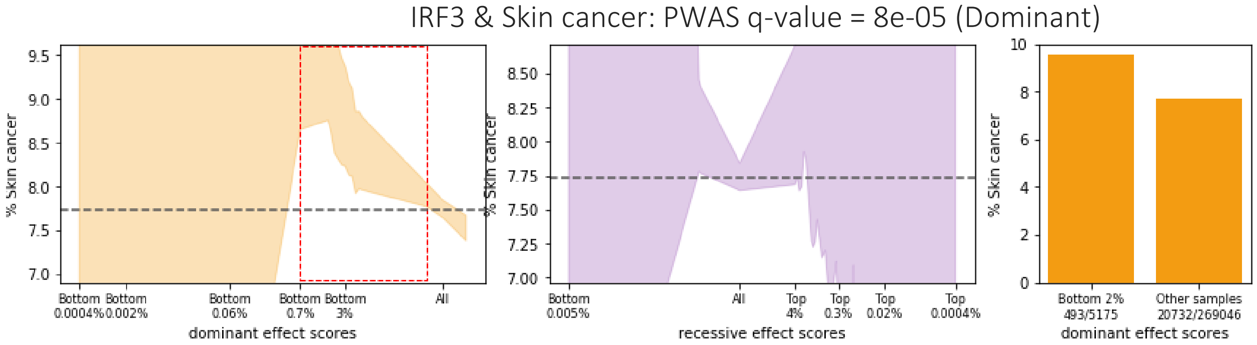

IRF3 & Skin cancer: PWAS q-value = 8e-05 (Dominant)
% Skin cancer
Bottom 0.0004%
Bottom 0.002%
Bottom 0.06%
Bottom 0.7%
Bottom 3%
All
dominant effect scores
Bottom 0.005%
All
Top 4%
Top 0.3%
Top 0.02%
Top 0.0004%
recessive effect scores
Bottom 2% 493/5175
Other samples 20732/269046
dominant effect scores

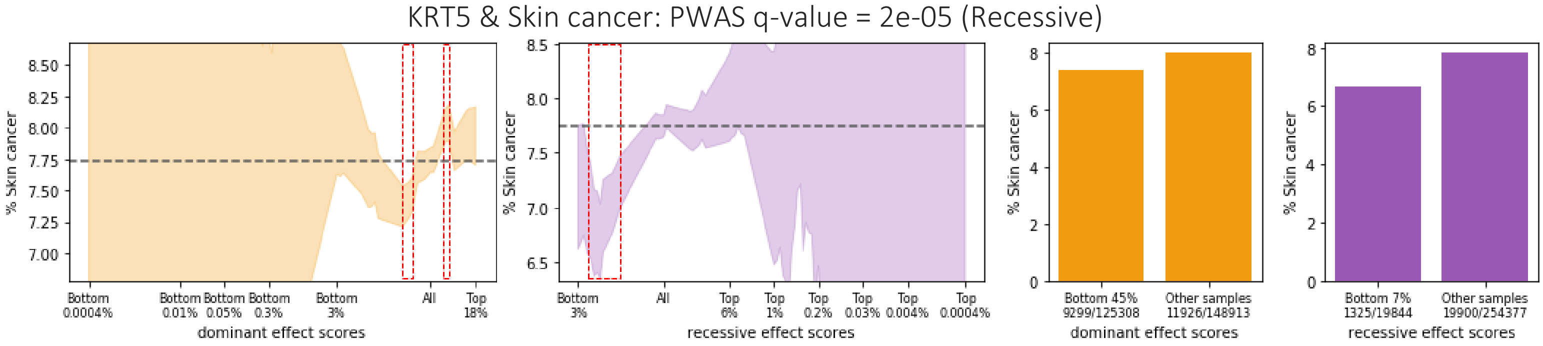

KRT5 & Skin cancer: PWAS q-value = 2e-05 (Recessive)
% Skin cancer
Bottom 0.0004%
Bottom 0.01%
Bottom 0.05%
Bottom 0.3%
Bottom 3%
All
Top 18%
dominant effect scores
Bottom 3%
All
Top 6%
Top 1%
Top 0.2%
Top 0.03%
Top 0.004%
Top 0.0004%
recessive effect scores
Bottom 45% 9299/125308
Other samples 11926/148913
dominant effect scores
Bottom 7% 1325/19844
Other samples 19900/254377
recessive effect scores

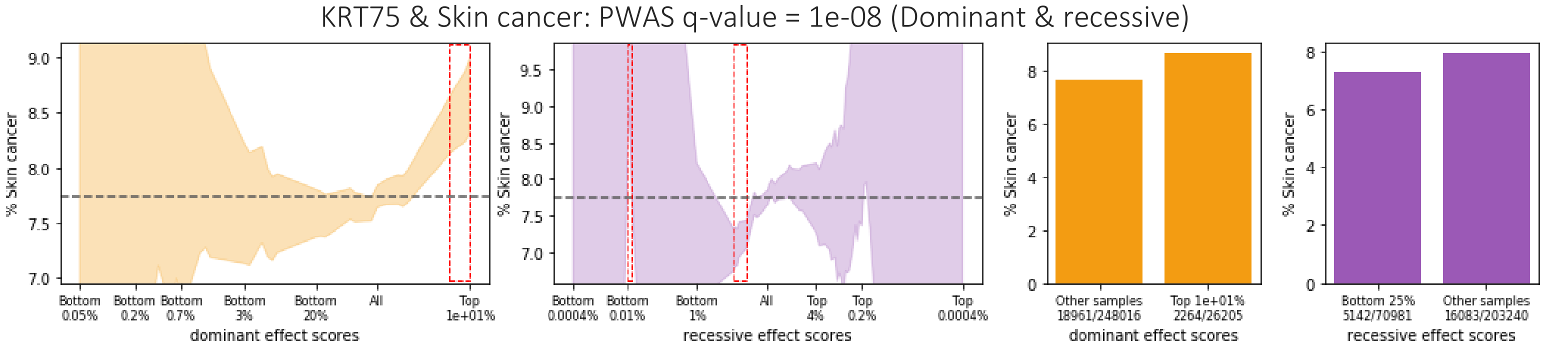

KRT75 & Skin cancer: PWAS q-value = 1e-08 (Dominant & recessive)
% Skin cancer
Bottom 0.05%
Bottom 0.2%
Bottom 0.7%
Bottom 3%
Bottom 20%
All
Top 1e+01%
dominant effect scores
Bottom 0.0004%
Bottom 0.01%
Bottom 1%
All
Top 4%
Top 0.2%
Top 0.0004%
recessive effect scores
Other samples 18961/248016
Top 1e+01% 2264/26205
dominant effect scores
Bottom 25% 5142/70981
Other samples 16083/203240
recessive effect scores

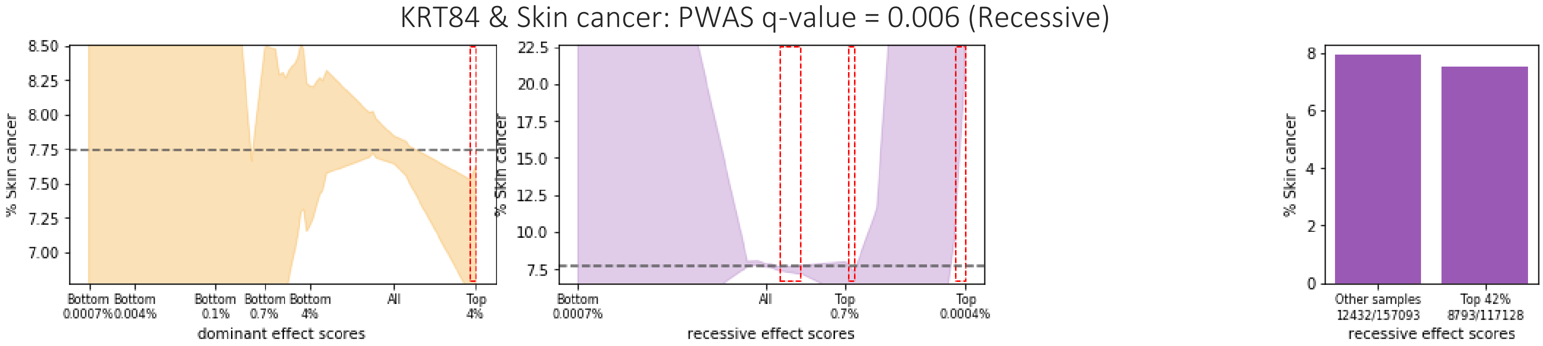

KRT84 & Skin cancer: PWAS q-value = 0.006 (Recessive)
% Skin cancer
Bottom 0.0007%
Bottom 0.004%
Bottom 0.1%
Bottom 0.7%
Bottom 4%
All
Top 4%
dominant effect scores
Bottom 0.0007%
All
Top 0.7%
Top 0.0004%
recessive effect scores
Other samples 12432/157093
Top 42% 8793/117128
recessive effect scores

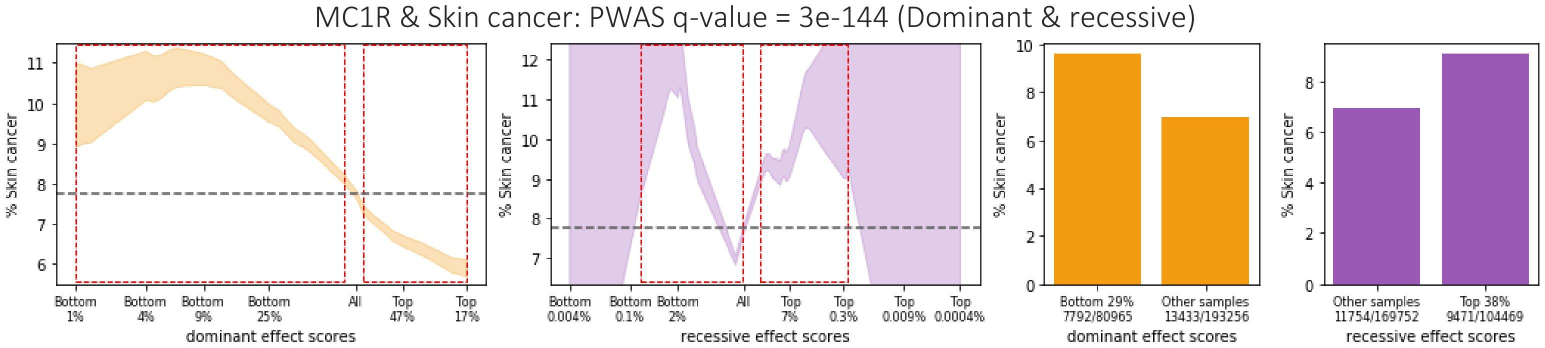
MC1R & Skin cancer: PWAS q-value = 3e-144 (Dominant & recessive)
% Skin cancer
dominant effect scores
recessive effect scores
Bottom 29% 7792/80965
Other samples 13433/193256
Other samples 11754/169752
Top 38% 9471/104469

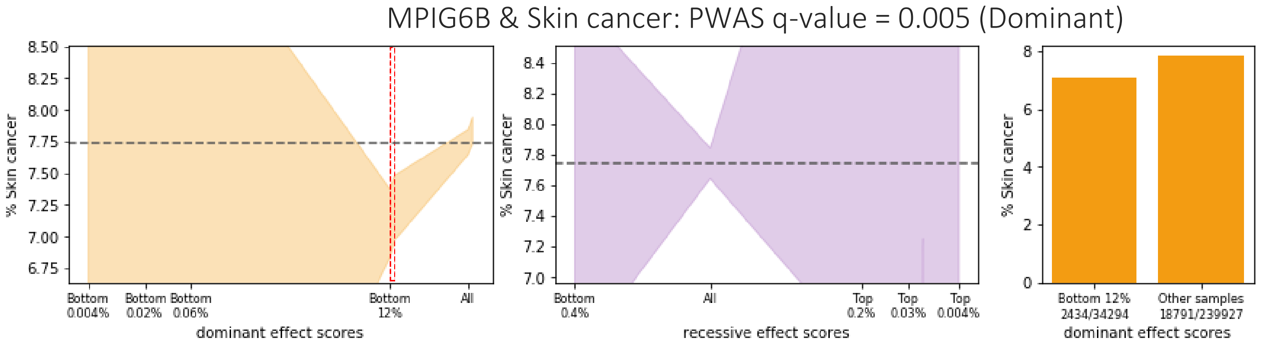
MPIG6B & Skin cancer: PWAS q-value = 0.005 (Dominant)
% Skin cancer
dominant effect scores
recessive effect scores
Bottom 12% 2434/34294
Other samples 18791/239927

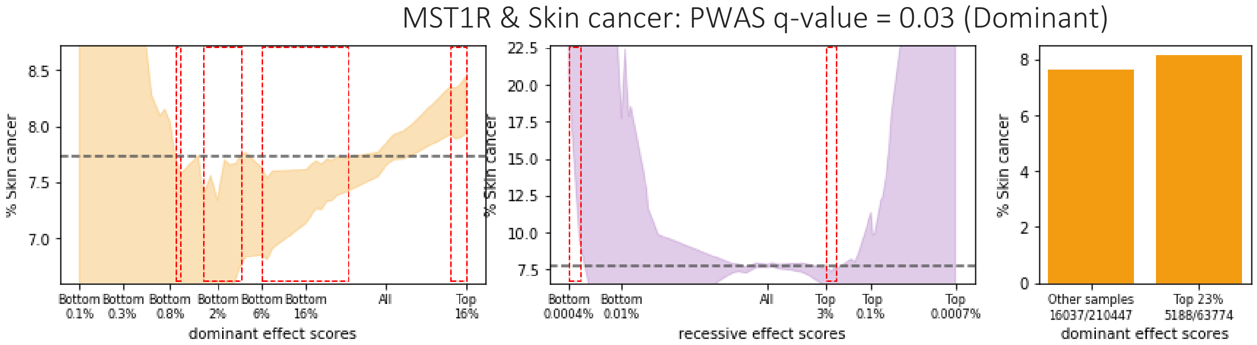
MST1R & Skin cancer: PWAS q-value = 0.03 (Dominant)
% Skin cancer
dominant effect scores
recessive effect scores
Other samples 16037/210447
Top 23% 5188/63774

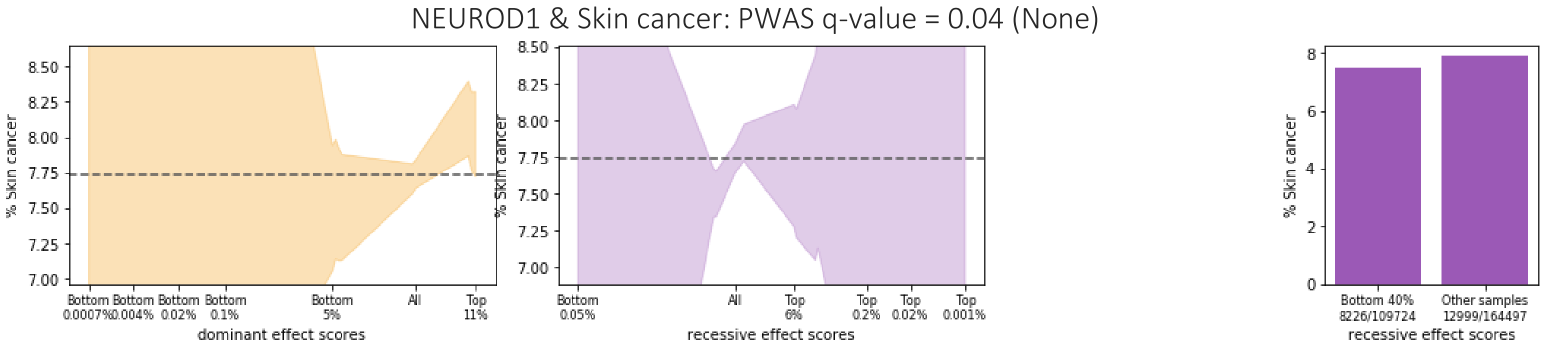
NEUROD1 & Skin cancer: PWAS q-value = 0.04 (None)
% Skin cancer
dominant effect scores
recessive effect scores
Bottom 40% 8226/109724
Other samples 12999/164497

OCA2 & Skin cancer: PWAS q-value = 0.05 (Dominant)

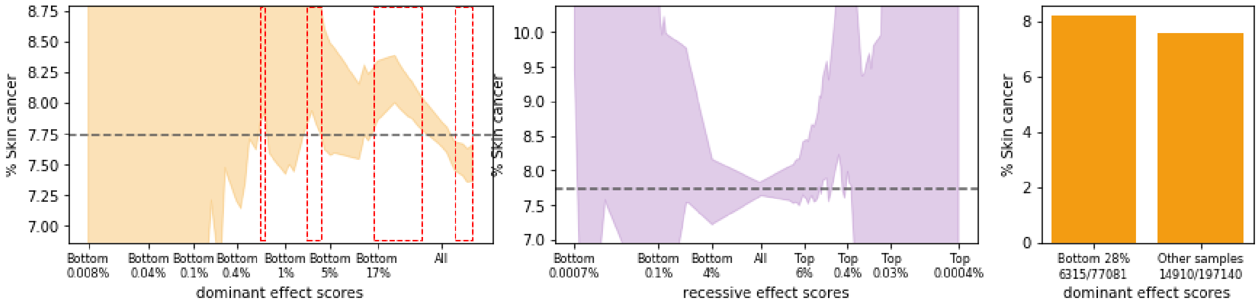

PPP1R18 & Skin cancer: PWAS q-value = 0.008 (Recessive)

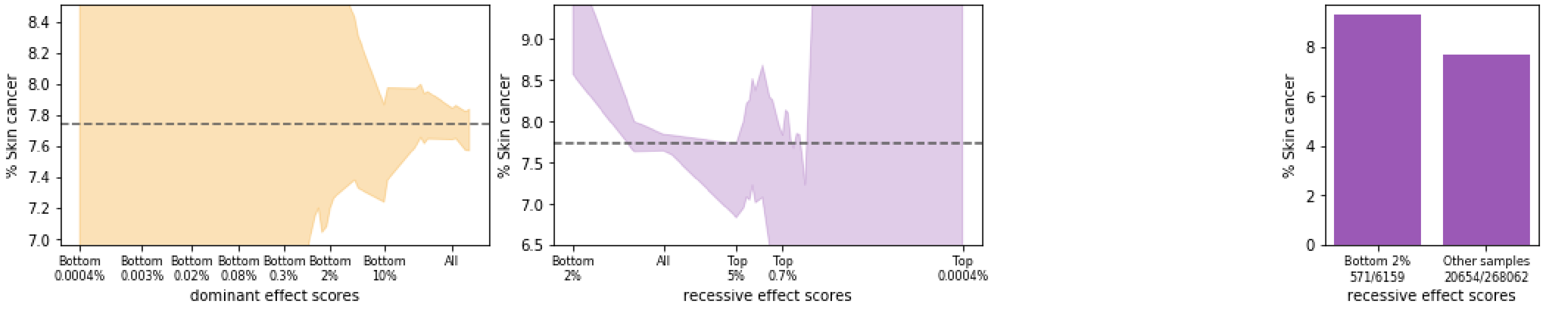

PRDM7 & Skin cancer: PWAS q-value = 2e-08 (Dominant)

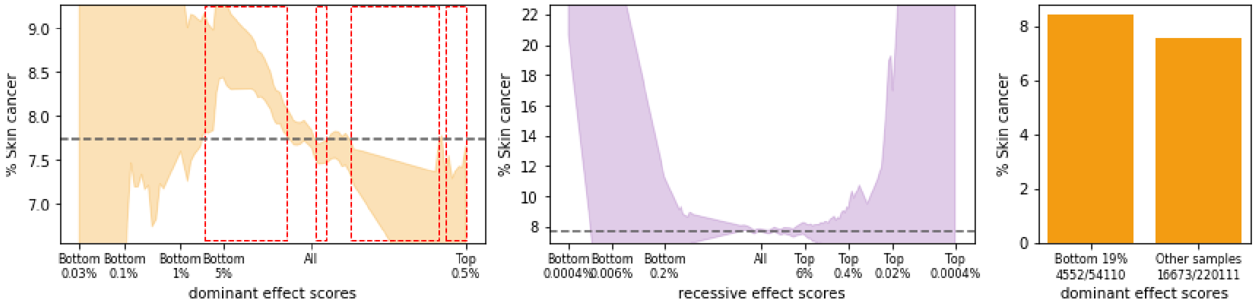

PTPN12 & Skin cancer: PWAS q-value = 0.003 (Recessive)

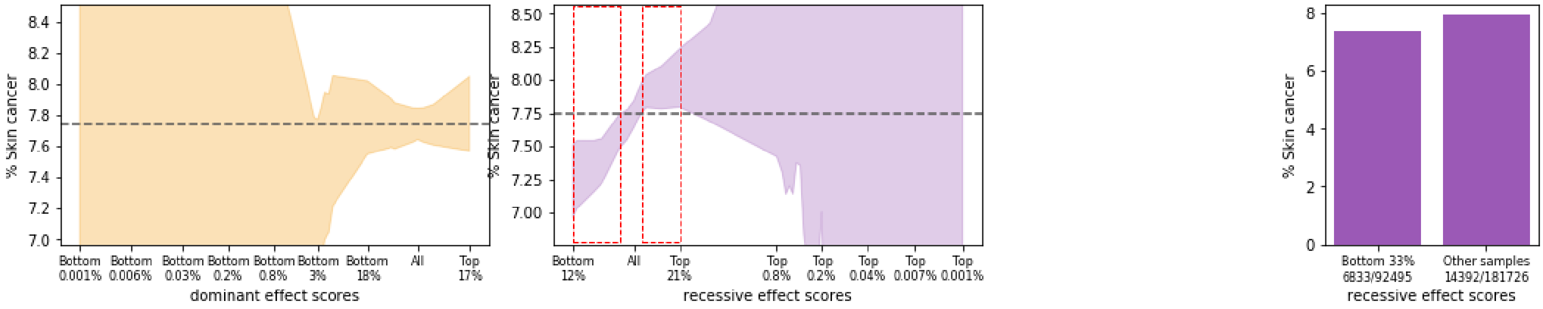

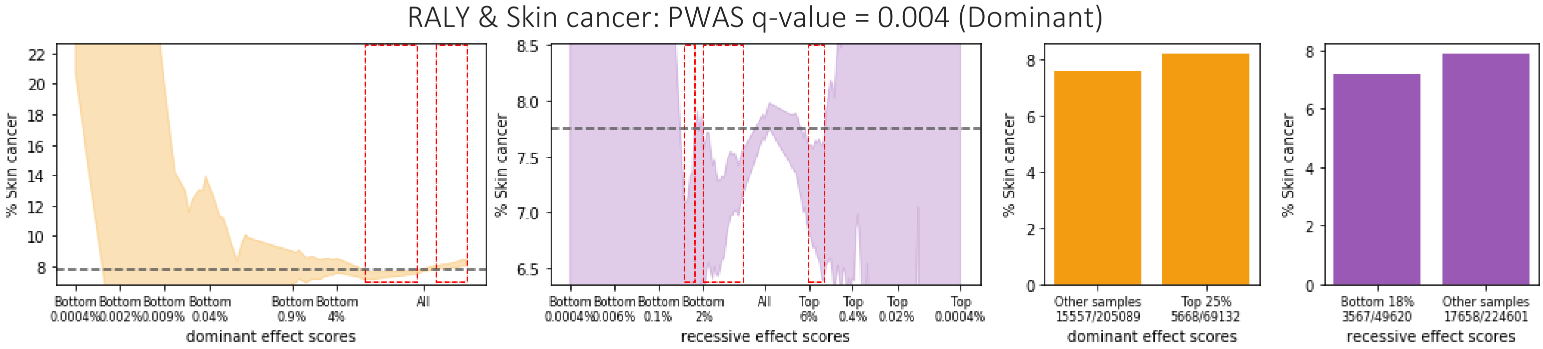
RALY & Skin cancer: PWAS q-value = 0.004 (Dominant)
% Skin cancer
dominant effect scores
recessive effect scores
Bottom 0.0004% Bottom 0.002% Bottom 0.009% Bottom 0.04% Bottom 0.9% Bottom 4% All
Bottom 0.0004% Bottom 0.006% Bottom 0.1% Bottom 2% All Top 6% Top 0.4% Top 0.02% Top 0.0004%
Other samples 15557/205089
Top 25% 5668/69132
Bottom 18% 3567/49620
Other samples 17658/224601

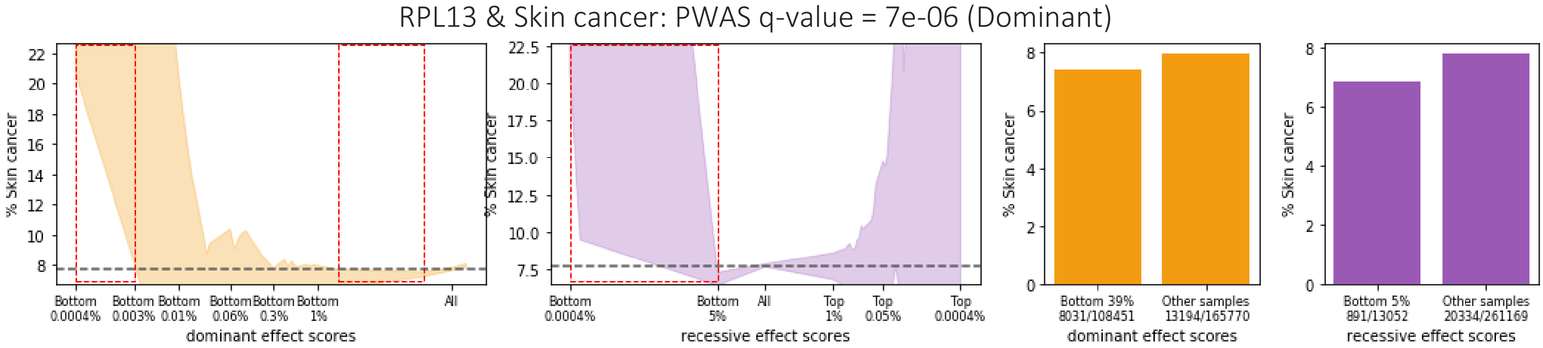
RPL13 & Skin cancer: PWAS q-value = 7e-06 (Dominant)
% Skin cancer
dominant effect scores
recessive effect scores
Bottom 0.0004% Bottom 0.003% Bottom 0.01% Bottom 0.06% Bottom 0.3% Bottom 1% All
Bottom 0.0004% Bottom 5% All Top 1% Top 0.05% Top 0.0004%
Bottom 39% 8031/108451
Other samples 13194/165770
Bottom 5% 891/13052
Other samples 20334/261169

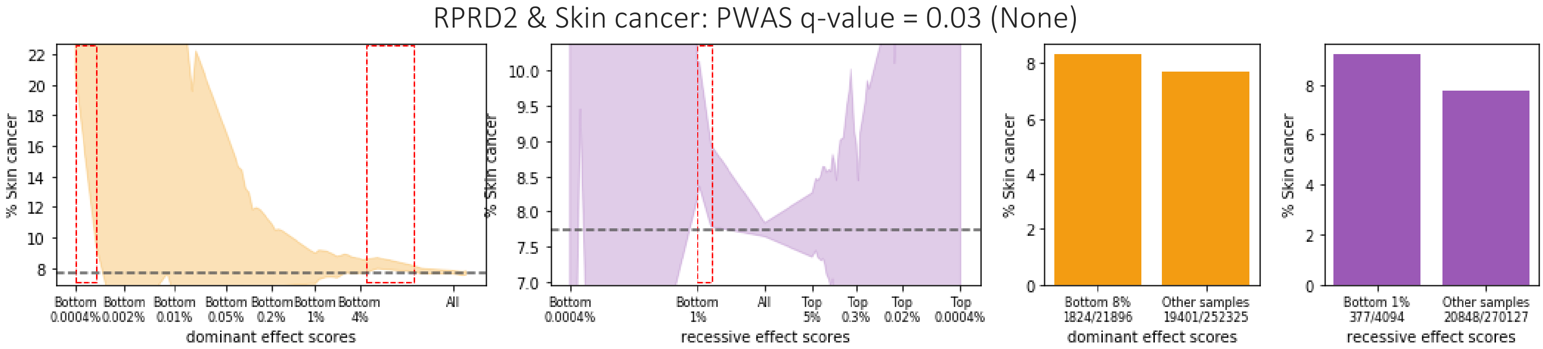
RPRD2 & Skin cancer: PWAS q-value = 0.03 (None)
% Skin cancer
dominant effect scores
recessive effect scores
Bottom 0.0004% Bottom 0.002% Bottom 0.01% Bottom 0.05% Bottom 0.2% Bottom 1% Bottom 4% All
Bottom 0.0004% Bottom 1% All Top 5% Top 0.3% Top 0.02% Top 0.0004%
Bottom 8% 1824/21896
Other samples 19401/252325
Bottom 1% 377/4094
Other samples 20848/270127

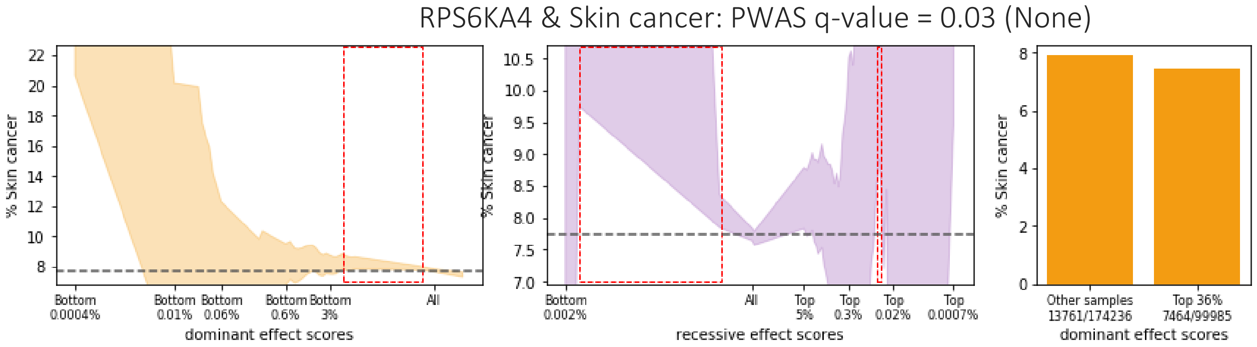
RPS6KA4 & Skin cancer: PWAS q-value = 0.03 (None)
% Skin cancer
dominant effect scores
recessive effect scores
Bottom 0.0004% Bottom 0.01% Bottom 0.06% Bottom 0.6% Bottom 3% All
Bottom 0.002% All Top 5% Top 0.3% Top 0.02% Top 0.0007%
Other samples 13761/174236
Top 36% 7464/99985

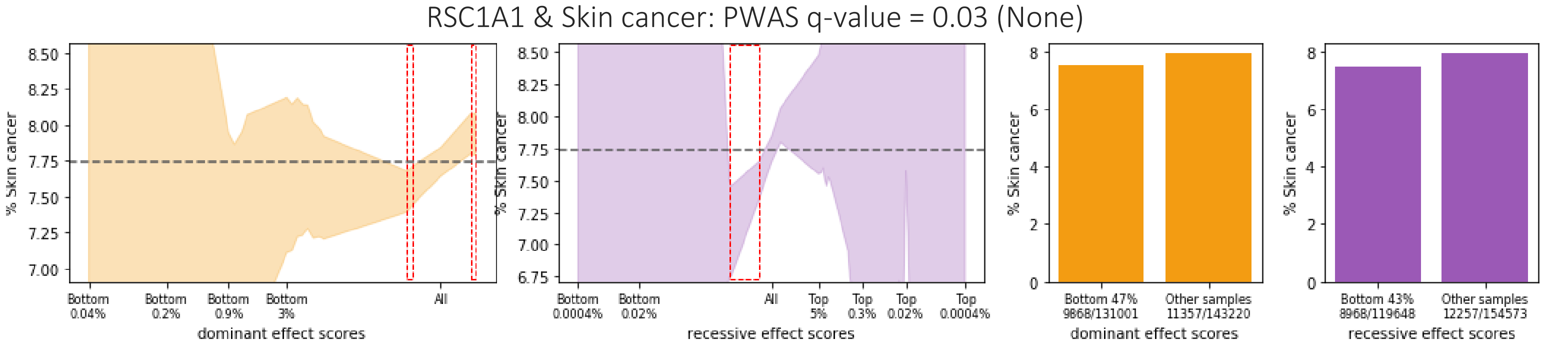
RSC1A1 & Skin cancer: PWAS q-value = 0.03 (None)
% Skin cancer
dominant effect scores
Bottom 0.04%
Bottom 0.2%
Bottom 0.9%
Bottom 3%
All
recessive effect scores
Bottom 0.0004%
Bottom 0.02%
Top 5%
Top 0.3%
Top 0.02%
Top 0.0004%
Bottom 47% 9868/131001
Other samples 11357/143220
Bottom 43% 8968/119648
Other samples 12257/154573

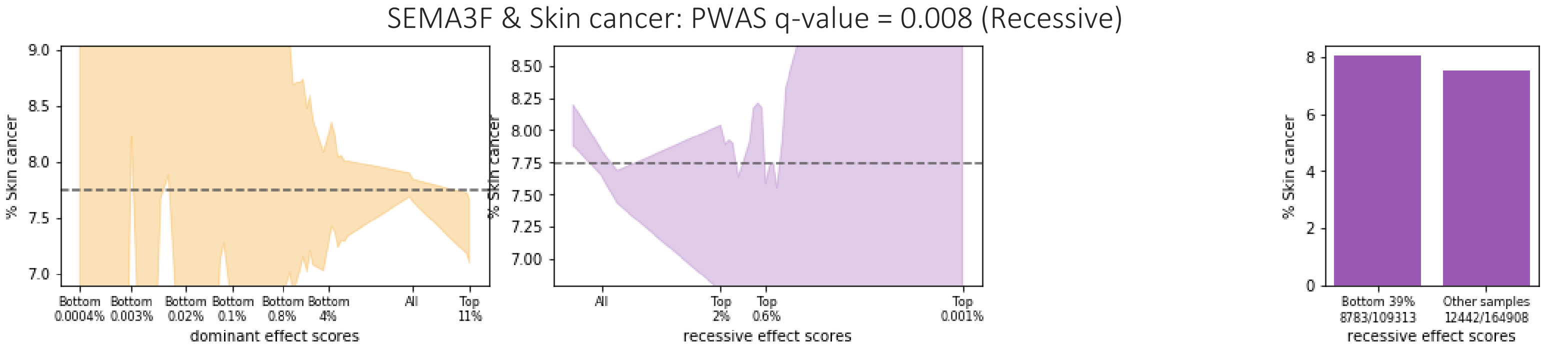
SEMA3F & Skin cancer: PWAS q-value = 0.008 (Recessive)
% Skin cancer
dominant effect scores
Bottom 0.0004%
Bottom 0.003%
Bottom 0.02%
Bottom 0.1%
Bottom 0.8%
Bottom 4%
All
Top 11%
recessive effect scores
Top 2%
Top 0.6%
Top 0.001%
Bottom 39% 8783/109313
Other samples 12442/164908

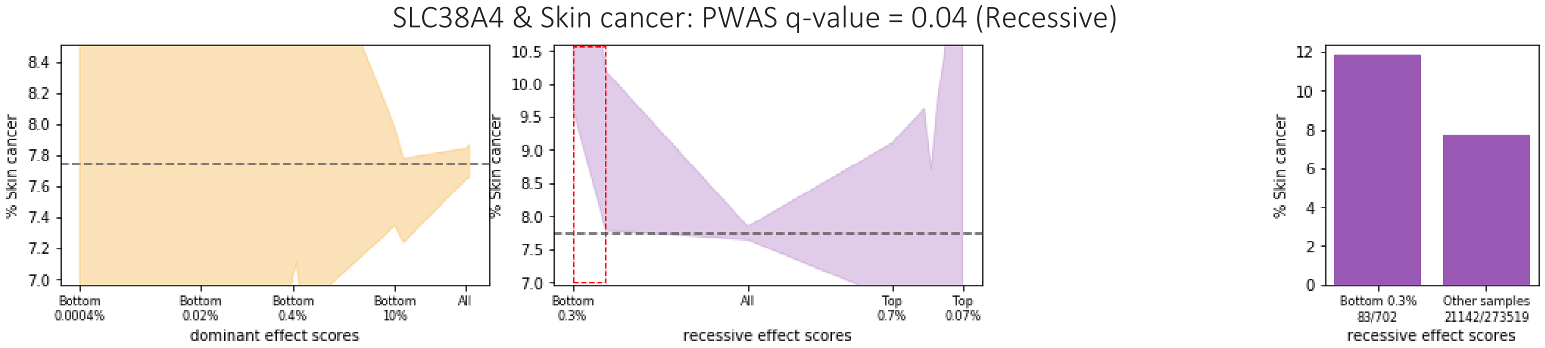
SLC38A4 & Skin cancer: PWAS q-value = 0.04 (Recessive)
% Skin cancer
dominant effect scores
Bottom 0.0004%
Bottom 0.02%
Bottom 0.4%
Bottom 10%
All
recessive effect scores
Bottom 0.3%
Top 0.7%
Top 0.07%
Bottom 0.3% 83/702
Other samples 21142/273519

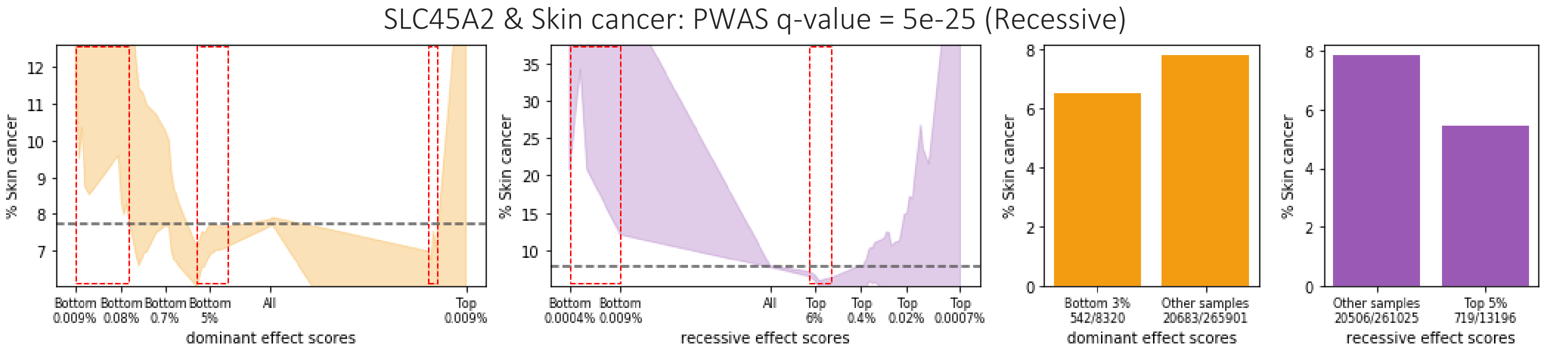
SLC45A2 & Skin cancer: PWAS q-value = 5e-25 (Recessive)
% Skin cancer
dominant effect scores
Bottom 0.009%
Bottom 0.08%
Bottom 0.7%
Bottom 5%
All
Top 0.009%
recessive effect scores
Bottom 0.0004%
Bottom 0.009%
Top 6%
Top 0.4%
Top 0.02%
Top 0.0007%
Bottom 3% 542/8320
Other samples 20683/265901
Other samples 20506/261025
Top 5% 719/13196

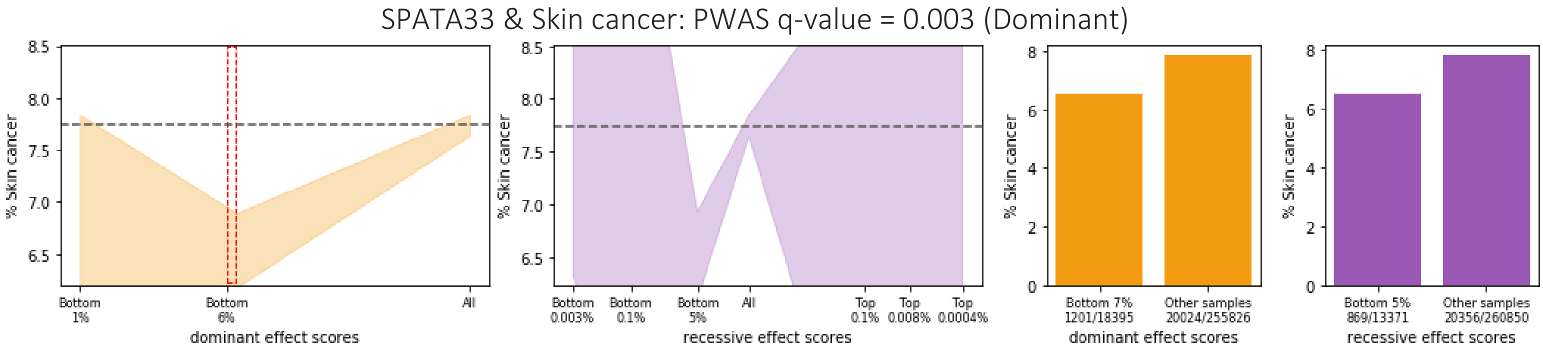
SPATA33 & Skin cancer: PWAS q-value = 0.003 (Dominant)
% Skin cancer
Bottom 1%
Bottom 6%
All
dominant effect scores
Bottom 0.003%
Bottom 0.1%
Bottom 5%
All
Top 0.1%
Top 0.008%
Top 0.0004%
recessive effect scores
Bottom 7% 1201/18395
Other samples 20024/255826
dominant effect scores
Bottom 5% 869/13371
Other samples 20356/260850
recessive effect scores

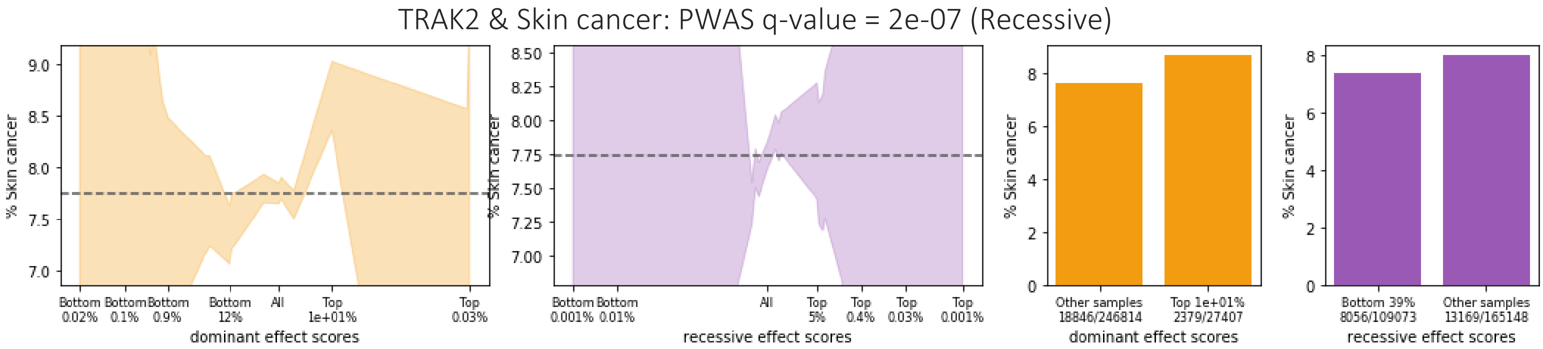
TRAK2 & Skin cancer: PWAS q-value = 2e-07 (Recessive)
% Skin cancer
Bottom 0.02%
Bottom 0.1%
Bottom 0.9%
Bottom 12%
All
Top 1e+01%
Top 0.03%
dominant effect scores
Bottom 0.001%
Bottom 0.01%
All
Top 5%
Top 0.4%
Top 0.03%
Top 0.001%
recessive effect scores
Other samples 18846/246814
Top 1e+01% 2379/27407
dominant effect scores
Bottom 39% 8056/109073
Other samples 13169/165148
recessive effect scores

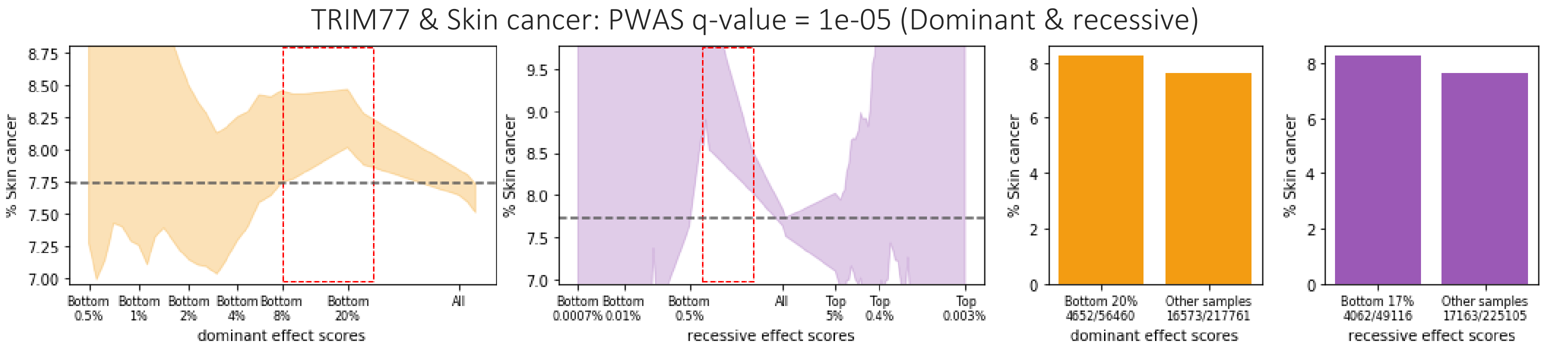
TRIM77 & Skin cancer: PWAS q-value = 1e-05 (Dominant & recessive)
% Skin cancer
Bottom 0.5%
Bottom 1%
Bottom 2%
Bottom 4%
Bottom 8%
Bottom 20%
All
dominant effect scores
Bottom 0.0007%
Bottom 0.01%
Bottom 0.5%
All
Top 5%
Top 0.4%
Top 0.003%
recessive effect scores
Bottom 20% 4652/56460
Other samples 16573/217761
dominant effect scores
Bottom 17% 4062/49116
Other samples 17163/225105
recessive effect scores

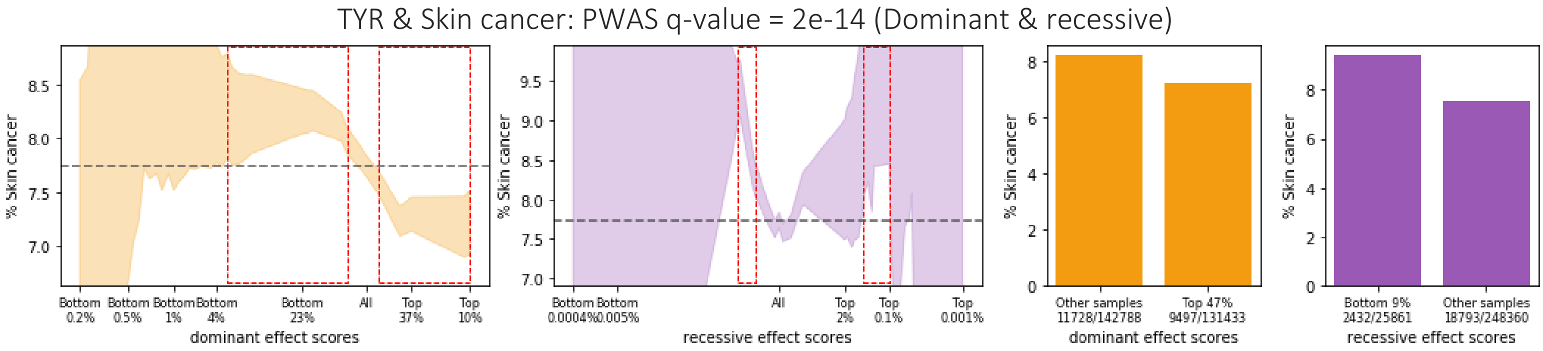
TYR & Skin cancer: PWAS q-value = 2e-14 (Dominant & recessive)
% Skin cancer
Bottom 0.2%
Bottom 0.5%
Bottom 1%
Bottom 4%
Bottom 23%
All
Top 37%
Top 10%
dominant effect scores
Bottom 0.0004%
Bottom 0.005%
All
Top 2%
Top 0.1%
Top 0.001%
recessive effect scores
Other samples 11728/142788
Top 47% 9497/131433
dominant effect scores
Bottom 9% 2432/25861
Other samples 18793/248360
recessive effect scores

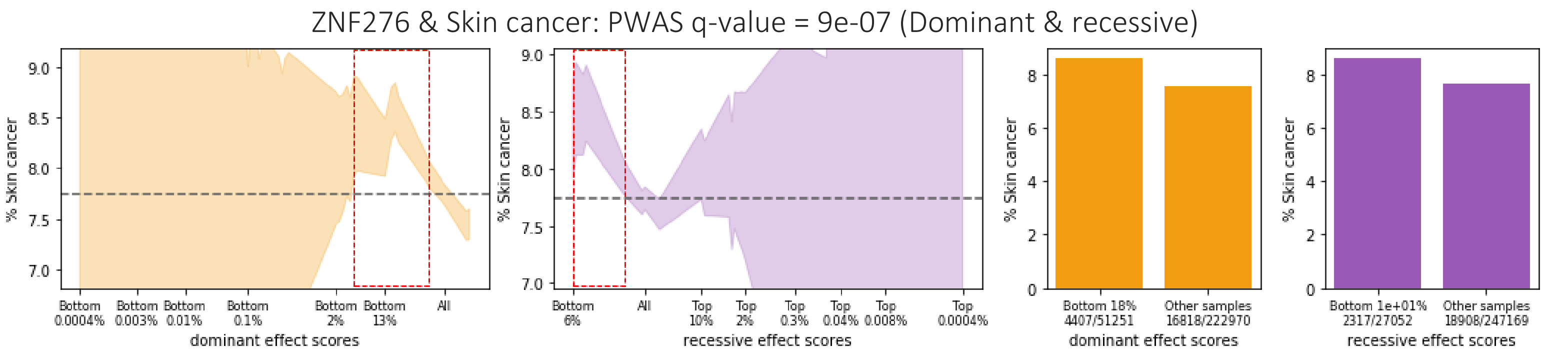
ZNF276 & Skin cancer: PWAS q-value = 9e-07 (Dominant & recessive)
% Skin cancer
dominant effect scores
Bottom 0.0004%
Bottom 0.003%
Bottom 0.01%
Bottom 0.1%
Bottom 2%
Bottom 13%
All
recessive effect scores
Bottom 6%
All
Top 10%
Top 2%
Top 0.3%
Top 0.04%
Top 0.008%
Top 0.0004%
Bottom 18% 4407/51251
Other samples 16818/222970
Bottom 1e+01% 2317/27052
Other samples 18908/247169
